# Description of new mandibular remains of *Microcolobus* from Nakali (ca. 10 Ma, Kenya): implications on the evolution of Miocene colobines

**DOI:** 10.1101/2024.03.31.587461

**Authors:** Laurent Pallas, Masato Nakatsukasa, Yutaka Kunimatsu

**Author notes:** Corresponding author: Dr. Laurent Pallas.

## Abstract

We have described mandibular specimens of fossil colobines from the late Miocene site of Nakali (Kenya). Using qualitative and quantitative dental and mandibular traits, we compared them to an extensive sample of extant colobines and Miocene fossil colobines. We tested the hypothesis that i) only one species was present in this newly described fossil sample, ii) this species is phenetically distinct from fossil colobines from Ngerngerwa and Ngorora, iii) this species is phenetically distinct from hitherto documented fossil Miocene colobines, and iv) this species is phenetically more different from extant African colobines than from Asian colobines. Bootstrap analyses demonstrated that the Nakali specimens belong to a single fossil species. Dental and symphyseal morphometric ratios and a morphometric geometric analysis of the corpus cross-section showed that the Nakali colobines belong to *Microcolobus tugenensis*. Bootstrap analyses failed to unambiguously confirm the distinct taxonomic status of isolated dental specimens from Ngerngerwa and Ngorora, and suggest that they may represent, along with the Nakali sample, a single species. A linear discriminant analysis established on dental linear dimensions of four Nakali specimens classified them within the African colobine tribe (Colobini). *Microcolobus* have a P_4_ occlusal morphology, a breadth differential of the symphyseal transverse tori, and an inclination of the symphysis similar to extant African colobines. However, the corpus shape of *Microcolobus* is closer to that of primitive cercopithecoids, in addition to the lack of the diagnostic mesiodistal elongation of the lower canine of extant African colobines, suggesting that *Microcolobus* is likely a stem Colobinae.

## INTRODUCTION

Extant colobines are small to medium-sized cercopithecid monkeys known from equatorial Africa and eastern Asia (Kingdon and Groves 2013; Rowe and Jacobs 2016). They originated in Africa, with a first appearance datum documented by isolated dental elements from late Miocene deposits in eastern Africa (Rossie et al. 2013). Cercopithecidae are sister taxa to the Victoriapithecidae (Szalay and Delson 1979, but see Benefit and Pickford 1986), an ancestral stock of small omnivorous and partly terrestrial monkeys documented in eastern and northern Africa (Benefit 1993, 1999, 2008; Miller et al. 2009; Locke et al. 2020). In contrast to the African Plio-Pleistocene fossil record, the early phases of the colobine evolution in the late Miocene are poorly known (Delson 1973; Benefit and Pickford 1986; De Bonis et al. 1990; Koufos et al. 2003; Hlusko 2006, 2007; Frost et al. 2009; Gilbert et al. 2010; Suwa et al. 2015; Pallas et al. 2019; Gommery et al. 2022). Hitherto, our knowledge of their masticatory adaptations has come from isolated dental elements from various fossil deposits in eastern and northern Africa, relatively complete mandibular, cranial and postcranial elements from Toros-Ménalla (Chad), partial skulls from Lukeino (Kenya), a partial skull from Wadi Natrun (Egypt), and a partial mandible from Ngerngerwa (or Ngeringerowa, Kenya).

Recent fossil discoveries in the late Miocene deposits of the Nakali Formation (Kenya) have greatly contributed to clarifying the functional adaptations and phylogeny of the earliest colobines through the description of two partial skeletons and recovery of associated and isolated dental, cranial and mandibular elements (Nakatsukasa et al. 2010). The partial skeletons KNM-NA 48916 and KNM-NA 48915 were tentatively allocated to *Microcolobus*. It is a small colobine (ca. 5 kg as inferred from its dental dimensions) originally described from the neighbouring site of Ngerngerwa, Tugen Hills. The taxonomic attribution of the Nakali skeletons to *Microcolobus* is based on the association of one molar (KNM-NA 48915A) with the partial skeleton KNM-NA 48915. The marked phenetic similarity between the two skeletons is the main rationale behind the association of the second partial skeleton (KNM-NA 48916) to *Microcolobus*. The postcranial anatomy of the Nakali *Microcolobus* depicts a colobine functionally adapted to the use of arboreal substrates, with a long pollex (thumb) that contrasts with the reduced one observed in extant African colobines (Colobini). The relatively long thumb of the Nakali *Microcolobus* is similar to that of the Eurasian early fossil colobine *Mesopithecus*, of which hundreds of fossil remains have been published, including fairly complete mandibular and dental elements from the late Miocene site of Pikermi, Greece (Delson 1973; De Bonis et al. 1990; Koufos et al. 2003).

Given the lack of published mandibular and cranial material from Nakali and the absence of a thorough functional analysis of the dental material associated with the partial skeleton KNM-NA 48915, the taxonomy of the Nakali colobine remains uncertain. The partial *Mi. tugenensis* mandible KNM-BN 1740, which notably preserves a complete toothrow, a preserved corpus outline at the M_1_/M_2_ level, and a partially preserved symphysis, is hitherto our best line of evidence on the mandibular and dental anatomy of the earliest African colobines (Benefit and Pickford 1986). A previous morphometric analysis of an isolated molar from Nakali (KNM-NA 305) by Benefit and Pickford (1986) concluded that this molar was distinct from that of KNM-BN 1740 (*Mi. tugenensis*). An isolated P_4_ KNM-BN 1251 from the nearby Ngorora site in the Tugen Hills also provided evidence, based partly on relative crown dimensions, for taxic diversity in the ca. 10 Ma colobine fossil record from soutwestern Kenya (Benefit and Pickford 1986; Rossie et al. 2013). An isolated lower molar (CHO-BT 78) from the younger site of Beticha (ca. 8 Ma, Ethiopia) was also likened to *Microcolobus* (Suwa et al. 2015), but given the small number of specimens from both Beticha and Ngerngerwa, no definitive conclusion has been made. The colobine Miocene fossil record is complemented, for the 7 – 6 Ma period, by partial mandibles from the taxa *Cercopithecoides bruneti*, *Paracolobus enkorikae*, and *Sawecolobus lukeinoensis* (Hlusko 2007; Pallas et al. 2019; Gommery et al. 2022). Specimens belonging to these taxa include partially preserved symphyses, corpus and toothrows.

Dental, symphyseal, and corpus anatomy of fossil and extant colobines proved useful for taxonomic discrimination. Precisely, the relative length of the lower canine (Pan 2006; Suwa et al. 2015), the relative proportions and position of the P_4_ cusps (Swindler and Orlosky 1974; Szalay and Delson 1979), the enlargement of the lower premolars (Pan 2006), the relative width and length of the lower molars (Delson 1973; Pan 2006; Gilbert et al. 2010), and notably the relative width of the M_3_ lophids, the development of the M_3_ hypoconulids (Delson 1973; Willis and Swindler 2004; Takai et al. 2015, 2016), the angulation of the symphysis (Pallas et al. 2019), the length of the planum alveolare (Pallas et al. 2019), the breadth proportions of the symphyseal transverse tori (Benefit and Pickford 1986; Pallas et al. 2019), the development of the digastric and submandibular fossae (Wright et al. 2008), and the development of the lateral prominence of the mandibular corpus (Benefit and Pickford 1986) were emphasized in taxonomic distinction and included in the diagnosis of fossil and extant colobines.

Here, we described six mandibular remains from the Nakali Formation, which includes specimens with well-preserved dental, symphyseal and corpus anatomy. We take advantage of the previously mentionned dental and mandibular diagnostic features to evaluate the following hypotheses on the Nakali colobine sample : H0_1_ The Nakali mandibular sample described here are consistent with a single colobine species; H0_2_ The newly discovered Nakali specimens are distinct from the penecontemporaneous colobines of Ngerngerwa and Ngorora, as previously suggested by Benefit and Pickford (1986); H0_3_ The newly discovered Nakali specimens are distinct from the Eurasian and African Miocene colobines documented to date; H0_4_ The newly discovered Nakali specimens are no more closely related to extant African colobines (Colobini) than to extant Asian colobines (Presbytini), as hypothesized by Nakatsukasa et al. (2010) and Benefit and Pickford (1986) on the basis of postcranial and dentognathic data, respectively. We also provided provisional sex assignment to selected specimens using the absolute and relative proportions of the dimorphic C_1_ and P_3_. Using univariate and multivariate morphometry, we investigated shape differences in mandibular and dental traits within the Nakali sample, and between the Nakali sample and available penecontemporaneous fossil colobines from the Eurasian and African Miocene fossil record. We reported data regarding the morphological and taxonomical homogenenity of the Nakali sample through bootstrap analyses and linear discriminant analyses. This study aims to clarify the taxonomic status and paleobiology of fossil colobines from Nakali, to clarify the taxonomy of isolated dental specimens penecontemporaneous to Nakali, and to provide new evidence on the date of appearance of diagnostic anatomical traits of extant colobines.

## MATERIAL AND METHODS

### COMPARATIVE EXTANT DATASET

#### Dental

We measured dental dimensions on male and female specimens of *Colobus guereza* (*n* = 20), *Piliocolobus badius* (*n* = 20), *Procolobus verus* (*n* = 20), *Trachypithecus cristatus* (*n* = 20), *Presbytis chrysomelas* (*n* = 18), and *Nasalis larvatus* (*n* = 20) specimens. Information on specimen accession numbers and the sex of the specimens used for the calculation of morphometric ratios and multivariate analysis can be found in Online Resource 1.

We evaluated qualitative dental traits on *Colobus guereza* (*n* = 20), *Colobus polykomos* (*n* = 39), *Piliocolobus badius* (*n* = 38), *Procolobus verus* (*n* = 39), *Trachypithecus* spp. (*n* = 27), *Presbytis* spp. (*n* = 35), and *Nasalis larvatus* (*n* = 20) specimens. Information on specimen accession numbers and sex can be found in Online Resource 2.

#### Mandibular

We measured symphyseal dimensions on *Colobus* (*n* = 58), *Piliocolobus* (*n* = 38), *Procolobus* (*n* = 23), *Nasalis* (*n* = 29), *Trachypithecus* (*n* = 27), *Presbytis* (*n* = 70), *Simias* (*n* = 25), *Semnopithecus* (*n* = 14), *Pygathrix* (*n* = 5), and *Rhinopithecus* (*n* = 2). See Online Resource 3 for information on the taxonomy, accession number, and sex of these specimens.

We digitized landmarks and semi-landmarks on the corpus of *Colobus* (*n* = 111), *Piliocolobus* (*n* = 42), *Procolobus* (*n* = 26), *Nasalis* (*n* = 49), *Trachypithecus* (*n* = 18), *Presbytis* (*n* = 69), *Simias* (*n* = 25), and *Semnopithecus* (*n* = 15). Information on accession numbers and specimen sex can be found in Online Resource 4.

### COMPARATIVE FOSSIL DATASET

#### Dental

We measured and compiled dental dimensions of 133 specimens of Mio-Pliocene colobine taxa and victoriapithecids. This includes the dental dimensions of specimens of *Mi. tugenensis*, *Ce. bruneti, Pa. enkorikae*, *Kuseracolobus aramisi, Me. pentelicus, Me. delsoni*, *S. lukeinoensis*, and *Victoriapithecus macinnesi*. Fossil colobine specimens of unknown generic attribution were also included as they may represent *Microcolobus* specimens. It comprises taxonomically indeterminate specimens from a site complex of the Ngorora and Lukeino Formation in the Tugen Hills region (including Ngerngerwa and the Mpesida Beds), the Beticha locality (Chorora Formation), the Lemudong’o Formation, the Lothagam region (Nawata Formation), the Middle Awash region (Sagantole and Adu-Asa formations), and the locality of Menacer. Detailed information on specimen accession numbers and source of the source data can be found in Online Resource 5.

#### Mandibular

We measured symphyseal dimensions on *V. macinnesi* (KNM-MB 18993), *Noropithecus bulukensis* (KNM-WS 123), *Mi. tugenensis* (KNM-BN 1740), *Pa. enkorikae* (KNM-NK 44770, KNM-NK 45912, KNM-NK 42276, and KNM-NK 36515), *Me. delsoni* (RZO 325), and *Me. pentelicus* (MNHN PIK 430 and MNHN PIK 428). We digitized landmarks and semi-landmarks on the corpus of *V. macinnesi* (KNM-MB 1, KNM-MB 18993, and KNM-MB 27876), *N*. *bulukensis* (KNM-WS 12638), *Me. pentelicus* (MNHN PIK 002, MNHN PIK 007, MNHN PIK 034, MNHN PIK 006, MNHN PIK 019B, BSM AS-II 15), *Pa. enkorikae* (KNM-NK 44770), and *Mi. tugenensis* (KNM-BN 1740).

### DENTAL MORPHOMETRY

We measured the maximum mesiodistal and buccolingual dimensions of extant and fossil colobines teeth using a Mitutoyo Digital Caliper to the nearest 0.1 mm according to the protocol illustrated in Figure 1. The assessment of the occlusal relief of KNM-NA 51103 (NH and NR variables) was done using the measurement method of Benefit and Pickford (1986) from photographs of the 3D generated surface. NH and NR dimensions were measured using ImageJ (Schneider et al. 2012). The dental dimensions of the Nakali fossil colobines described here and those of the extant colobines included in the comparative dataset were acquired by L.P.

**Figure 1:**
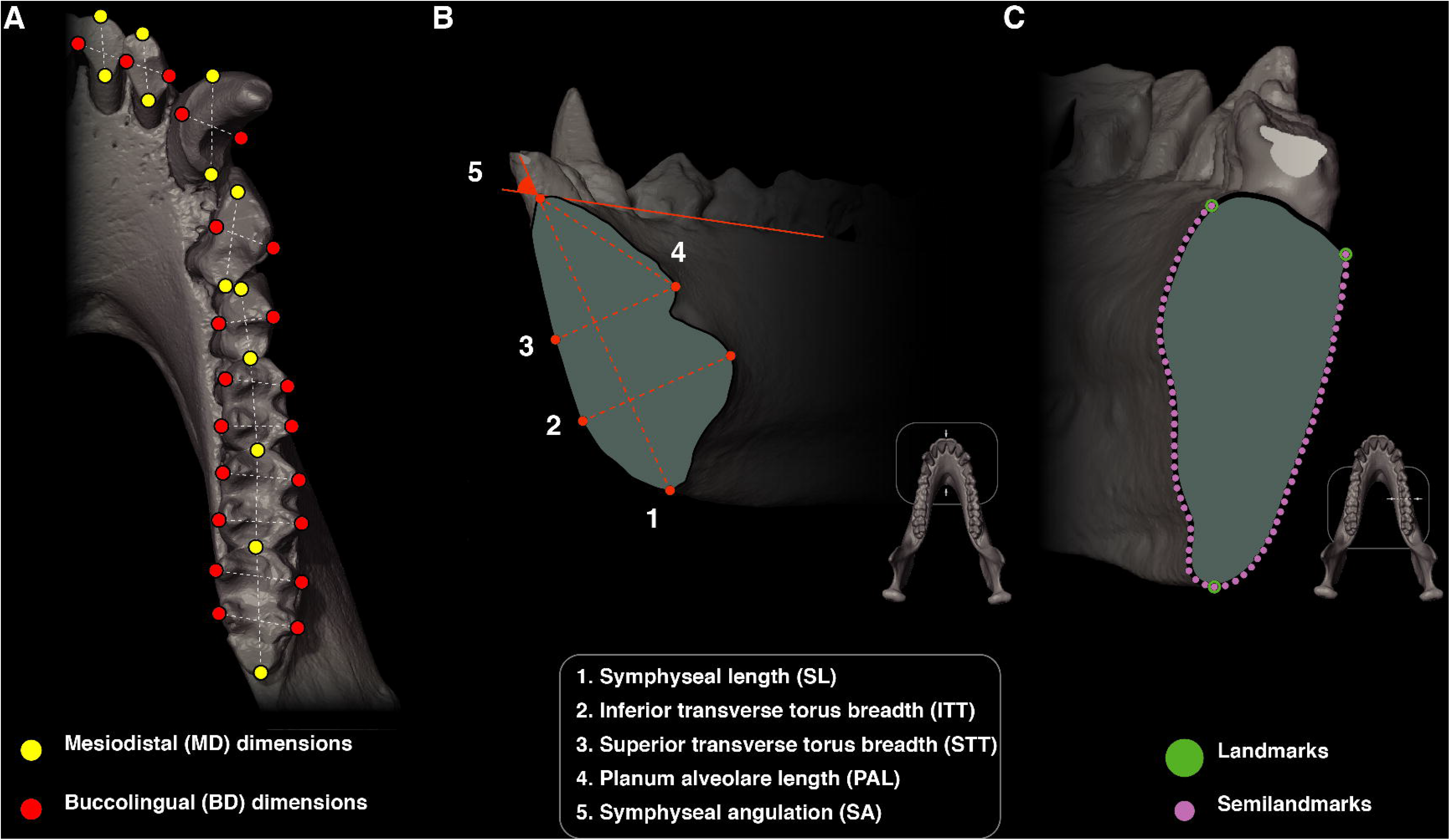
Illustrations of A) the dental morphometric protocol, B) the symphyseal morphometric protocol, and C) the positioning of the *n* = 75 semilandmarks and *n* = 3 landmarks on the cross-section of the corpus at M_1_-M_2_ junction.

Raw dental morphometric data for the Nakali specimens are provided in Online Resource 6. Descriptive statistics of the raw dental morphometric data for the extant colobine sample are provided in Online Resource 7 and descriptive statistic of the dental indices of the extant colobine sample can be found in Online Resource 8.

### MANDIBULAR MORPHOMETRY

We took four linear and one angular measurements on the symphysis sagittal cross-section of extant and fossil colobines. The protocol is illustrated in Figure 1. Measurements were acquired using ImageJ v.1.50e (Schneider et al. 2012) by L.P. Symphyseal measurements from KNM-BN 1740 were not included in the analysis as its inferior transverse torus is damaged and most of it is missing according by our computed tomographic estimation (Online Resource 9). Raw data of the fossil sample for the symphyseal measurements (including Nakali) is provided in Online Resource 10.

We digitized landmarks (*n* = 3) and semi-landmarks (*n* = 75) on the corpus at M_1_-M_2_ junction among extant and fossil colobines using the protocol presented in Pallas et al. (submitted) and illustrated in Figure 1. Landmarks and semi-landmarks were digitized using tpsDig v.2.32 by L.P.

### 3D SCANS

3D data for the Nakali fossil colobine specimens illustrated here were obtained using an EinScan Pro 2X (Shining 3D, Huangzhou) structured light 3D scanner. The data were then processed using Geomagic Wrap (3D Systems, Rockhill) and photographs of the reconstructed models acquired with Avizo v.7.0 (Thermo Fisher Scientific, Waltham).

### DENTAL AND SYMPHYSEAL MORPHOMETRIC RATIO

To identify differences between extant colobines and isolated fossil specimens that retain too little data to be included in multivariate analyses, we calculated 7 dental ratios. These ratios quantify the relative width (buccolingual: BL) of the molar lophids, the area of the P_4_ relative to that of M_1_, the length of the M_1_ relative to the breadth of its mesial lophid, the relative length (mesiodistal: MD) of the M_1_ relative to that of the M_2_, the shape of the P_4_ crown and the length of the C_1_ relative to the geometric mean (GM) of the M_1_ measurements (one MD and two BL dimensions across the lophs). When available, measurements of the right teeth were used to compute the indices in the Nakali specimens.

We assessed differences in symphyseal shape between taxa using two indices, the breadth differential of the symphyseal transverse tori and the length of the planum alveolare relative to the GM of four symphyseal measurements. Formulas of the indices are indicated in Table 1.

**Table 1:** Name, formula and abbreviation of the morphological indices used in the study.

| Index | Formula | Abbreviations |
| --- | --- | --- |
| Length of the C<sub>1</sub> relative to GM of M<sub>2</sub> | $C_1 \text{ MD} / M_1 \text{ GM}$ | C<sub>1</sub> Rel. |
| P<sub>3</sub> shape | $P_3 \text{ MD} / P_3 \text{ BL}$ | P<sub>3</sub> shape |
| P<sub>4</sub> shape | $P_4 \text{ BL} / P_4 \text{ MD}$ | P<sub>4</sub> shape |
| Area of the P<sub>4</sub> relative to area of the M<sub>1</sub> | $[(P_4 \text{ MD} * P_4 \text{ BL}) / (M_1 \text{ MD} * M_1 \text{ DBL})]$ | P<sub>4</sub> ar. M<sub>1</sub> ar. |
| M<sub>1</sub> length relative to M<sub>2</sub> length | $M_1 \text{ MD} / M_2 \text{ MD}$ | M<sub>1</sub> lg. M<sub>2</sub> lg. |
| Breadth differential of the M<sub>1</sub> lophids | $M_1 \text{ MBL} / M_1 \text{ DBL}$ | M<sub>1</sub> loph. diff. |
| Breadth differential of the M<sub>2</sub> lophids | $M_2 \text{ MBL} / M_2 \text{ DBL}$ | M<sub>2</sub> loph. diff. |
| Breadth differential of the M<sub>3</sub> lophids | $M_3 \text{ MBL} / M_3 \text{ DBL}$ | M<sub>3</sub> loph. diff. |
| Breadth differential of the symphyseal transverse tori | STT / ITT |  |
| Relative length of the planum alveolare | PL / GM |  |

### DENTAL QUALITATIVE TRAITS

In addition to quantitative dental traits, we evaluated differences between extant and fossil colobines using qualitative traits. Several qualitative dental traits were used to distinguish between extant colobine subfamilies and genera. An illustration of those dental traits, for the extant colobines considered in this study, is provided in Figure 2. Numbers followed by the letter ‘a’ illustrate the Colobini condition while numbers followed by the letter ‘b’ illustrate the condition observed in Presbytini.

**Figure 2:**
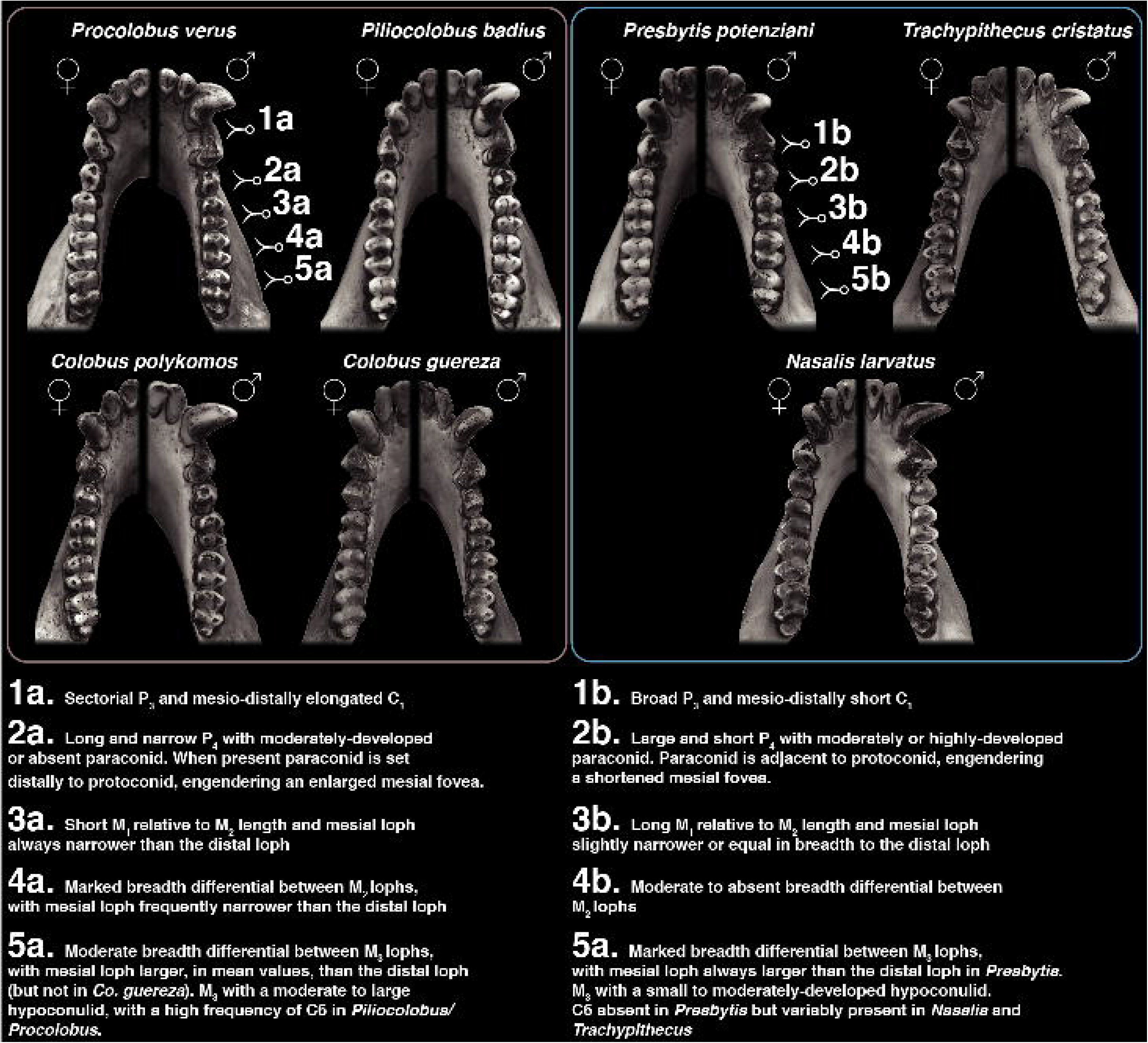
Dental qualitative traits considered in this study, illustrating the differences between African (left column) and Asian (right column) colobines.

We also evaluated the percentage frequency of some of these qualitative dental features. From photographs, we scored the position of the P_4_ metaconid, the position of the M_3_ hypoconulid and the absence / presence of a tuberculum sextum (C6) on the M_3_. The P_4_ metaconid is either adjacent or distal to the protoconid. The M_3_ hypoconulid is positioned either buccally or centrally to the long axis of the anterior portion of the teeth (anterior to the distal lophid).

### STATISTICAL ANALYSES

#### Multivariate dental analysis (Principal Component Analysis and Linear Discriminant Analysis)

We ordered and explored variation in dental dimensions using Principal Component Analysis (PCA). We ran two PCA, one with a ‘sparse dataset’ that comprised 10 variables to include a maximum number of fossil specimens, and a second one, that comprised 13 variables and included only the best-preserved specimens. Specifically, we used the following 13 dental variables for the ‘complete dataset’: C_1_ MD, C_1_ BL, P_3_ MD, P_3_ BL, P_4_ MD, P_4_ BL, M_1_ ML, M_1_ MBL, M_1_ DBL, M_2_ ML, M_2_ MBL, M_2_ DBL, M_3_ MBL, and M_3_ DBL (MBL: mesial BL, DBL: distal BL). We used the following 10 dental variables for the sparse dataset: P_3_ MD, P_3_ BL, P_4_ MD, P_4_ BL, M_1_ ML, M_1_ MBL, M_1_ DBL, M_2_ ML, M_2_ MBL, M_2_ DBL, and M_3_ MBL. First, in each PCA, we limited the effect of size differences on the dental variables using the log-shape ratio method (Mosimann 1970). PCA, on scaled and centered data, was then computed using the function prcomp() of the ‘stats’ package.

To test the hypothesis of a close phyletic relationship between *Microcolobus* and extant Colobini, we computed a Linear Discriminant Analysis (LDA) based on the first four PCs obtainted from each of the two PCAs. Those PCs account for 86% of the variance (PC1 = 31.88%, PC2 = 21.95%; PC3 = 21.99%; PC4 = 10.30%) for the sparse dataset and 78% of the variance for the complete dataset (PC1 = 37.55%, PC2 = 22.96%; PC3 = 9.32%; PC4 = 7.73%). For each LDA, prior to including fossil specimens, we tested the efficiency of the model by splitting the dataset into a training and a test set, with 60% of the specimens randomly selected to be part of the training sample and 40% used to assess the classification rate of the model. The LDA model established on the training set of the sparse PCA correctly classified the subfamily of 89.6% of the specimens from the test sample, hence validating the use of the LDA model to classify four of the Nakali fossil specimens (KNM-NA 48912, KNM-NA 48913, KNM-NA 51102, and KNM-NA 54785). The LDA model computed using the complete dataset correctly classified the subfamily of 89.4% of the specimens from the test sample.

#### Multivariate corpus analysis

We used a PCA to order and explore variations in corpus shape. A detailed description of the statistical methods underlying our multivariate corpus analysis is provided in Pallas et al. (submitted).

#### Statistical tests on dental morphometric ratio and PCA scores

As required before using parametric tests, we evaluated the normality and homoscedasticity of the model’s residuals. Normality was evaluated with a Shapiro-Wilk test using the shapiro.test() function, and homoscedasticity was assessed with a Bartlett test using the bartlett.test() function in the ‘stats’ package. When homoscedasticity of the residuals was not demonstrated, we used non-parametric tests. Results of these tests are found in Online Resource 11 and we have set the level of significance of these tests at 5%.

To assess the presence of significant differences in dental morphometric ratio between extant colobines, we used the parametric Analysis of Variance (ANOVA) test and the non-parametric Kruskall-Wallis test. To identify pairs of significantly distinct genera, we used parametric (Tukey Honest Significant Difference) and non-parametric (Dunn’s Kruskal-Wallis Multiple Comparisons) post-hoc tests. Tukey Honest Significant Difference (HSD) was calculated using the TukeyHSD() of the ‘stats’ package, and Dunn’s test using the dunnTest() function of the ‘FSA’ package. We have set the level of significance of these tests at 5%. We evaluated, using these tests, whether extant colobine subfamilies and genera differed significantly in values of dental morphometric ratio.

#### Coefficient of variation and bootstrap analysis

We evaluated whether one or more species is present in the Nakali fossil sample described here. We also assessed whether other penecontemporaneous fossil colobines from Ngorora and Ngerngerwa can be accomodated within the range of variation of the Nakali sample. To this end, we compared the range of variation of the Nakali sample and the Nakali sample combined with other fossil colobines from Ngerngerwa and Ngorora with that of extant colobine species. First, we used the coefficient of variation (CV; standard deviation / mean) to assess whether a considered fossil sample exceeds the variation of that of an extant colobine species for each dental morphometric ratio. Second, considering that the CV is influenced by the sample size, we designed a bootstrap analysis, with a protocol following of Plavcan and Cope (2001:216) to evaluate the probability that a fossil sample exceeds the variation of a sample of extant colobine species of equal size. Such a protocol was also used in other paleontological studies dealing with cercopithecoids and has proven to be useful (Miller et al. 2009; Locke et al. 2020). For each extant colobine species, we generated 10,000 distributions using the descriptive statistics (mean and standard deviation) observed in our dental morphometric ratio dataset (Online Resource 8). To generate the theoretical distributions in R, we used the replicate() function. We designed a loop to sample, with replacement, for each colobine species, a number of specimens equal to that of the fossil sample considered. For example, we calculated the ‘M1 lophid. diff. ratio’ on *n* = 6 Nakali specimens and on *n* = 4 specimens for ‘M3 lophid. diff.’ ratio. Therefore, we randomly sampled *n* = 6 specimens from each of the 10,000 distributions generated for each of the colobine species to evaluate whether the Nakali sample, in ‘M1 lophid. diff. ratio’, is more variable than that of extant colobine species. To compare the variability between samples, we calculated the CV for each of the 10,000 generated distributions from *n* = 6 specimens of extant colobines. To test if the CV of the fossil sample is significantly exceeding that of extant colobine species, we followed the Cope and Lacy test, and calculated the percentage of CV, per each extant colobine species distribution, that is exceeding that of the fossil sample. In cases where percentage was <5%, we rejected the null hypothesis of a single species in the fossil sample.

## RESULTS

### DESCRIPTION

#### KNM-NA 51102 (mandible with a subcomplete tooth row)

KNM-NA 51102 is a 45.9 mm long mandibular fragment preserving most of the right hemimandible and a ca. 10.0 mm long portion of the left hemimandible (Figure 3). The specimen preserves a partial tooth row with P_3_-P_4_ and M_1_-M_3_. The corpus is deformed and fractured in several aspects. A large fracture that runs from the M_2_ / M_3_ junction to the inferior margin of the corpus, roughly at the level of P_4_ / M_1_, resulted in a ca. 3.0 mm long and a ca. 6.0 mm high loss of bone on the inferior margin of the corpus. The posterior part of the mandibular fragment is also slightly dislocated medially due to this fracture. The alveolar margin is slightly abraded on the post-incisor tooth row but markedly abraded on the incisor row.

**Figure 3:**
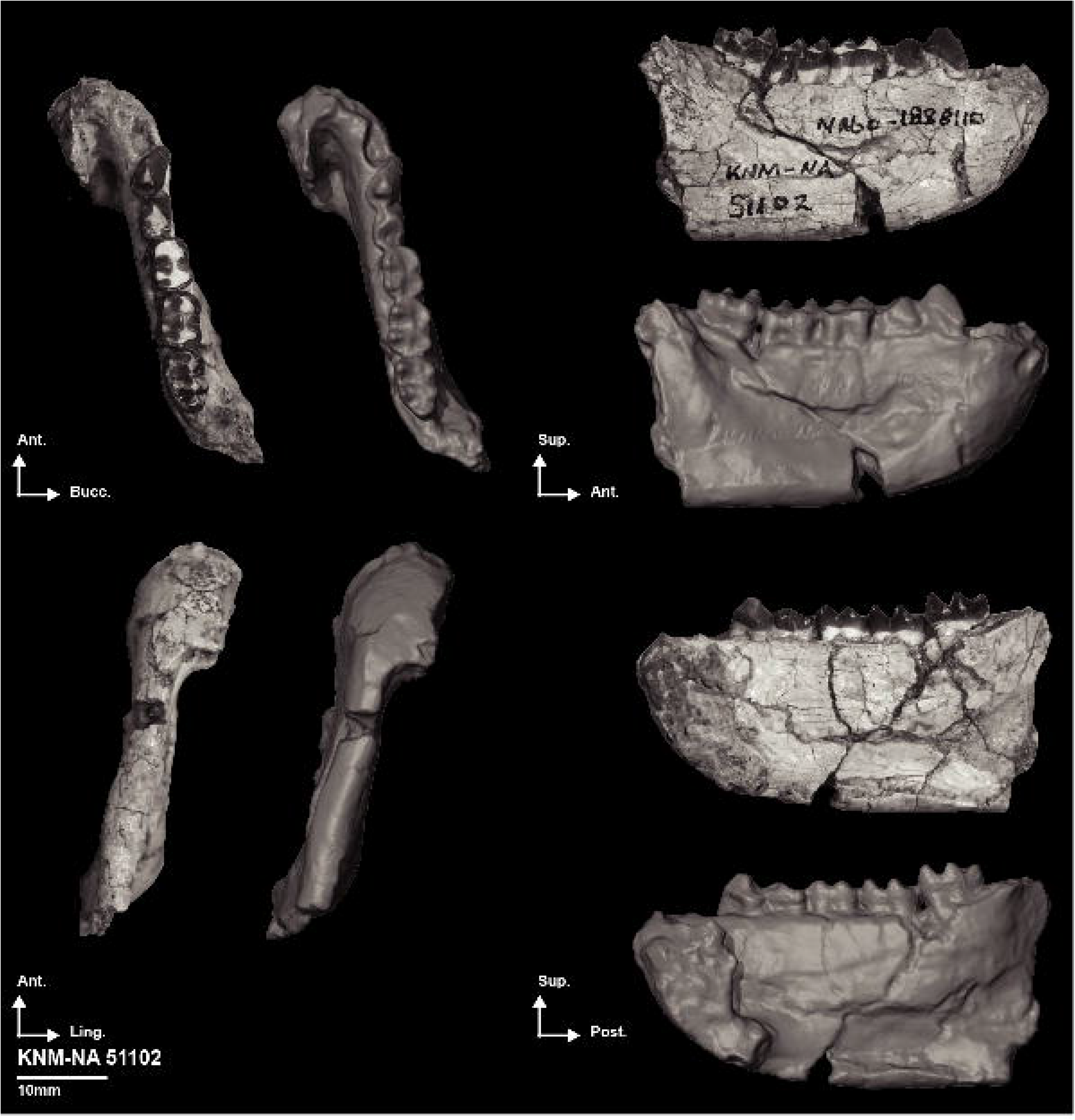
Photographs and 3D generated surface of the right partial hemimandible KNM-NA 51102.

Despite the abrasion of the incisor alveolar margin, sub-incisal hollowing is still visible below the right I_1_ - I_2_. The labial face of the symphysis is slightly convex and moderately sloping. The incisor arch is extremely narrow. The medial part of the left canine alveolus is fractured and glued. This precludes a precise assessment of the width of the anterior part of the mandible but the labial side of the symphysis and the preserved portion of the incisor row support the presence of a narrow mandibular arch. The maximum height of the symphysis is 20.4 mm. The planum alveolare is concave and is 11.4 mm long. The maximum breadth of the symphysis at the level of the superior transverse torus is 7.3 mm and is 6.7 mm at the inferior transverse torus level. The apex of the inferior transverse torus is set 4.3 mm superiorly from the inferior margin of the symphysis. The digastric fossa is palpable along the inferior margin of the symphysis and corpus. The morphology of the corpus is difficult to assess due to fractures and bone dislocation but it is quite thin and high. A fossa is visible below P_4_ / M_1_ but its depth is accentuated by the fractures. The inferior margin of the corpus is quite rounded below M_2_ and M_3_. The buccinator groove is faintly expressed and poorly roughened. The extramolar sulcus is 3.8 mm wide and, despite the missing ramus, the retromolar space appears to have been quite narrow. A mental foramen is present at P_4_ / M_1_ level. The foramen is located 5.7 mm superiorly to the inferior margin of the corpus. The maximum depth of the corpus is assessable at P_4_ / M_1_ (ca. 18.0 mm), at M_1_ / M_2_ (ca. 19.3 mm) and at M_2_ / M_3_ (ca. 19.7 mm).

The alveolus of the canine is preserved and is ca. 4.0 mm mesiodistally and 3.4 mm buccolingually. The apex of the P_3_ protoconid is worn and its anterior and posterior fovea are devoid of cuspulid. The P_3_ is 6.4 mm mesiodistally, with an occlusal length of 4.4 mm, and 3.5 mm buccolingually. The flange extends 6.2 mm superoinferiorly and the portion preserved of the protoconid elevates 1.7 mm proximally from the basin of the distal fovea. The P_4_ is severely worn, precluding a precise assessment of its morphology. A flange is present on mesiobuccal corner of the tooth. The preserved portion of the flange is 3.5 mm long. The P_4_ is 4.7 mm mesiodistally and 3.8 mm buccolingually. As for the P_4_, the occlusal surface of M_1_ is severely worn with connections of the dentine pools between the mesial and distal cusps. The M_1_ is 5.6 mm mesiodistally and the mesial lophid, with a buccolingual dimension of 4.6 mm, is slightly narrower than the posterior one (5.0 mm buccolingually). A wear differential between the lingual and buccal cusps is seen on M_2_, with the wear being more progressed buccally. The occlusal relief of the lingual cusp is moderate. The M_2_ is 6.1 mm mesiodistally and like M_1_, the mesial lophid is slightly narrower (5.0 mm) than the posterior one (5.3 mm). The M_3_ present a wear differential similar to M_2_. The hypoconulid is large (2.9 mm buccolingually) and is placed buccally on the tooth. The M_3_ is 7.2 mm mesiodistally and present an inverted breadth differential relative to M_1_-M_2_ regarding its mesial and distal lophids with a mesial lophid 4.9 mm wide and a distal lophid 4.7 mm wide. The length of the M_1_-M_3_ row is 19.5 mm.

#### KNM-NA 48912 (mandible with a complete tooth row)

KNM-NA 48912 is a 44.4 mm long mandible preserving a complete tooth row (Figure 4). The right hemimandible preserves all the teeth but the crown of the C_1_ is missing and the crown of the M_3_ is detached from the cervix and conserved separately. The inferior margin of the right hemimandible corpus is missing from P_4_ / M_1_ posteriorwards. The left hemimandible preserves all the teeth but shows severe damage to the left C_1_ and M_1_ crowns. Like the right hemimandible, the inferior margin of the corpus is missing from P_4_ / M_1_ posteriorwards. A coronal fracture is visible on the left corpus at M_1_ / M_2_ level. The posterior portion of the corpus and tooth row are dislocated medially due to this fracture. The cortical bone is fractured all along the surface of the specimen, presumably due to matrix infilling and expansion. The morphology of the left corpus anterior to the P_4_ / M_1_ junction and most of the superior aspect of the right corpus are nevertheless reliably assessable. The inferior transverse torus is damaged (Online Resource 9). A sagittal fracture with dislocation is visible on the left canine. A line of fracture set coronally is visible on the right P_4_. This fracture resulted in a loss of a flake of enamel on the lingual side of the tooth.

**Figure 4:**
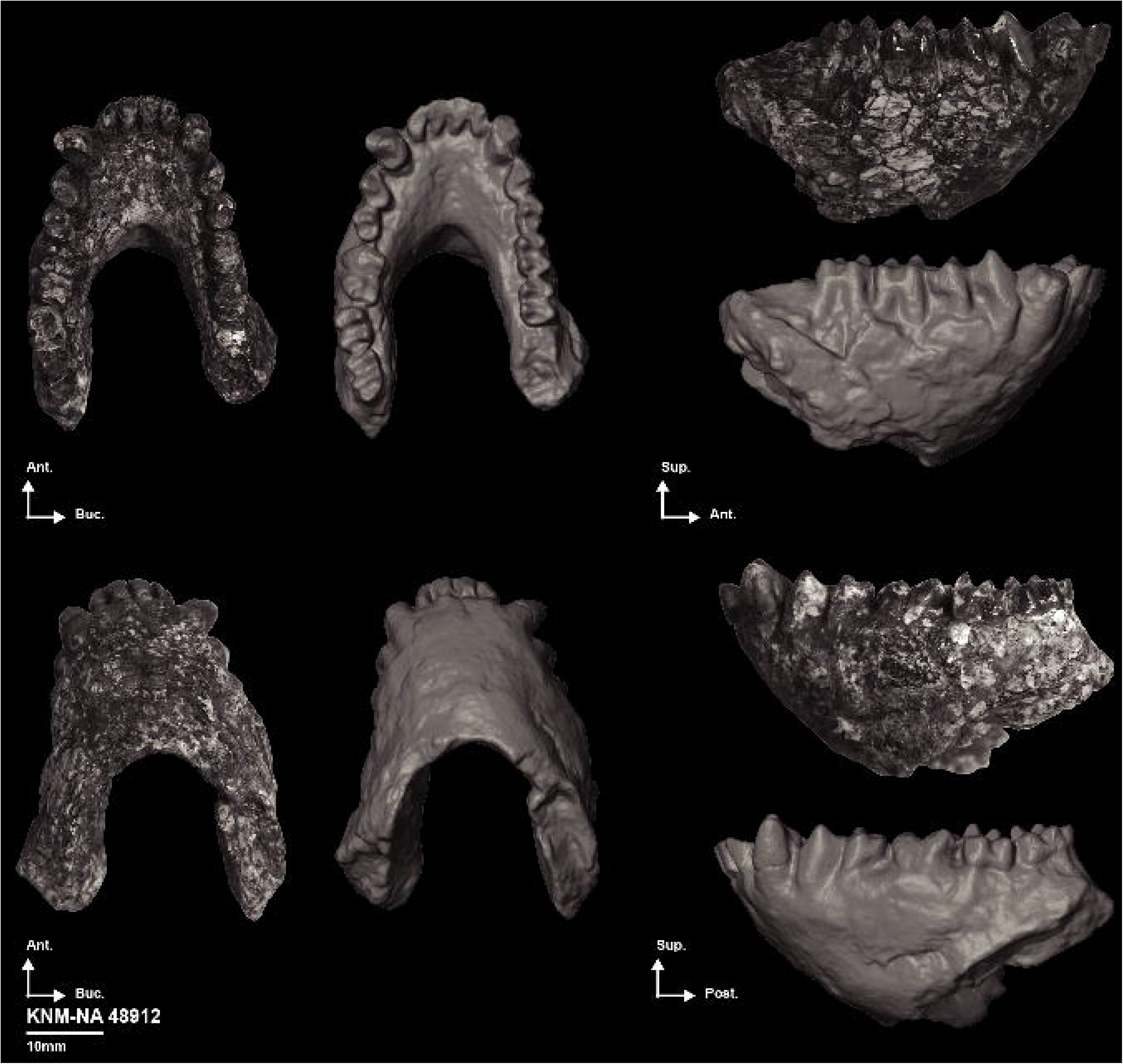
Photographs and 3D generated surface of the partial mandible KNM-NA 48912.

The labial face of the symphysis is slightly concave and sloping. The maximum height of the symphysis is 22.6 mm. The planum alveolare is poorly concave and is 12.9 mm long. The anterior portion of the mandible is quite large, with an inter-canine breadth of ca. 9.7 mm and an inter-P_3_ breadth of 13.0 mm (distance taken from alveolar lingual sides). The breadth of the symphysis at the superior transverse torus is 10.3 mm and 6.6 mm at the preserved portion of the inferior transverse torus. The posterosuperior aspect of the corpus is raised, which may indicate the presence of a developed lateral prominence but given the state of preservation of the specimen, we stay cautious regarding the presence of this feature in KNM-NA 48912. The lingual side of the incisors are quite worn and a large planar dentine exposure is visible. This basin is surrounded by a thin layer of enamel. The crown of I_2_ is quite asymmetrical with a raised distolingual corner. The incisor arch is ca. 10.3 mm long (distance taken from distal sides). The canine is quite robust (5.8 mm buccolingually and 4.7 mm mesiodistally for the right canine), suggesting a male status of KNM-NA 48912. The distal shoulder of the left canine is smooth and devoid of cuspulids. The P_3_ is sharp, with a flange extending 7.8 mm superoinferiorly on the right P_3_. A slight layer of dentine is visible on the apex of the P_3_ protoconid. The lingual face of the protoconid forms a strong pillar that delineates the anterior and posterior foveae. No cuspulids are visible on the mesial margin of the anterior fovea. The maximum MD dimension of the P_3_ is 7.1 mm and its occlusal MD length is 5.3 mm. The maximum BL dimension of the P_3_ is 3.5 mm. The protoconid and metaconid of P_4_ are quite worn, especially the protoconid, and the metaconid therefore appears higher than the protoconid. A flange, ca. 4.8 mm long on the right P_4_, is visible on the mesiobuccal corner of the tooth. The right P_4_ is 4.4 mm mesiodistally and 3.7 mm buccolingually. A wear differential on M_1_ - M_3_ in favor of the lingual cusps is observed. The relief of the lingual cusp is moderate. No mesial buccal cleft is visible on M_1_ and only a shallow distal buccal cleft is visible. The median buccal cleft is devoid of cuspulid. The right M_1_ is 5.6 mm mesiodistally and presents a breadth differential of the lophids in favor of the distal one (4.6 mm at the distal lophid vs. 4.55 mm at the mesial lophid). The M_2_ is devoid of any peculiar structures and differs from the M_1_ only by its dimensions and the higher depth of its mesial and distal buccal clefts. The M_2_ is 5.9 mm mesiodistally and also presents a breadth differential of the lophids in favor of the distal one (5.4 mm at the distal lophid vs. 5.1 mm at the mesial lophid). A large (2.8 mm buccolingually) and buccally placed hypoconulid is visible on M_3_. The M_3_ is 7.0 mm long, with a mesial and distal lophids similar in breadth (4.8 mm).

#### KNM-NA 48913 (mandible with a subcomplete toothrow)

KNM-NA 48913 is a 39.0 mm long mandible preserving most of the corpus of the left and right hemimandible and a subcomplete tooth row, lacking only the left and right I_2_ as well as the right I_1_ (Figure 5). The left I_1_ is detached from its alveolus and conserved separately. The right corpus is fractured at the P_4_ / M_1_ junction. This fracture results in a distortion of the bone. Significant portions of the bone, up to 2.0 mm long, are missing on the lingual side of the corpus. The right canine is broken at its base and glued back, which resulted in a slight offset at the cervix. Thin lines of fracture are visible on the labial face of the symphysis but they do not alter its morphology. A fracture located at the C_1_ / P_3_ junction of the left corpus intersects with the left margin of the genioglossal fossa. This fracture also leads to a significant loss of bone at the lingual side of the alveolar margin of C_1_. Another fracture runs obliquely from the P_4_ / M_1_ junction to the inferior margin. This fracture caused a lateral dislocation of the posterior portion of the corpus. The cortical bone of the inferior margin of the corpus is slightly abraded, but the level of abrasion observed does not significantly impact the measurements of the corpus. The left P_3_ and P_4_ are fractured at their cervix, with portions of the crown missing. They were then glued back in a position that resulted in a slight offset from the assumed original anatomical position. The superomesial corner of the left I_1_ is missing. The right P_4_ is fractured at mid-length, resulting in a slight dislocation of the distal portion of the crown relative to the mesial portion. The right P_4_ was also fractured at the cervix but the fracture did not result in a major offset of the tooth.

**Figure 5:**
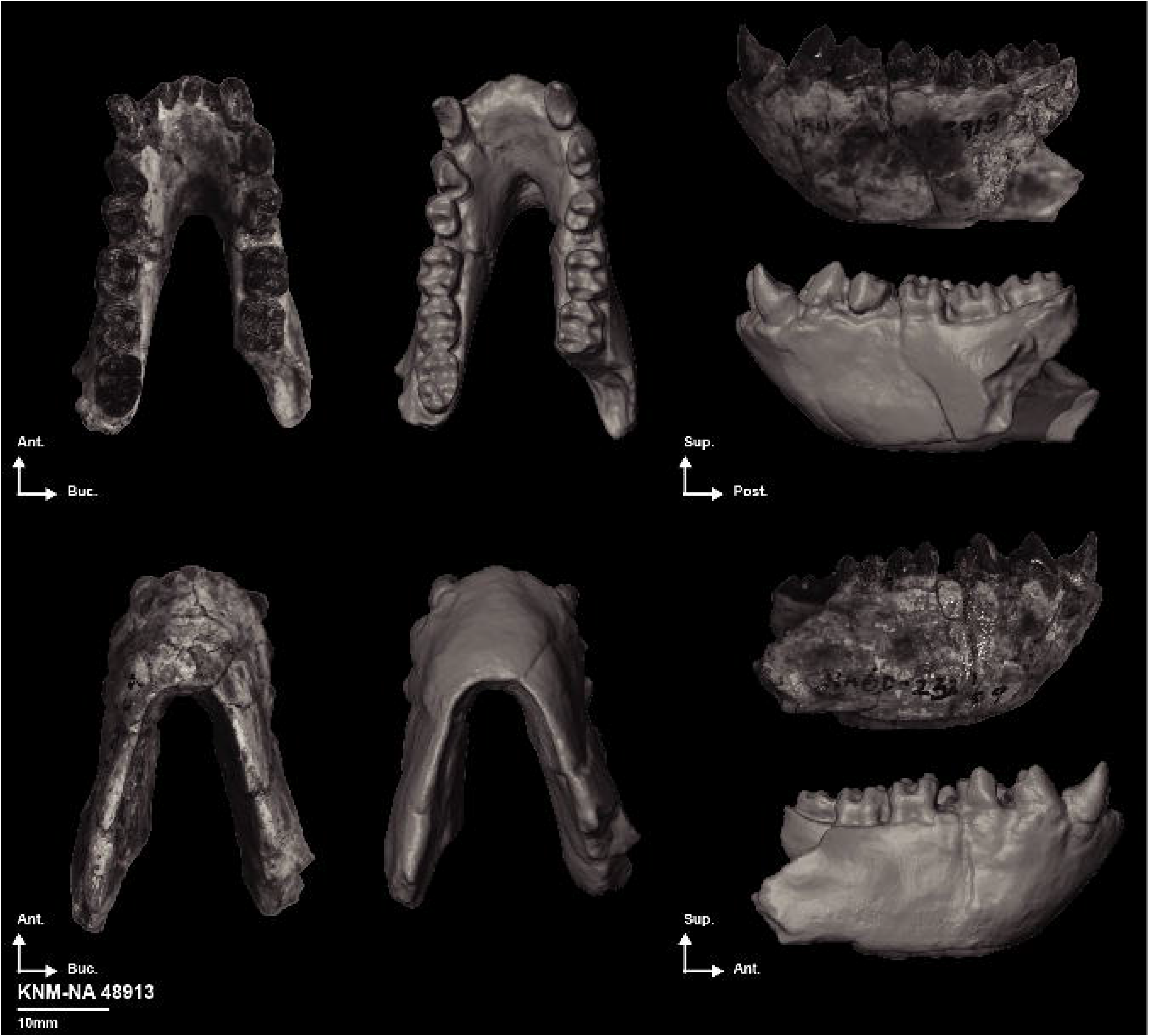
Photographs and 3D generated surface of the partial mandible KNM-NA 48913.

The labial face of the symphysis is straight, with a slight break in slope on its inferior portion. Sub-incisal hollowing is visible below I_2_ but does not extend to I_1_ level. The maximum height of the symphysis is 18.6 mm and the maximum breadth is 7.1 mm, a breadth measured at the level of the superior transverse torus. The breadth of the inferior transverse torus is 5.1 mm. The submandibular fossa is restricted to the inferior margin of the corpus and the apex of the inferior transverse torus is set 3.85 mm superiorly to the inferior symphyseal margin. The submandibular fossa extends posteriorly to M_1_ / M_2_ level (Online Resource 9). The planum alveolare is 12.2 mm long, slightly concave and is located below a narrow incisor arch. The incisor arch is approximately 7.5 mm wide (from the distal margins of the preserved incisor alveoli). The bicanine breadth is approximately 8.6 mm, from the lingual sides of the canines, but this dimension is most probably overestimated due to the fracture of the corpus. The inferior margin of the corpus is sharp. The superior aspect of the corpus is slightly hollowed below the premolars as the result of the anterior expansion of the lateral prominence on the inferior margin of the corpus. Lateral prominences are palpable inferior to the M_2_s. No rugosity or groove is visible for the insertion of the m. buccinator. The retromolar space seems minimal. No mental foramen is visible on the specimen despite the adequate preservation of the cortical bone surface. In lateral view, the inferior margin of the corpus is straight and does not present a clear distal bulging. The left corpus is 16.0 mm deep and 6.1 mm wide at the P_3_ / P_4_ junction. The right corpus is 13.9 mm deep and 6.4 mm wide at the M_1_ / M_2_ junction.

The crown of the I_1_ is symmetric but narrow mesiodistally. The tooth is 2.5 mm mesiodistally and 3.7 mm buccolingually. The buccal side of the I_1_ is quite convex and its lingual aspect is devoid of enamel apart from a small rim that surrounds the crown. The tooth preserves 6.3 mm of the root. The root is mesiodistally compressed and buccolingually elongated. At mid-length of its preserved portion, the root is 1.7 mm mesiodistally and 4.0 mm buccolingually. A sulcus is present on the mesial and distal side of the root and separates the buccal canal from the lingual canal. The canals are well visible at the fracture level. The canines are gracile and the apex of the crown of the left canine elevates 3.15 mm from its posterior margin. The left canine is 3.5 mm buccolingually and 4.8 mm mesiodistally. The P_3_ is sharp, with a 5.8 mm long flange. A small tubercle is visible on the mesiobuccal corner of the mesial fovea. The crown is slightly depressed on its buccodistal aspect. The maximum mesiodistal dimension of the right P_3_ is 5.7 mm and its occlusal length is 4.7 mm long. The maximum buccolingual dimension of the right P_3_ is 3.2 mm. The protoconid elevates 2.6 mm from the distolingual margin of the tooth. A 5.0 mm long flange is visible on the right P_4_. The protoconid is slightly worn but no dentine is visible on the apex of the crown of the metaconid. Even worn, the protoconid is higher than the unworn metaconid. The margins of the fovea are sharp but does not present any tubercle. The left P_4_ is 4.5 mm long and 3.6 mm wide. The molars are moderately worn, with a wear differential in favor of the lingual cusps. The mesial buccal cleft is extremely shallow on the M_1_. The breadth differential between the lophids is marked, with a 4.6 mm wide mesial lophid and a 4.7 mm wide distal lophid on the right M_1_. The right M_1_ is 5.5 mm mesiodistally. No remarkable features are visible on the M_2_’s. The right M_2_ is 5.5 mm mesiodistally, and has a mesial lophid (5.0 mm) subequal in breadth to the distal lophid (4.9 mm). The hypoconulid is quite large (ca. 2.3 mm) and is centrally placed on the left M_3_. While the mesial lophid of the M_3_ is oriented perpendicular to the mesiodistal axis of the tooth, its distal lophid is set at an oblique angle regarding the mesiodistal axis of the tooth. The breadth differential of the lophids is in favor of the anterior one (4.7 mm buccolingually at the mesial lophid and 4.41 mm buccolingually at the distal lophid). The left M_3_ is 6.7 mm long. The length of the molar tooth row is 18.0 mm.

#### KNM-NA 54785 (mandible with a subcomplete toothrow)

KNM-NA 54785 is a 49.5 mm long mandible preserving most of the right corpus and the left corpus, approximately up to the M_2_ level (Figure 6). The left M_2_ is detached from the rest of the mandible. The specimen also preserves a subcomplete toothrow, excluding the incisors. Matrix expansion led to minor bone dislocation and fractures but the morphology and morphometry of the specimen can be reliably assessed. A large coronal fracture is visible on the right corpus. This fracture engendered a slight lateral displacement of the posterior part of the corpus and significant cortical bone loss, up to 4.3 mm anteroposteriorly, on the buccal side of the corpus. A fracture, running from the junction of the I_1_ / I_2_ alveolus down the midline of the inferior symphyseal margin, resulted in minor bone dislocation but significant bone loss on the symphyseal labial side. Approximately 2.0 mm of bone is missing on some areas of the symphyseal labial side. A third line of fracture joins the previously described fracture at the level of the superior transverse torus, forming a triple junction that resulted in loss of cortical bone on the posterior aspect of the superior transverse torus. A fourth fracture, set at the P_4_ / M_1_ junction, led in a complete loss of the cortical bone on the lingual side and in medial dislocation of the posterior part of the corpus. The crown of the left canine is missing and the apex of the P_3_ protoconid is damaged. The apex was glued back, causing the tip of the P_3_ to shift in relation to the rest of the crown. The right premolars are badly damaged, with their buccal side missing. The protoconid of the right M_1_ and the metaconid of the left M_1_ are also missing. The hypoconulid of the M_3_ is missing and the crown of the entoconid and metaconid is badly damaged, precluding an assessment of their morphology.

**Figure 6:**
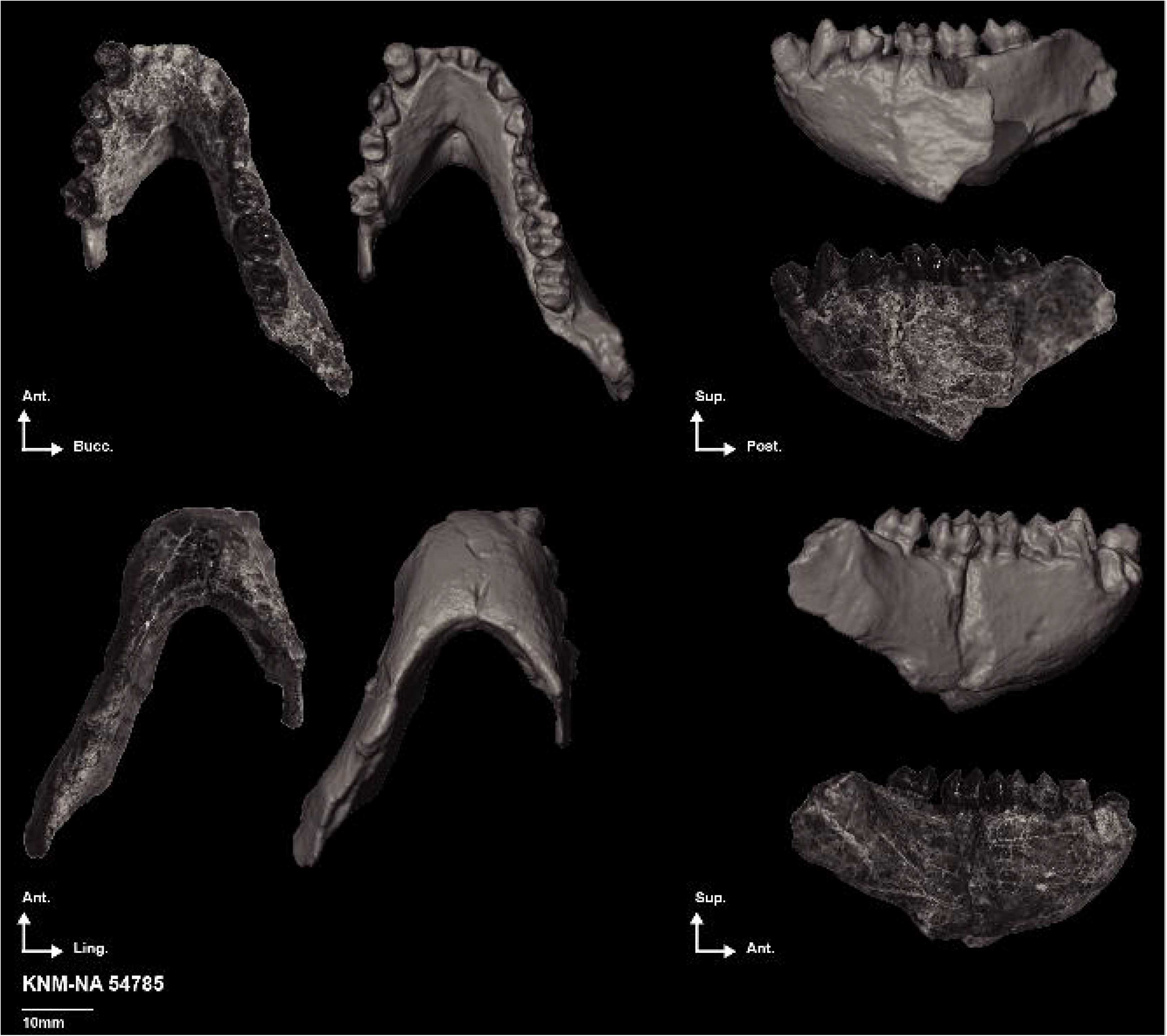
Photographs and 3D generated surface of the partial mandible KNM-NA 54785.

A slight sub-incisal hollowing is palpable below the left and right I_2_. The labial side of the symphysis is convex and sloping. The midline of the symphyseal labial side is damaged precluding an assessment of the presence / absence of a median mental foramen. The planum alveolare is moderately concave and is 11.4 mm long. Superior to the planum alveolare is an obtuse incisor arch. The incisor arch is 9.3 mm wide (taken from the distal margins of the I_2_ alveoli). The maximum breadth of the symphysis is 9.1 mm. This dimension is reached at the level of superior transverse torus. The inferior transverse torus is large, forming a developed shelf below the genioglossal fossa. The breadth of the symphysis at the level of the inferior transverse torus is 6.9 mm. The apex of the inferior transverse torus is set 4.8 mm superiorly from the inferior margin of the symphysis. The posterior expansion of the superior and inferior transverse tori defines a slight sulcus (i.e., intertoral sulcus) on the lingual side of the corpus, well palpable below M_1_. The submandibular fossa is palpable and extends posteriorly to the M_1_ / M_2_ level. A fossa is palpable on the buccal side of the corpus below the premolars, highlighting the anterior extension of the lateral prominence. Two mental foramina are visible below the P_4_ / M_1_ junction. The superior one is larger than the inferior one and its center is located 7.2 mm from the adjacent inferior margin of the corpus. The minute foramen is located inferoposteriorly to the larger one and is situated 5.6 from the inferior margin of the corpus. The right corpus is 17.5 mm deep and 7.3 mm wide at the M_1_ / P_4_ level, 17.7 mm deep at M_1_ / M_2_ level (taken anterior to the fracture level). A rough groove marks the insertion of m. buccinatorius inferior to M_3_. The extramolar sulcus is approximately 4.8 mm wide and the retromolar space is minimal.

Our description of the dental anatomy will focus on the left C_1_, P_3_, P_4_, M_1_ and on the right M_2_ and M_3_. The left canine is large, with a maximum buccolingual dimension of 4.6 mm and a maximum mesiodistal dimension of 6.3 mm. The posterior part of the crown is slightly damaged but no tubercle is visible on its surface. The alveolus of the right canine is 5.8 mm mesiodistally. The left P_3_ is sharp, with an 8.8 mm long flange on its buccomesial corner. The maximum mesiodistal dimension of the P_3_ is 8.25 mm, with an occlusal length of 5.6 mm. The maximum buccolingual dimension of the P_3_ is 3.6 mm. No accessory tubercles are visible on the margins of the distal and mesial fovea. The protoconid elevates 3.7 mm from the distolingual margin of the tooth, with the lingual side of its apex being only slightly worn. The left P_4_ is 5.2 mm mesiodistally and 4.1 mm buccolingually. A flange of 5.1 mm is visible on its mesiobuccal corner. The apex of the metaconid is damaged and the protoconid is slightly worn. In this condition, the metaconid is higher than the protoconid. A shallow cleft is visible on the buccodistal corner of the tooth. The left M_1_ is 5.9 mm mesiodistally. The mesial buccal cleft is faintly expressed compared to the distal buccal cleft. A wear differential is visible between the lingual and buccal cusps. The dentine basins of the buccal cusps are not connected. The distal lophid of the M_1_ is 4.9 mm and the mesial lophid is 4.7 mm. Despite a significant occlusal relief, the crown of the right M_2_ is high at the level of the lingual notch. The lingual cusps are only slightly worn, with small circles of dentine visible on the buccal aspects of the metaconid and entoconid. The M_2_ is 6.7 mm mesiodistally and has lophids subequal in breadth (5.8 mm). The maximum preserved length of the M_3_ is ca. 7.2 mm. The breadth of the tooth at the mesial lophid is 5.3 mm.

#### KNM-NA 46288 (mandibular fragment)

KNM-NA 48912 is a 41.22 mm long right mandibular fragment preserving P_4_ and M_1_ and the basal most portion of the M_2_ crown (Figure 7). A maximum of ca. 15.8 mm of the superoinferior dimension of the corpus is preserved below M_1_. A slight portion of the anterior aspect of the ramus is also preserved. The bony anatomy is largely distorted by matrix expansion. There is no cortical bone on the inferior margin of the fragment which indicates that the inferior most margin of the corpus is preserved. The P_4_ is perfectly preserved but a thin line of fracture is visible on the median buccal cleft of M_1_ and a pit of enamel is seen on the distolingual aspect of its entoconid. The description will focus on the dental anatomy of the specimen.

**Figure 7:**
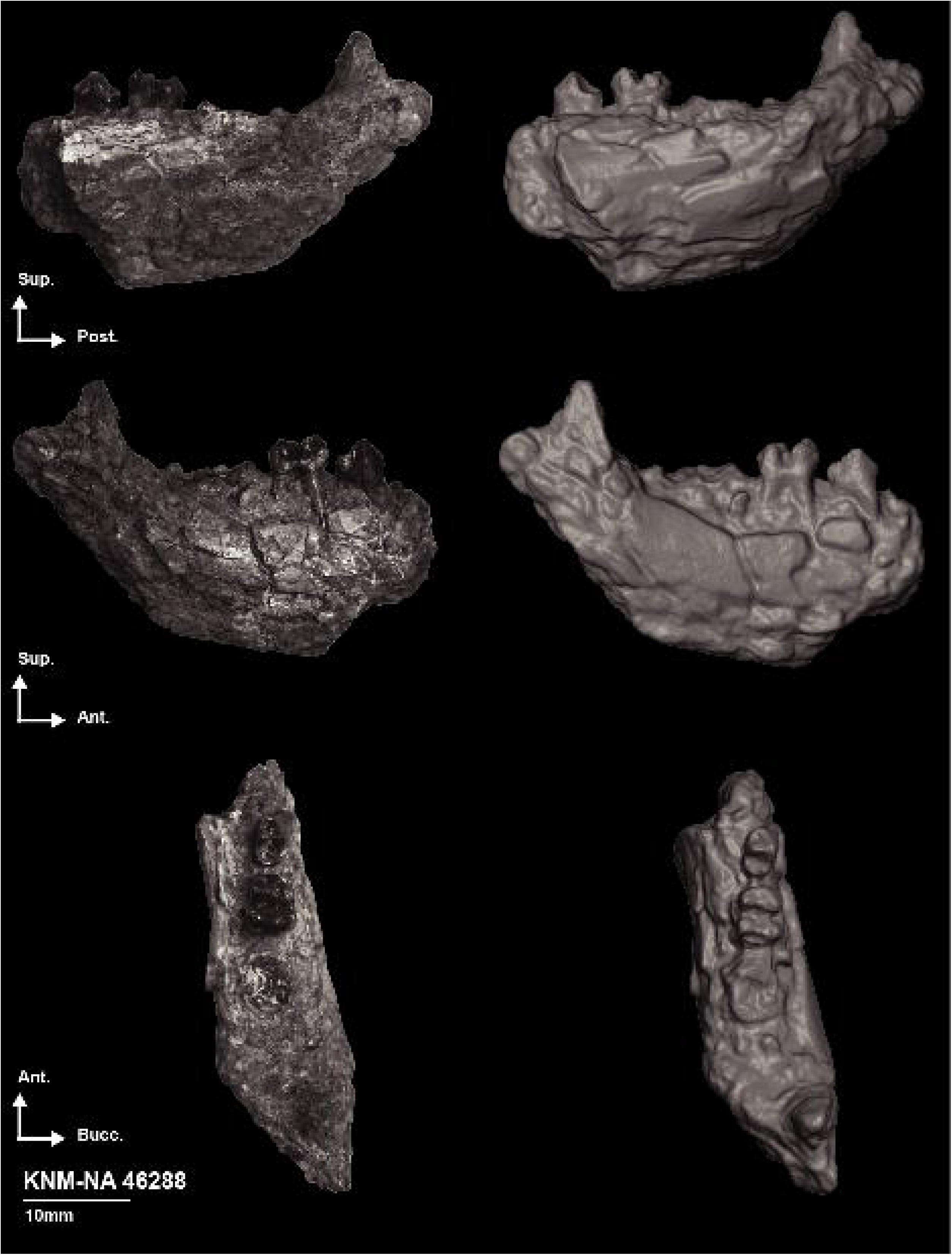
Photographs and 3D generated surface of the right partial hemimandible KNM-NA 46288.

The protoconid and metaconid of the P_4_ are slightly worn and heteromorphic with the protoconid higher than the metaconid. The anterior and distal fovea are quite deep. A cuspulid is visible on the distolingual side of the distal fovea. A 5.8 mm long flange is visible on the mesiobuccal corner of the tooth. The P_4_ is 4.8 mm mesiodistally and 3.4 mm buccolingually. A wear differential is visible on the lingual and buccal cusps of the M_1_, with the lingual cusps less worn than the latter. The relief of the lingual cusps is quite marked. The mesial and distal buccal clefts are easily identifiable. The M_1_ is 5.7 mm mesiodistally and the breadth differential between the mesial and distal lophid is in favor of the latter (4.5 mm at the distal lophid vs. 4.4 mm at the anterior one).

#### KNM-NA 51103 (mandibular fragment)

KNM-NA 51103 is a 18.7 mm long fragment of a left mandible preserving M1 - M3 and ca. 10 mm superoinferiorly of the superior portion of the corpus (Figure 8). A thin line of fracture is visible below M_2_ which resulted in a loss of cortical bone inferior to 0.50 mm on the lingual side of the fragment. The teeth are perfectly preserved and slightly worn. A slight oblique ridge is visible on the buccal side and marks the location of the lateral prominence and the presence of an extramolar sulcus. The relief of the cusp is moderate (Online Resource 12), elevating 1.4 mm from the notch, and with a crown measuring 1.3 mm (NC and NR variables *sensu* Benefit and Pickford, 1986). Small circles of dentine are visible on the buccal cusps of M_1_ and M_2_ while M_3_ is barely unworn. No remarkable features are visible on the molars and they display a typical colobine morphology. The breadth differential of the lophids are in favor of the distal one for M_1_ and M_2_ while the reversed is observed for M_3_. The M_3_ hypoconulid is moderate in breadth (2.25 mm) and placed buccally. The length of the molar tooth row is 17.3 mm.

**Figure 8:**
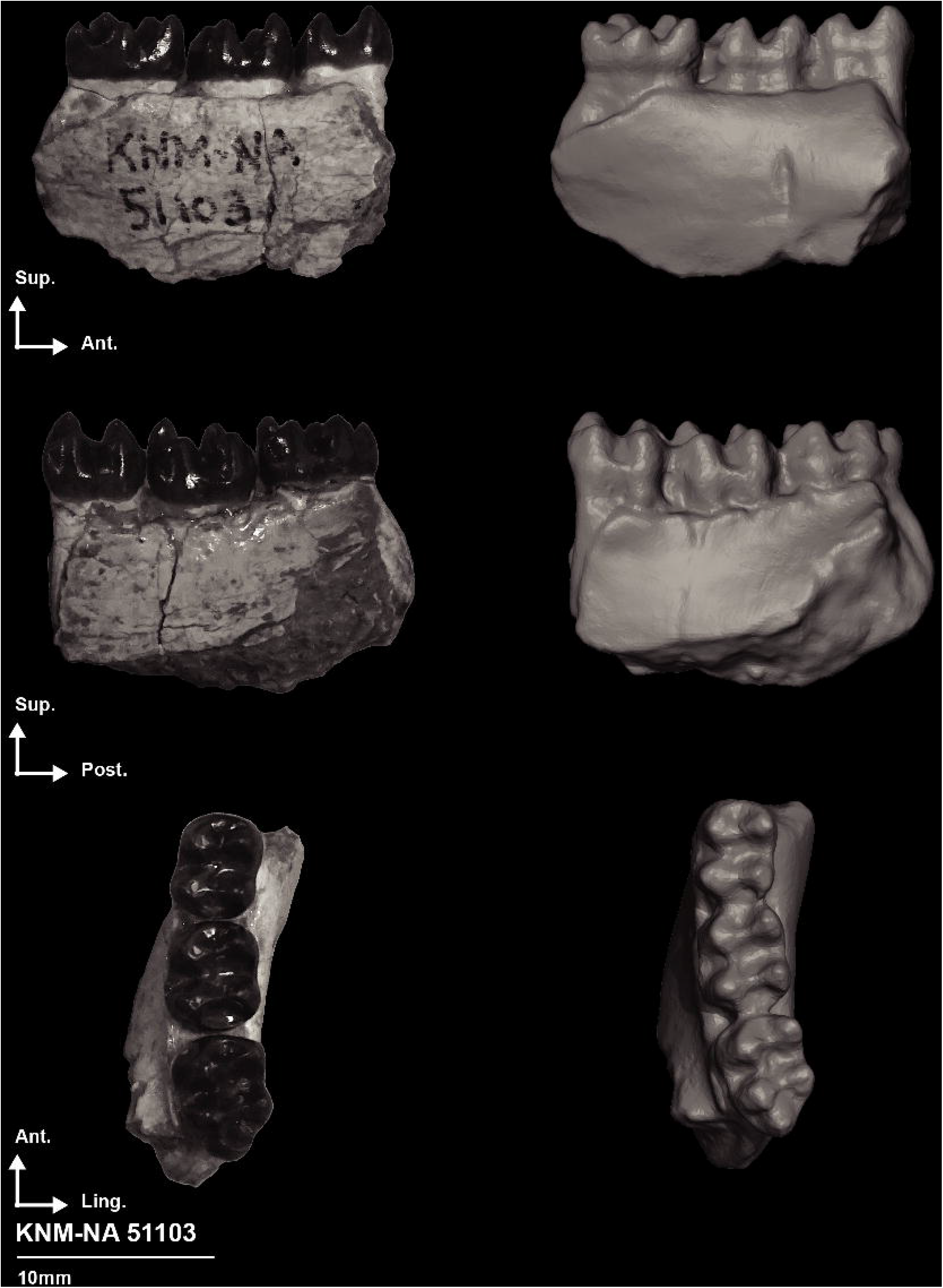
Photographs and 3D generated surface of the left partial hemimandible KNM-NA 51103.

### DENTAL MORPHOMETRY AND COMPARATIVE MORPHOLOGY

#### Bivariate and univariate dental data

In the geometric mean of the M_1_ and M_2_, the Nakali specimens overlap with the interquartile range of *Pre. chrysomelas* and *Tr. cristatus,* and fit within the upper quartile of *Pro. verus* (see the grey envelope in Figure 9). The Nakali specimens are also consistent in size with the M_1_ dimensions of *Mi. tugenensis* (KNM-BN 1740) and are slightly smaller than the isolated molar KNM-NA 305 from Nakali and CHO-BT 78 from Beticha (Chorora), from Nawata Colobinae gen. indet. sp. A KNM-LT 24107, and the small colobine from Lemudong’o KNM-NK 36514 (Figure 9). In M_2_ geometric mean, the Nakali specimens are similar to the Lukeino M_2_ KNM-TH 36742. The geometric mean of the M_1_ and M_2_ is a raw estimation of the overall dental size, as megadonty and increased buccal flaring have a great impact on dental size. However, buccal flaring is reduced in colobines compared to cercopithecines and basal cercopithecoids (Benefit, 1993). This is exemplified here by *V. macinnesi*, which falls within the interquartile of *Pi. badius*, and lower quartile of *Co. guereza* and *N. larvatus* (Figure 9) despite having an absolutely smaller size (Blue et al. 2006).

**Figure 9:**
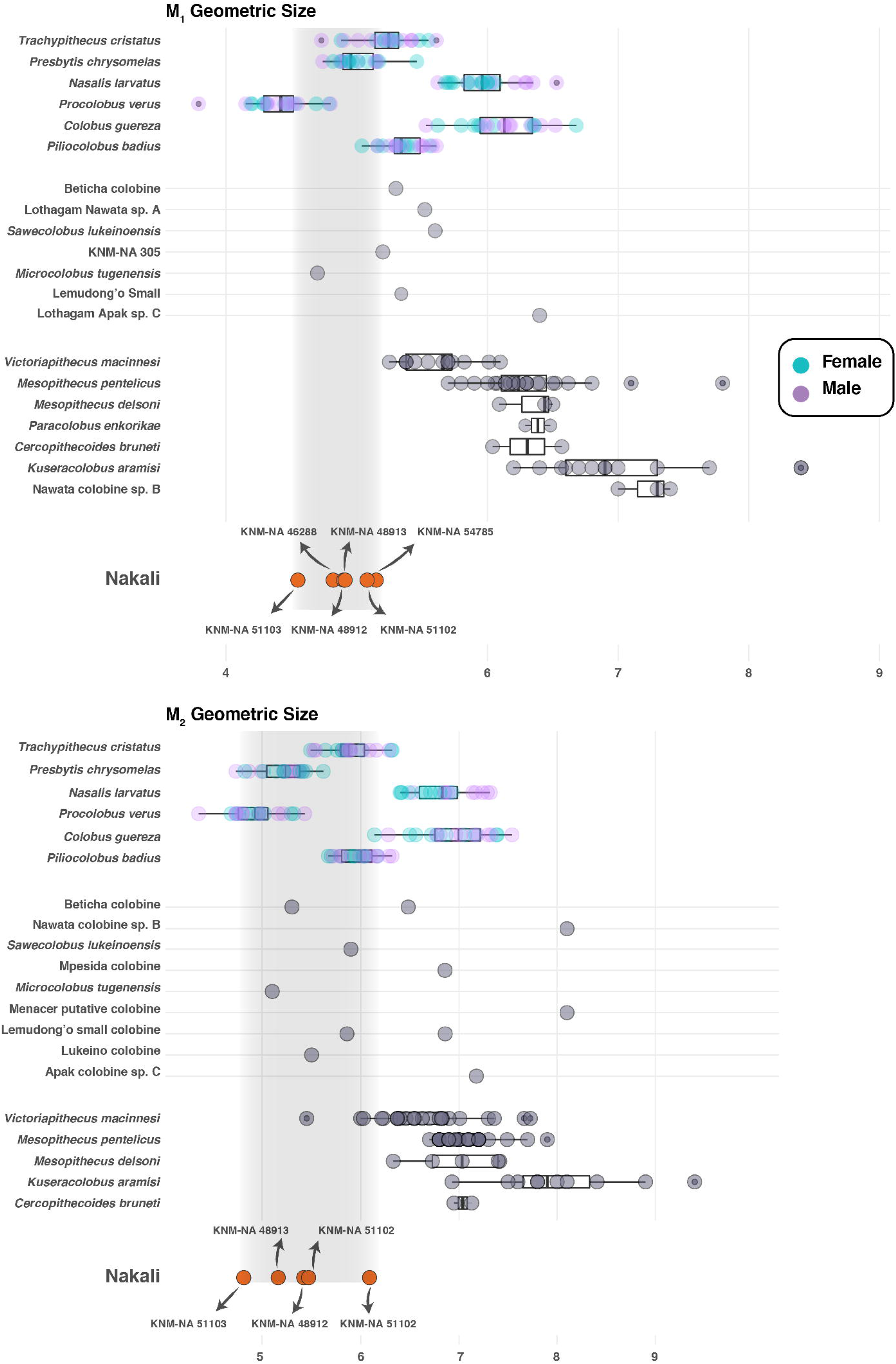
Boxplots of M_1_ and M_2_ geometric size of extant and fossil colobines. Boxplots with first, third quartile, and median (black line).

Canine relative length (i.e., C_1_ length / M_2_ GM) is differentiating African colobines from Asian colobines, with male specimens of the former displaying a markedly elongated canine (Online Resource 13). The magnitude of dimorphism in the Nakali specimens (KNM-NA 54785 and KNM-NA 48913) is comparable to that of *Tr. cristatus* but does not match the extremely elongated lower canine of *Pro. verus* and *Pi. badius* males. None of the fossil African colobine *Pa. enkorikae* and *K. aramisi* fit with the canine shape of male *Pro. verus* and *Pi. badius*. Similarly, *Me. pentelicus* and *Me. delsoni* does not present a canine dimorphism similar to *Pro. verus* and *Pi. badius*, but rather shows a range of variation consistent with that of *N. larvatus* (Online Resource 13). The C_1_ of *V. macinnesi* KNM-MB 18993 is also extremely reduced in relative length compared to most extant colobines.

When the natural logarithm of canine length (mesiodistal dimension) is regressed on the natural logarithm of M_2_ GM, *Pi. badius* and *Pro. verus* shows extremely long canines given their M_2_ GM (Figure 10). The y-intercept of the Colobini regression line (a = 0.88, y-intercept = 0.32) is higher than that of the Presbytini (a = 0.84, y-intercept = 0.25). The magnitude of sexual dimorphism of the Nakali specimens corresponds to the Presbytini regression line, as is also the case for *Me. pentelicus*. The canine of male specimens of *K. aramisi* (ASI-VP 1/87 and ASI-VP 1/306) and *Pa. enkorikae* (KNM-NK 44770 and KNM-NK 36515) is intermediate in terms of relative canine elongation and fit between *Co. guereza* and *N. larvatus*, presenting a pattern comparable to the Nakali male KNM-NA 54785 (Figure 10). The absolute canine dimensions of KNM-NA 54785 are also similar to those of the Chorora specimen CHO-BT 123 and of the Colobinae gen. indet. sp. A KNM-LT 26383 from the Nawata Formation.

**Figure 10:**
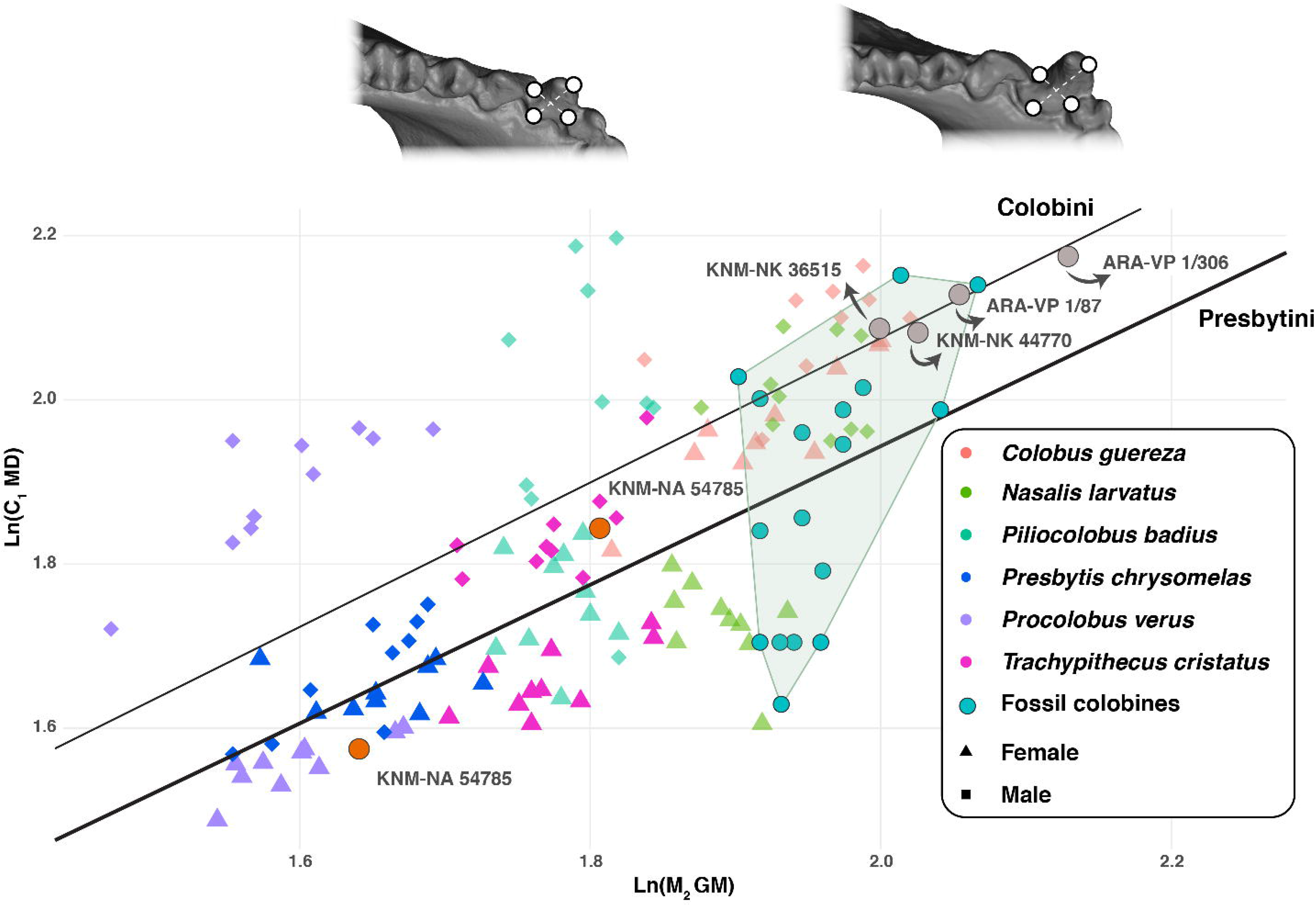
Least-squares regression of the natural logarithm of C_1_ MD on the natural logarithm of M_2_ GM of extant and fossil colobines.

Least-squares regression of the natural logarithms of the square roots of the crown dimension of C_1_ on occlusal dimensions of P_3_ enables us to distinguish between male and female of extant colobines (Figure 11). Indeed, absolute P_3_ and C_1_ crown dimensions are larger in males than in females. KNM-NA 48913 present absolute P_3_ and C_1_ dimensions in the size range of female *Pre. chrysomelas* and *Pro. verus* specimens while KNM-NA 54785 is similar to that of male *Pro. verus* and *Tr. cristatus*.

**Figure 11:**
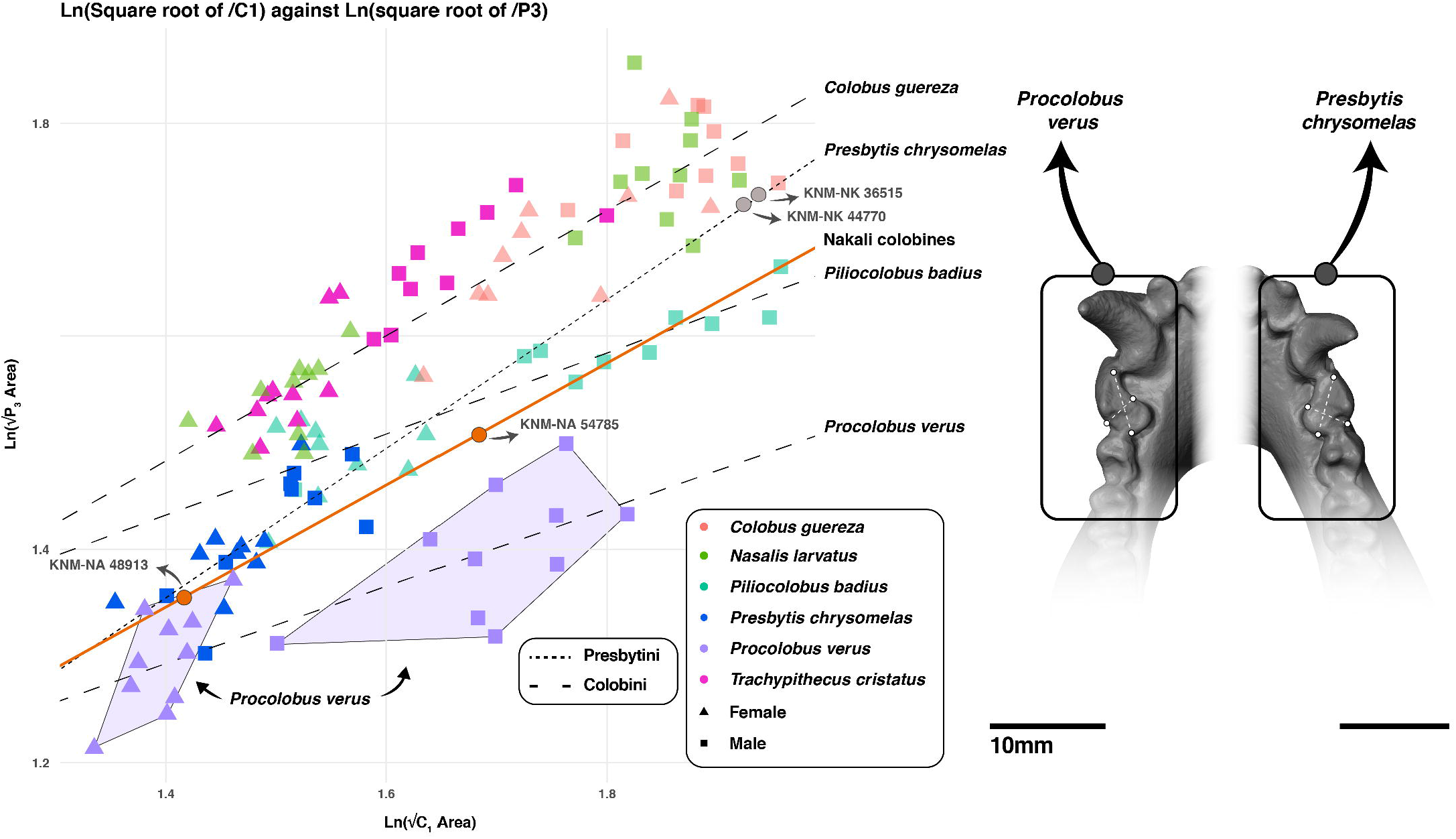
Least-squares regression of the natural logarithm of the square root of P_4_ area on the natural logarithm of the square root of C_1_ area of extant and fossil colobines.

The magnitude of difference in absolute P_3_ and C_1_ dimensions between KNM-NA 48913 and KNM-NA 54785 matches that of *Pro. verus* but are greater than that of *Pre. chrysomelas*. Slope differences are observed between the regression lines of *Pro. verus* (a = 0.36, y-intercept = 0.79) - *Pi. badius* (a = 0.38, y-intercept = 0.90) on one side and *Co. guereza* (a = 0.59, y-intercept = 0.66) - *Pre. chrysomelas* (a = 0.70, y-intercept = 0.38) on the other side. In the latter group, the P_3_ is larger in geometric mean for any given C_1_ GM (Figure 11). The regression line of KNM-NA 48913 and KNM-NA 54785 is similar to that of *Co. guereza* and *Pre. chrysomelas*, demonstrating their relatively large P_3_ relative to C_1_ area.

The P_3_ of male colobines is more elongated compared to female specimens (Online Resource 14). When compared with the similar-sized *Pro. verus*, KNM-NA 51102 and KNM-NA 48913 falls in the female range while KNM-NA 48912 and KNM-NA 54785 falls in the male range (Online Resource 14). P_4_ crown shape ratio (CSR: buccolingual / mesiodistal dimensions) differentiates the Colobini taxa from the Presbytini in our sample (Figure 12). The former present a relatively narrow (buccolingually) P_4_ compared to the latter, and notably that of *Pre. chrysomelas*. The Nakali specimens present a range of variation of P_4_ CSR values overlapping with that of Colobini (Figure 12). It also overlaps with *Kuseracolobus aramisi*, *Me. pentelicus*, *Ce. bruneti* (TM 266 03-099), *S. lukeinoensis* (OCO 607’10), and the Ngorora isolated P_4_ KNM-BN 1251. *Mi. tugenensis* KNM-BN 1740 present a slightly higher P_4_ CSR compared to the Nakali sample. The high P_4_ CSR values of *Pa. enkorikae* and the small Lemudong’o colobine (KNM-NK 36514) are outside the range of variation of that of the Nakali specimens (Figure 12). *V. macinnesi* present a bimodal distribution inconsistent with the range of variation observed in extant and fossil colobines. Most of the Nakali colobines falls in the interquartile range of *V. macinnesi*.

**Figure 12:**
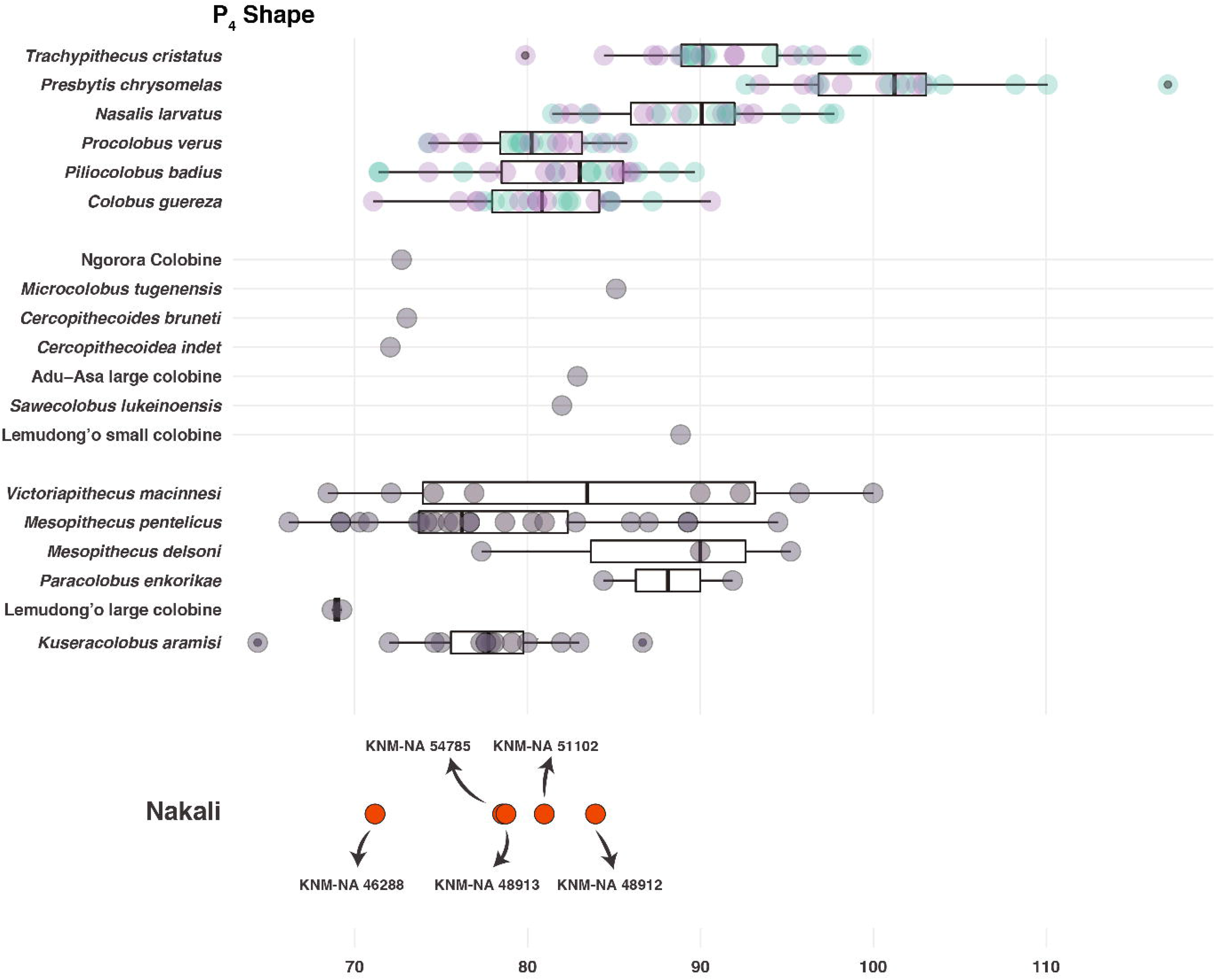
Boxplots of the P_4_ shape index of extant and fossil colobines. Boxplots with first, third quartile, and median (black line). Note the enlarged P_4_ of extant Presbytini.

The P_4_ area / M_1_ area ratio distinguishes *Tr. cristatus*, *Pre. chrysomelas*, and *Co. guereza* from *Na. larvatus*, *Pro. verus*, and *Pi. badius* (Figure 13). In the latter group, the area of the P_4_ is reduced in relative to the area of the M_1_. The Nakali specimens present index values and a range of variation comparable to that of *Na. larvatus*, *Pro. verus*, and *Pi. badius*, illustrating their relatively small P_4_. Within the Nakali sample, the high index value of KNM-NA 54785 is to be noticed (Figure 13). The lower index values of the Nakali specimens are comparable to the interquartile range of *Me. pentelicus* and *Me. delsoni*, and to the lower quartile of *V. macinnesi*. Nakali values are also matched by *Ce. bruneti* (TM 266 03-099) and *Pa. enkorikae* (KNM-NK 44770) but are outside the range of variation of *K. aramisi*. Additionally, *S. lukeinoensis* (OCO 607’10) present a slightly lower index value compared to the Nakali sample while the small Lemudong’o colobine (KNM-NK 36514) shows a slightly higher one (Figure 13). The index value of *Mi. tugenensis* KNM-BN 1740 is extremely low and fits only with the lower quartile of *Pro. verus* and *Na. larvatus*.

**Figure 13:**
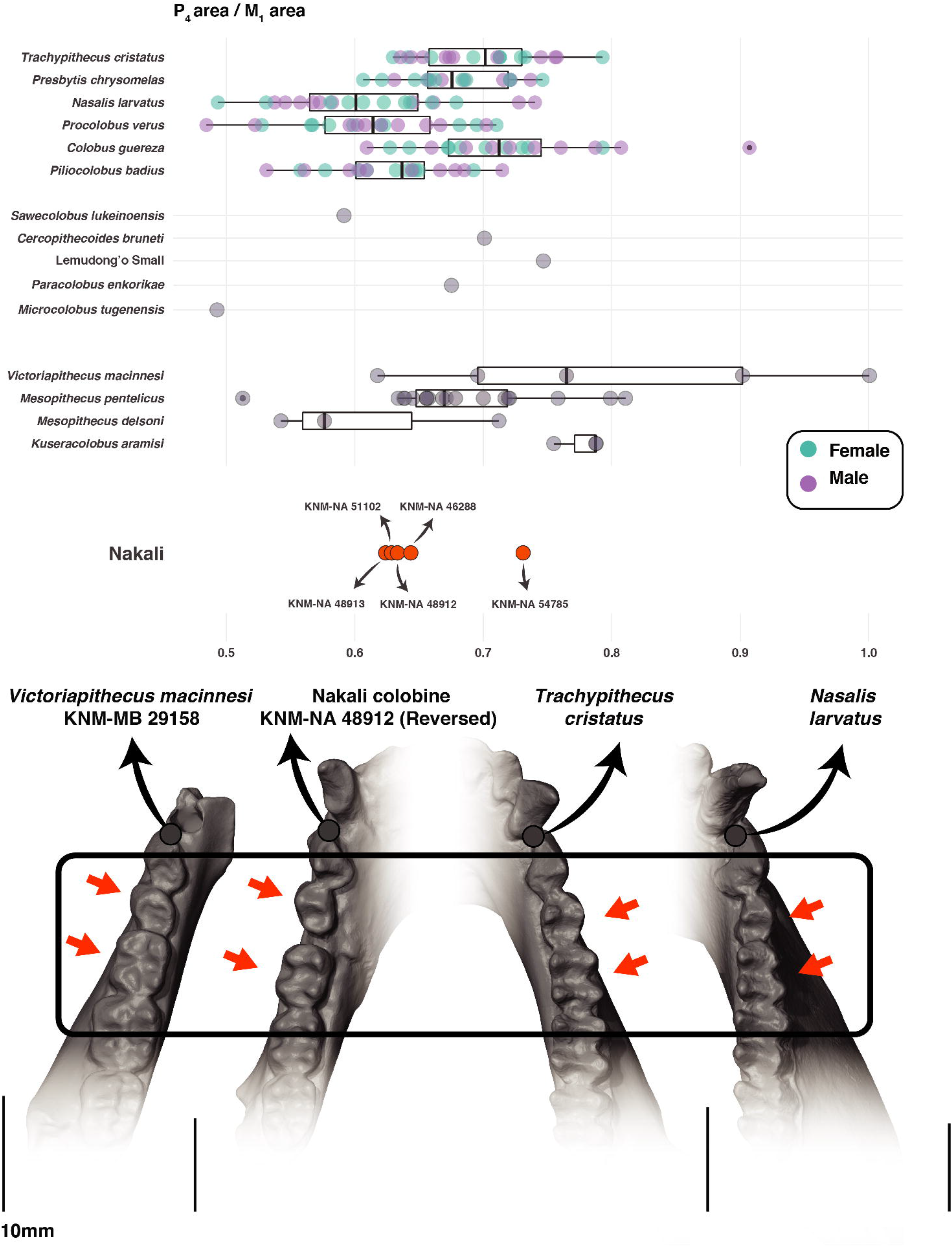
Boxplots of the P_4_ area / M_1_ area index of extant and fossil colobines. Boxplots with first, third quartile, and median (black line). Fossil cercopithecids and extant colobines are shown on the bottom of the graph to compare them and to illustrate the relatively enlarged M_1_ of *N. larvatus* and the relatively enlarged P_4_ of *Tr. cristatus*.

The index of the breadth differential of the M_1_ lophids distinguishes *Tr. cristatus* from all other extant colobines (Figure 14). *Tr. cristatus* shows, on average, mesial and distal lophids of nearly equal breadth. In contrast, a marked difference in breadth of the lophids is seen on the M_1_ of *Na. larvatus*, *Pro. verus*, and *Pi. badius*, with the mesial lophid narrower than the distal lophid. The index values of the Nakali specimens are most comparable to that of *Pre. chrysomelas* and *Tr. cristatus,* hence illustrating lophids of nearly equal breadth (Figure 14). The narrowness of the M_1_ mesial lophid of KNM-NA 51102 is noticeable. Apart from KNM-NA 51102, the Nakali specimens fall within the interquartile range of *K. aramisi* and *V. macinnesi*, but outside that of *Me. pentelicus* and *Me. delsoni*. They are also comparable to the small Lemudong’o colobine (KNM-NK 36514) and the Nawata colobine sp. A KNM-LT 24107, but have lower values than KNM-NA 305, *Mi. tugenensis* KNM-BN 1740, and *Ce. bruneti*.

**Figure 14:**
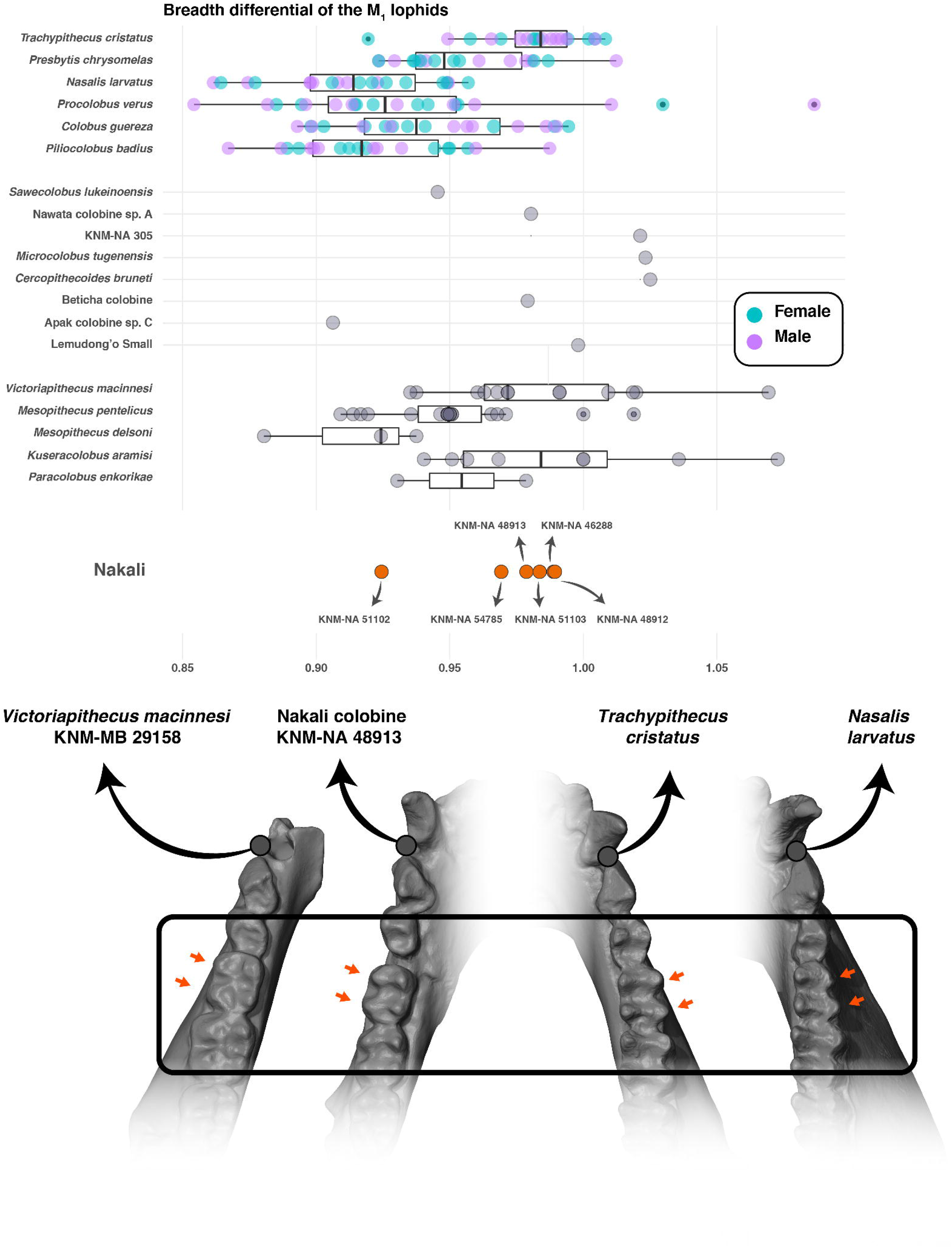
Boxplots of the breadth differential of the M_1_ lophids of extant and fossil colobines. Boxplots with first, third quartile, and median (black line). Fossil cercopithecids and extant colobines are shown on the bottom of the graph to compare them and to illustrate the subequal breadth of the M_1_ lophids of *Tr. cristatus* and the marked breadth differential of the M_1_ lophids of *N. larvatus*.

The M_1_/M_2_ length differential index permit to distinguish Presbytini. High ratio values are seen in *Pre. chrysomelas*, while relatively lower values are observed in *Na. larvatus* and *Tr. cristatus* (Figure 15). This index highlights the presence of a relatively short M_1_ in *Na. larvatus* and *Tr. cristatus*, but a relatively long M_1_ in *Pre. chrysomelas*. Colobini ratio values are intermediate between that of the above-mentionned Presbytini. An extensive range of variation is observed in the Nakali sample, with some specimens showing a relatively short M_1_ (i.e., KNM-NA 54785 and KNM-NA 51102) while others a relatively long M_1_ (i.e., KNM-NA 48913 and KNM-NA 51103). The high ratio values of KNM-NA 48913 and KNM-NA 51103 are similar to those of *S. lukeinoensis* (OCO 607’10) and *Mi. tugenensis*. Overall, the Nakali specimens present ratio values comparable to those of other Miocene colobines (Figure 15). However, they can clearly be discriminated, along with other Miocene colobines, from the extremely low ratio values, and hence reduced M_1_ length, of *V. macinnesi*.

**Figure 15:**
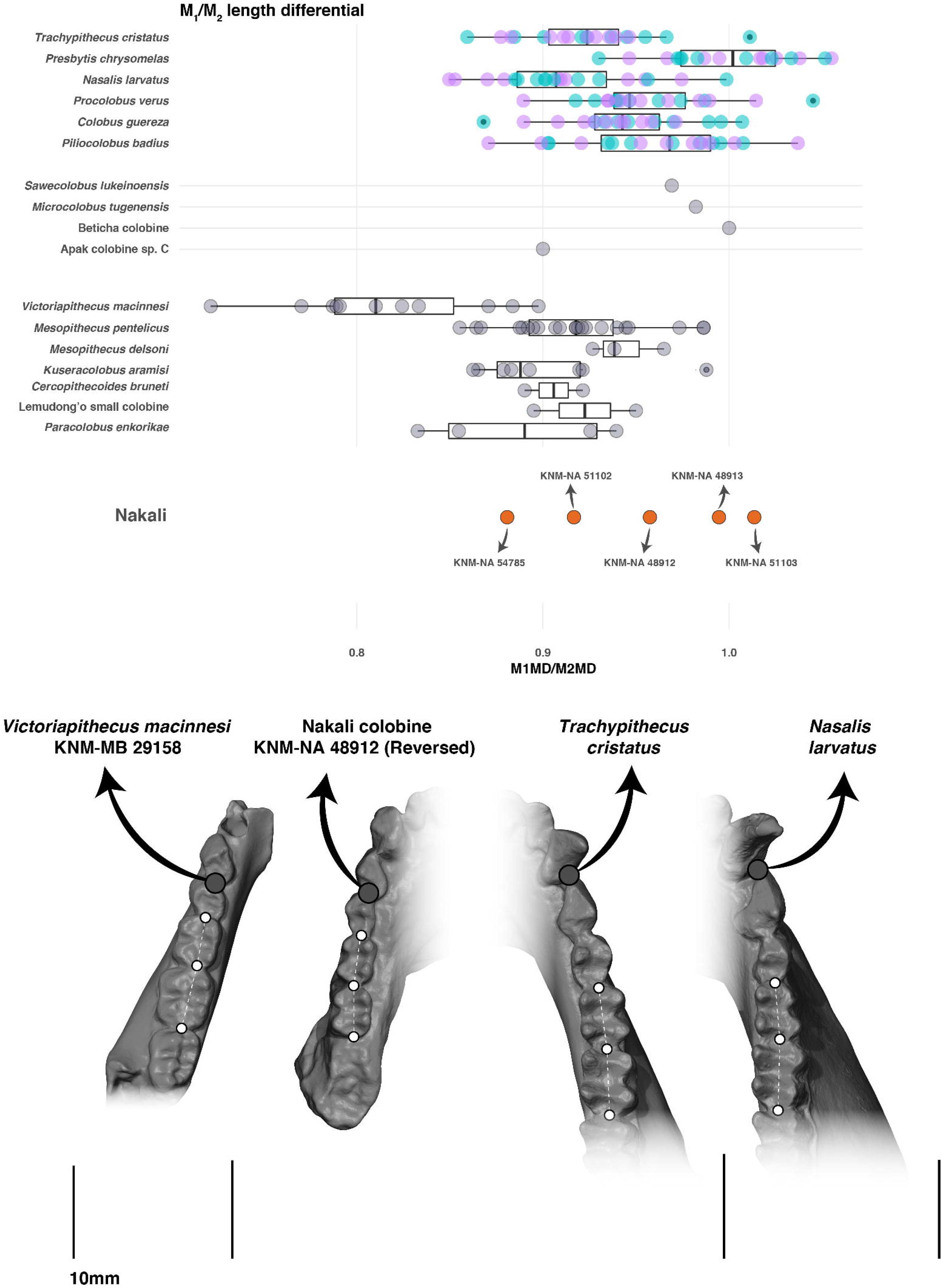
Boxplots of the length differential of M_1_ relative to M_2_ of extant and fossil colobines. Boxplots with first, third quartile, and median (black line). Fossil cercopithecids and extant colobines are shown on the bottom of the graph to compare them and to illustrate the subequal length of the M_1_ and M_2_ of *Tr. cristatus* and the marked length differential of the M_1_ and M_2_ of *N. larvatus*.

The index of the breadth differential of the M_2_ lophids permit to discriminate Presbytini from Colobini (Figure 16). The former present, on average, greater ratio values than Colobini, highlighting the relatively narrow M_2_ mesial lophid of Colobini. The Nakali sample shows index values most comparable to that of Colobini than Presbytini, specifically with the specimens KNM-NA 51102 and KNM-NA 48912 falling outside the range of variation or within the lower quartile of Presbytini taxa. In index values, the Nakali sample is similar to *Mesopithecus* spp., *K. aramisi* and *Pa. enkorikae*, but is quite distinct from the high index values of *V. macinnesi*. As a whole, the Nakali sample encompasses the range of variation of most fossil colobines, apart from the Lukeino M_2_ KNM-TH 36742 and *Ce. bruneti* TM 266 03-099, with slightly lower and higher index values, respectively.

**Figure 16:**
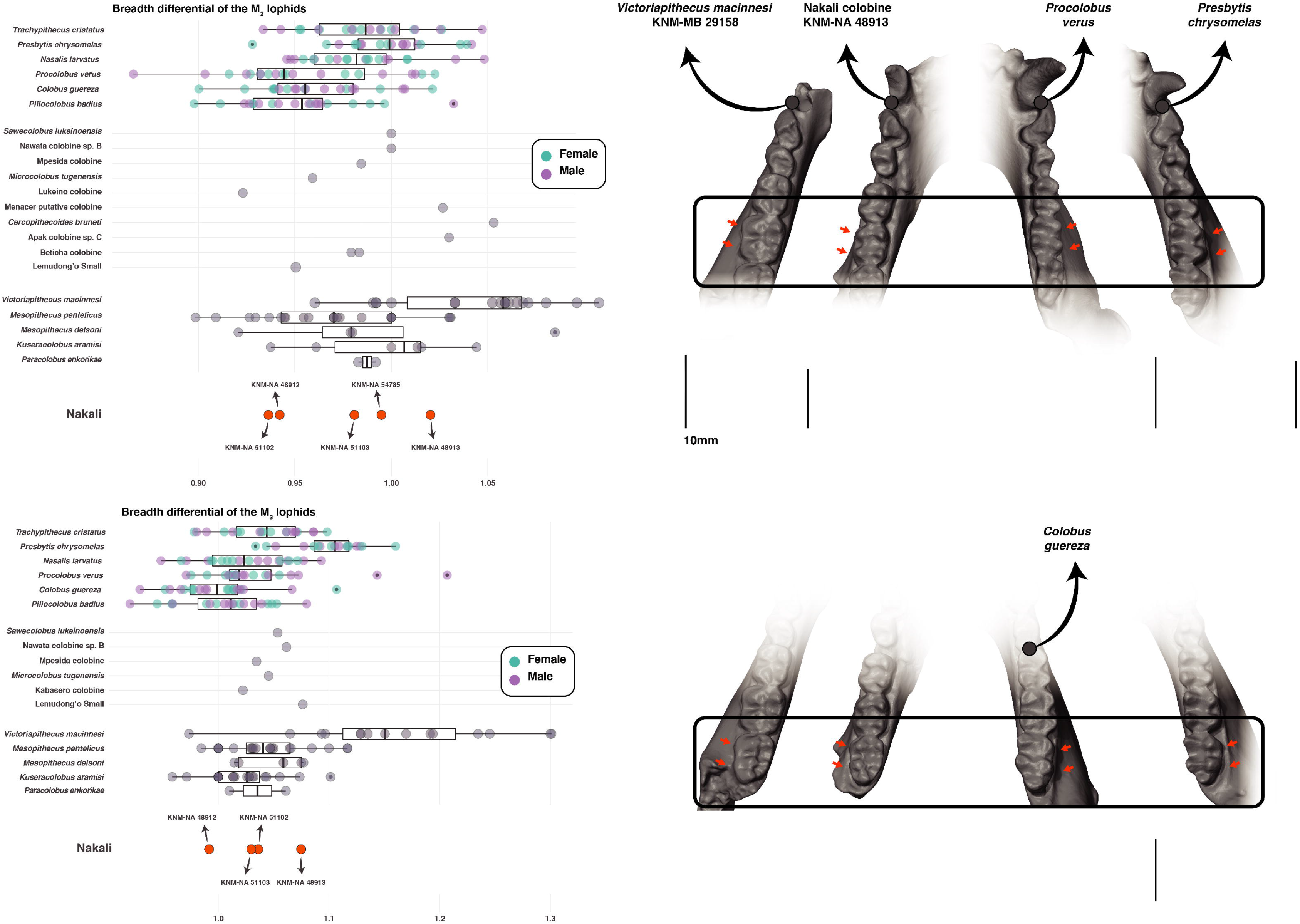
Boxplots of the breadth differential of the M_2_ lophids and M_3_ lophids of extant and fossil colobines. Boxplots with first, third quartile, and median (black line). Fossil cercopithecids and extant colobines are shown on the bottom of the graph to compare them and to illustrate the subequal breadth of the M_2_ and M_3_ of *Pre. chrysomelas,* the marked breadth differential of the M_2_ of *Pro. verus*, and the marked breadth differential of the M_3_ of *Co. guereza*.

The occlusal relief of the M_2_ (NC/NR in Benefit and Pickford, 1986 and NH / NR in Benefit, 1993) of KNM-NA 51103 is moderate, with a value of 108 (Online Resource 12), and is in the range of variation of extant colobines (µ = 143 ± 37.4; Benefit and Pickford, 1986:458) but outside that of extant cercopithecines (µ = 47 ± 17.9; Benefit and Pickford, 1986:458). The cusps height of KNM-NA 51103 is also slightly higher than that of *Me. pentelicus* NHMW 1998 0077 (NH / NR = 96; Online Resource 12), higher than the value reported for the isolated M_3_ from Kabasero KNM-TH 48368 (NH / NR = 75 in Rossie et al. 2013:5820), from that of *Mi. tugenensis* KNM-BN 1740 (NC / NR = 94 in Benefit and Pickford, 1986:458) but lower than the M_2_ from the Mpesida colobine KNM-TH 30975 (NH / NR = 130 in Gilbert et al. 2010:467) and that from the Lukeino colobine KNM-TH 36742 (NH / NR = 150 in Gilbert et al. 2010:467). Relative to extant colobines, KNM-NA 51103 is much closer in value to *Co. guereza* (µ = 122 ± 31.5 in Benefit and Pickford, 1986:458) than to *Pro. verus* (µ = 168 ± 38.2) and *Pi. badius* (µ = 157 ± 36.5 in Benefit and Pickford, 1986:458).

The index of the breadth differential of the M_3_ lophids of *Pre. chrysomelas* is higher than that of other extant colobines, illustrating a mesial lophid always larger than the distal one (Figure 16). A reverse pattern is seen in *Co. guereza*, with on average, a lower index value, and hence lophids of nearly equal width. The M_3_ lophid proportions of the Nakali sample is broadly comparable to that of most extant colobines (except *Pre. chrysomelas*), and to all Miocene colobines. Index values are consistent between the Nakali specimens and *Mi. tugenen s*(*i*K*s*NM-BN 1740), *S. lukeinoensis* (OCO 607’10), the Mpesida (Tugen Hills) M_3_ KNM-TH 23169, the putative Kabasero (Tugen Hills) M_3_ KNM-TH 48368, and the Nawata (Lothagam) sp. B colobine M_3_ KNM-LT 23083. However, the Nakali specimens are outside the interquartile range of *Vi. macinnesi*.

#### Variation in dental indices between the Nakali sample, penecontemporaneous fossil colobines, and extant colobines

The coefficient of variation (CV) of the P_4_ CSR of the Nakali sample (6.00; Table 2) is similar to that of extant colobine species (e.g., 5.99 in *Pre. chrysomelas*). When the isolated P_4_ KNM-BN 1251 and *Mi. tugenensis* KNM-BN 1740 are combined with the Nakali sample, the overall CV is slightly higher than that of the most variable colobine species considered here (6.85 vs. 6.43 in *Pi. badius*; Table 2). Bootstrap analysis failed to demonstrate a higher CV in Nakali than that of any extant colobine species (Figure 17). This result is valid when, independently, KNM-BN 1740 and KNM-BN 1251 are added to the Nakali sample. However, when KNM-BN 1740 and KNM-BN 1251 are added together to the Nakali sample, the variation exceeds that observed in *Procolobus* and *Trachypithecus*.

**Table 2:** Coefficient of variation of selected dental indices for extant colobines and different sample of fossil specimens.

|  | M<sub>1</sub><br>Geometric<br>mean | M<sub>2</sub><br>Geometric<br>mean | P<sub>4</sub> shape | P<sub>4</sub> area<br>relative<br>to M<sub>1</sub><br>area | M<sub>1</sub> length<br>relative<br>breadth of<br>the mesial<br>lophid | M<sub>1</sub><br>length<br>relative<br>to M<sub>2</sub><br>length | M<sub>1</sub> breadth<br>differential<br>of the<br>lophids | M<sub>2</sub> breadth<br>differential<br>of the<br>lophids | M<sub>3</sub> breadth<br>differential<br>of the<br>lophids |
| --- | --- | --- | --- | --- | --- | --- | --- | --- | --- |
| *Co. guereza* | 4.74 | 5.47 | 5.44 | 9.73 | 4.52 | 3.60 | 3.54 | 3.14 | 4.03 |
| *Pi. badius* | 2.80 | 3.18 | 6.43 | 7.75 | 5.29 | 4.53 | 3.24 | 3.32 | 4.07 |
| *Pro. verus* | 5.23 | 5.27 | 4.49 | 10.06 | 4.96 | 3.68 | 5.82 | 4.38 | 5.33 |
| *Pre.*<br>*chrysomelas* | 3.40 | 4.61 | 5.99 | 6.02 | 5.21 | 3.56 | 2.63 | 2.79 | 4.05 |
| *N. larvatus* | 4.17 | 4.40 | 5.51 | 10.72 | 4.20 | 4.28 | 3.21 | 2.82 | 3.75 |
| *Tr. cristatus* | 4.21 | 4.24 | 5.25 | 6.67 | 5.36 | 3.73 | 2.17 | 3.03 | 3.46 |
| Nakali | 4.64 | 8.72 | 6.00 | 6.15 | 3,75 | 5.26 | 3.26 | 4.12 | 2.91 |
|  | (including<br>KNM-NA<br>305) |  |  |  | (including<br>KNM-NA<br>305) |  | (including<br>KNM-NA<br>305) |  |  |
| Nakali +<br>Ngeringerowa<br>(KNM-BN 1740) | 4.67 | 8.23 | 6.85<br>(including<br>KNM-BN<br>1740 and<br>KNM-BN<br>1251) | 12.90 | 3.48 | 5.20 | 3.06 | 3.09 | 2.91 |
| Nakali + Beticha<br>(CHO-BT 78) | 4.81<br>(including<br>CHO-BT 78) | 7.50<br>(including<br>CHO-BT 78) | NA | NA | 5.45 | NA | 3.06 | NA | NA |

**Figure 17:**
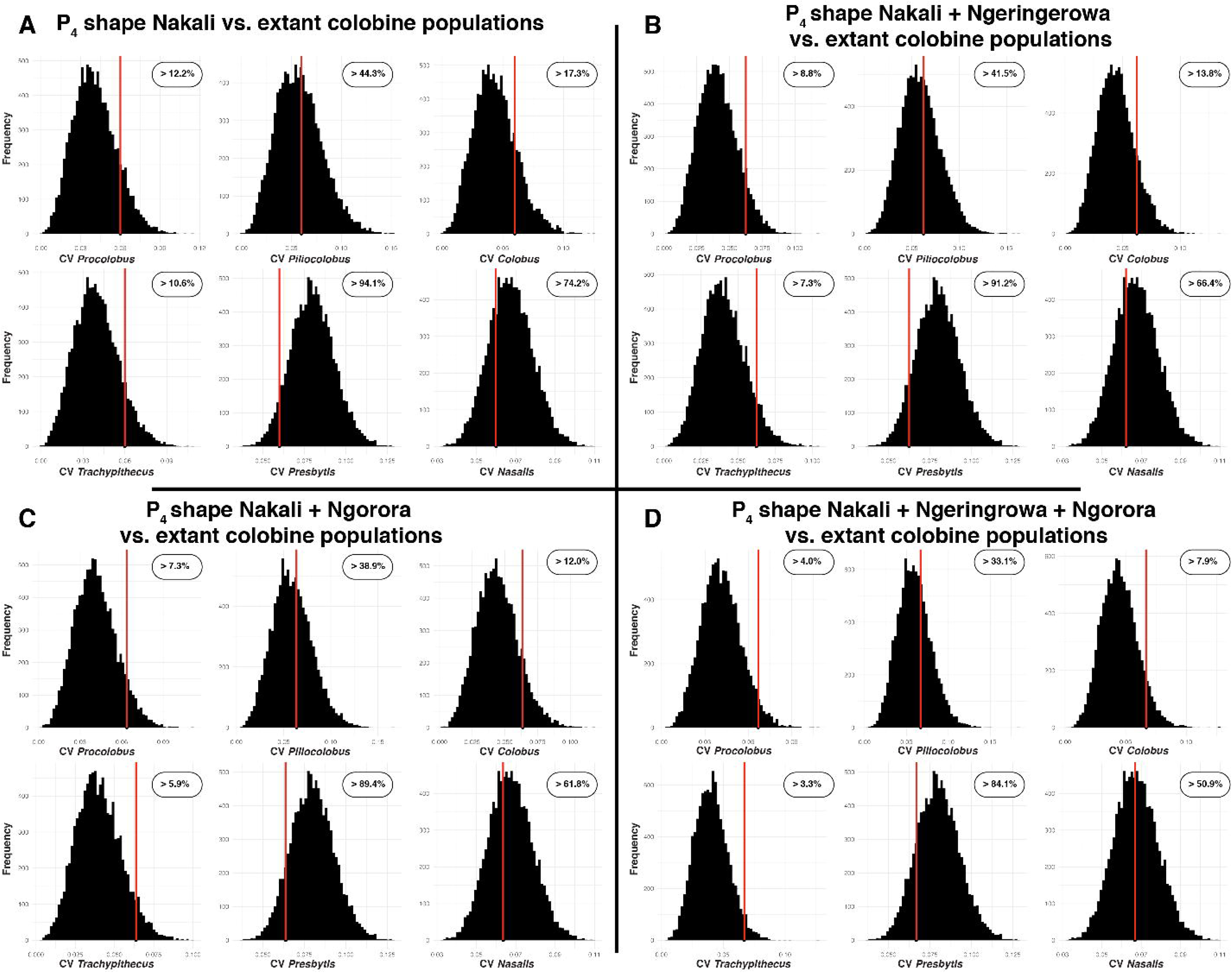
Frequency distribution of the coefficient of variation of the generated distributions of extant colobine species for the P_4_ shape index. Probability of observing a coefficient of variation higher than that of the considered fossil sample is shown on the upper right of each frequency distribution. The observed coefficient of variation for the considered fossil sample is indicated by a red vertical line. Panel A represent the coefficient of variation of the Nakali sample, panel B the combined Nakali and Ngerngerwa sample, panel C the combined Nakali and Ngorora sample, and panel D the combined Nakali, Ngerngerwa, and Ngorora sample.

The CV of the P_4_ area / M_1_ area ratio of the Nakali specimens is quite low (6.10; Table 2), comparable to that of *Pre. chrysomelas* (6.02) and does not exceed the one observed in other colobine species (e.g., 10.06 in *Pro. verus*). When KNM-BN 1740 is added to the Nakali sample, the overall CV is higher than that of any single extant colobine species (12.90 vs. 10.72 in *Na. larvatus*). When compared to the distribution obtained with the bootstrap analysis, the CV of the Nakali sample significantly exceeds that of *Pi. badius*, *Tr. cristatus*, and *Pre. chrysomelas* (Figure 18).

**Figure 18:**
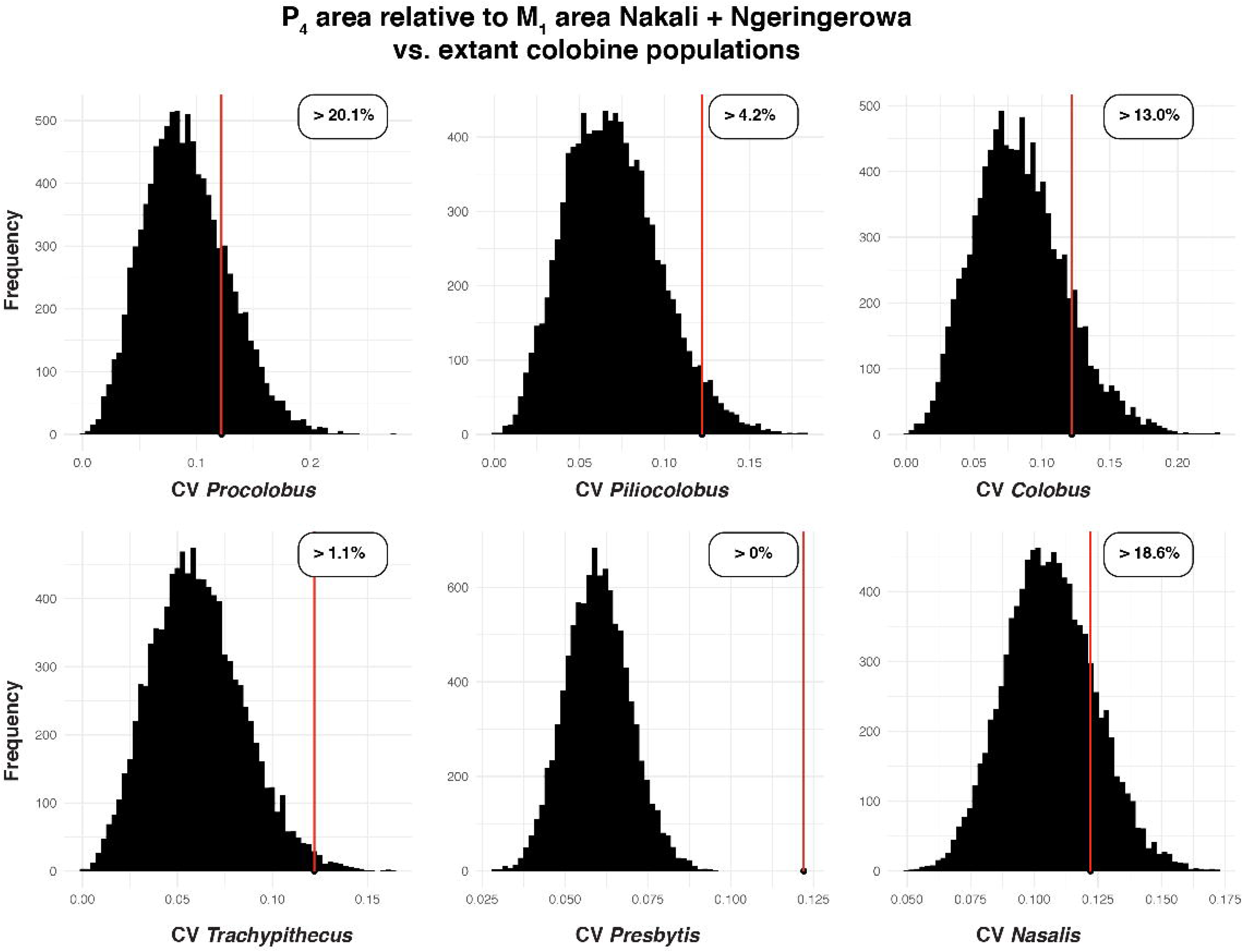
Frequency distribution of the coefficient of variation of the generated distributions of extant colobine species for the P_4_ area relative to M_1_ area index. Probability of observing a coefficient of variation higher than that of the considered fossil sample is shown on the upper right of each frequency distribution. The observed coefficient of variation for the considered fossil sample is indicated by a red vertical line.

The CV of the index of the breadth differential of the M_1_ lophids of the Nakali sample (including the isolated molar KNM-NA 305) is comparable to that of an extant colobines species (e.g., 3.26 vs. 3.21 in *Nasalis larvatus*; Table 2). A lower CV is observed when the possible *Microcolobus* specimen BT-78 from Beticha (Chorora) and *Mi. tugenensis* KNM-BN 1740 are added to the Nakali sample (3.06). However, considering the uncertainty as to which locus BT-78 represents (M_1_ or M_2_), we did not include it in the bootstrap analysis. When we added KNM-BN 1740 and KNM-NA 305 to the Nakali sample, we failed to demonstrate a higher CV in the fossil sample than in most extant colobine species, with the sole exception of the comparison of the fossil sample with *Tr. cristatus* (Online Resource 15).

The CV of the M_1_ length relative to its mesial lophid breadth of the Nakali sample (3.75, including KNM-NA 305; Table 2) is quite low compared to extant colobine species. We confirm this result through bootstrap analyses, where the combined Nakali and Ngerngerwa sample is less variable than most colobine species. It is even significantly less variable than the generated samples of *Pre. chrysomelas* and *N. larvatus* (Figure 19).

**Figure 19:**
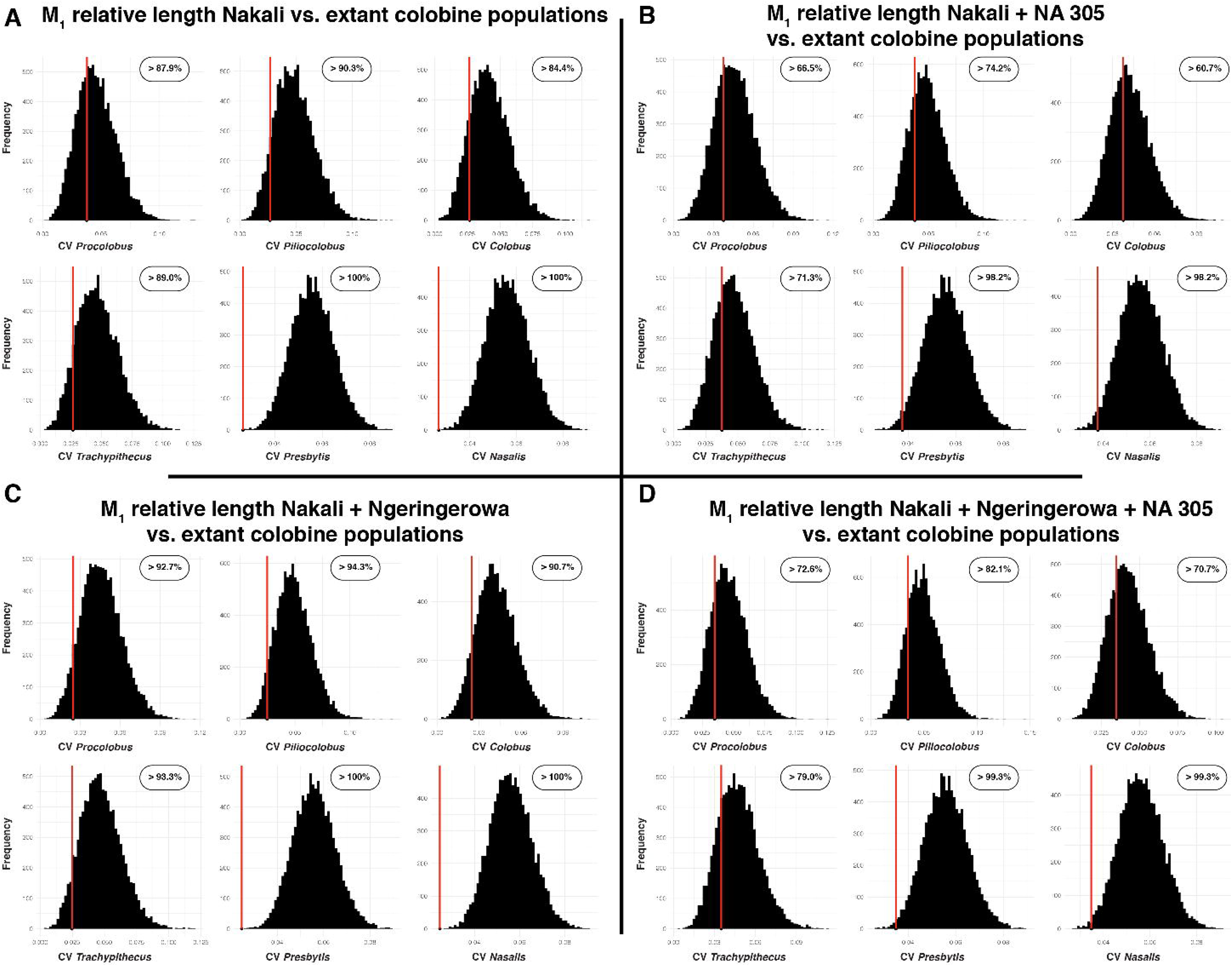
Frequency distribution of the coefficient of variation of the generated distributions of extant colobine species for index of the relative length of the M_1_. Probability of observing a coefficient of variation higher than that of the considered fossil sample is shown on the upper right of each frequency distribution. The observed coefficient of variation for the considered fossil sample is indicated by a red vertical line. Panel A represent the coefficient of variation of the Nakali sample, panel B the combined Nakali and KNM-NA 305 sample, panel C the combined Nakali and Ngerngerwa sample, and panel D the combined Nakali, Ngerngerwa, and KNM-NA 305 sample.

The CV of the M_1_/M_2_ length differential index of the Nakali sample (5.26; Table 2) is higher than that of any extant colobine species (4.53 in *Pi. badius*). However, the CV of the Nakali sample decreases when KNM-BN 1740 is added to it. Bootstrap analysis reveals that the Nakali sample is indeed highly variable in comparison with extant colobine species, since its CV significantly exceeds that of *Pro. verus*, *Pre*. *chrysomelas*, and *N*. *larvatus* (Online Resource 15).

The CV of the index of the breadth differential of the M_2_ lophids of the Nakali sample (4.12; Table 2) is consistent with that of extant colobine species (e.g., 4.38 in *Pro. verus*). A lower CV is observed when KNM-BN 1740 is added to the Nakali sample (3.09). However, bootstrap analyses yielded another result, with the Nakali sample exceeding the variation seen in *Presbytis* and *Nasalis* (Online Resource 15).

The CV of the index of the breadth differential of the M_3_ lophids of the Nakali sample (2.91; Table 2) is low compared to most extant colobine taxa, especially *Pro. verus* (5.33). Similarly, no bootstrap analysis of extant colobine species present a CV significantly higher than that of the Nakali sample (Online Resource 15).

#### Significant differences in dental ratio between extant colobines subfamilies and genera

Colobini and Presbytini (i.e., *Trachypithecus*, *Nasalis*, and *Presbytis*) can be significantly discriminated in intrinsic breadth differential of the M_1_, M_2_, and M_3_ lophids as well as in relative length of the C_1_ (Online Resource 16). Presbytini have M_1_ lophids of nearly equal width relative to Colobini. This is marked in *Tr. cristatus* among Presbytini genera which is significantly different from all Colobini genera and *N. larvatus*. Presbytini also have M_2_ lophids of nearly equal width compared to Colobini. This morphological pattern is well expressed in *Pre. chrysomelas*, which differs from *Pi. badius*, *Pro. verus*, and *N. larvatus*. Presbytini have a breadth differential of the M_3_ lophids distinct from that of Colobini, with a mesial lophid wider than the distal lophid. *Pre. chrysomelas* is significantly distinct from all other colobines in this dental ratio, presenting a mesial lophid significantly broader than the distal lophid. The P_4_ of Presbytini is broader than longer compared to that of the Colobini. Among Presbytini, the P_4_ of *Pre. chrysomelas* is significantly shorter and broader than that of *N. larvatus*. The C_1_ of Colobini is significantly more elongated relative to M_2_ GM than that of Presbytini. *Co*. *guereza* is significantly distinct from all the considered Presbytini taxa in this dental index.

Although non-significant differences between subfamilies are observed for the ratio of the P_4_ area relative to that of M_1_, we have found significant differences between extant colobine genera of the same subfamily (Online Resource 16). The area of the P_4_ relative to that of the M_1_ of *Co. guereza* is significantly distinct and higher from that of *Pi. badius* and *Pro. verus*. Similarly, *N. larvatus* present a relatively reduced area of the P_4_ compared to M_1_ area and a significantly shorter M_1_ relative to its M_2_ length compared to *Tr. cristatus* and *Pre. chrysomelas*.

#### PCA

For the sparse dataset, PC1 explains 31.9% of the variance, with positive scores on this axis mostly influenced by the length of the M_2_ and the breadth of the mesial lophid of the M_3_ (Figure 20A). Negative scores are mainly driven by the breadth of the mesial lophid of the M_1_ and the breadth of the P_4_. The Nakali specimens KNM-NA 48913, KNM-NA 48912 and KNM-NA 51102 show negative scores on PC1 while KNM-NA 54785 exhibits a slightly positive one. PC1 discriminates *Trachypithecus* and *Presbytis*, whose average scores are negative, from the other colobine taxa, whose scores are positive. PC2 accounts for 21.9% of the variance (Figure 20B). Negative scores on this axis are influenced by the breadth of the distal lophid of the M_2_ and M_1_. Positive scores on PC2 are induced by the length and breadth of the P_3_. All the Nakali specimens exhibit negative scores on PC2. The average scores of *Colobus* and *Trachypithecus* are positive and contrast with the negative scores of the other taxa. PC3 contributes to 13% of the variance, with positive scores on this axis explained by the length of the M_1_ and negative scores by the breadth of the M_2_. Apart from the slightly positive scores of KNM-NA 51102, all the Nakali specimens have negative scores on PC3. *Piliocolobus* and *Presbytis* have positive average scores on PC3, contrasting with the negative ones of *Trachypithecus* and *Colobus*. *Nasalis* and *Procolobus* have average PC3 scores clustering around zero. On both biplots, the average PC scores of the Nakali specimens fall near that of *Nasalis* and *Procolobus*.

**Figure 20:**
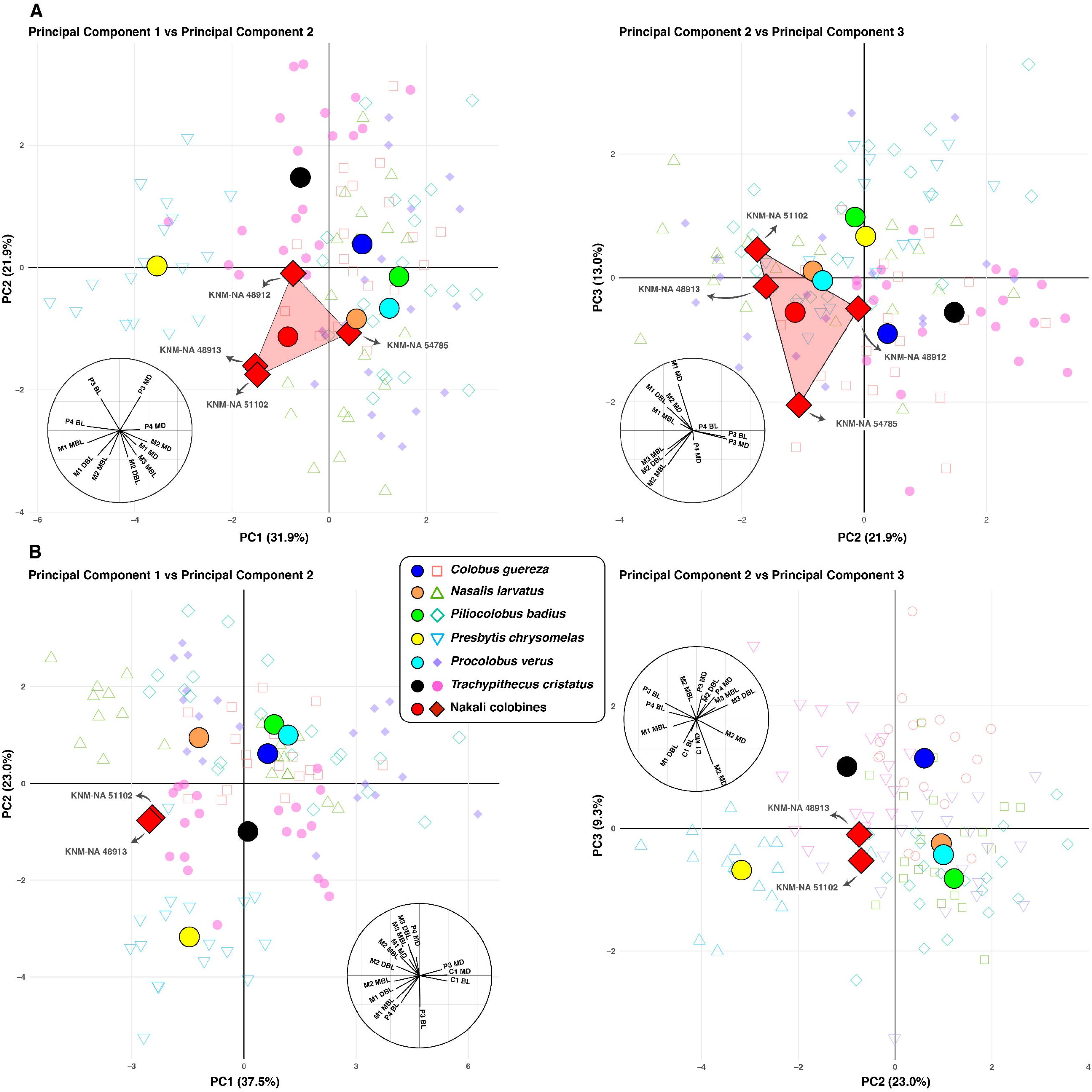
Biplot of PC1 and PC2 scores along with the biplot of PC2 against PC3 scores for the principal component analysis obtained using dental linear dimensions on the sparse dataset (panel A) and complete dataset (panel B). Correlation circles are visible on the lower left part of each biplot.

The complete dataset yielded result similar to the sparse dataset, also the inclusion of C_1_ MD and C_1_ BL permit to discriminate African colobines on PC1, which present, on average, positives scores on that axis. Contrary to Colobini, KNM-NA 48913 and KNM-NA 51102 have negative scores on PC1. M_3_ DBL also permit to distinguish colobines on PC2, with African colobines and *Nasalis* showing, on average, positive scores. KNM-NA 48913 and KNM-NA 51102 have slightly negative scores on PC2.

#### Linear Discriminant Analysis

All the considered Nakali specimens have LD1 scores that falls within the interquartile range of extant Colobini and in the lower quartile of extant Presbytini (Figure 21). They are also classified, with a high percentage of posterior probability (>80%), into Colobini. Precisely, KNM-NA 48913 is predicted to be a Colobini with an 88.8% posterior probability, KNM-NA 54785 with a 97.9% posterior probability, KNM-NA 51102 with a 90.4% posterior probability and KNM-NA 48912 with a 80.2% posterior probability.

**Figure 21:**
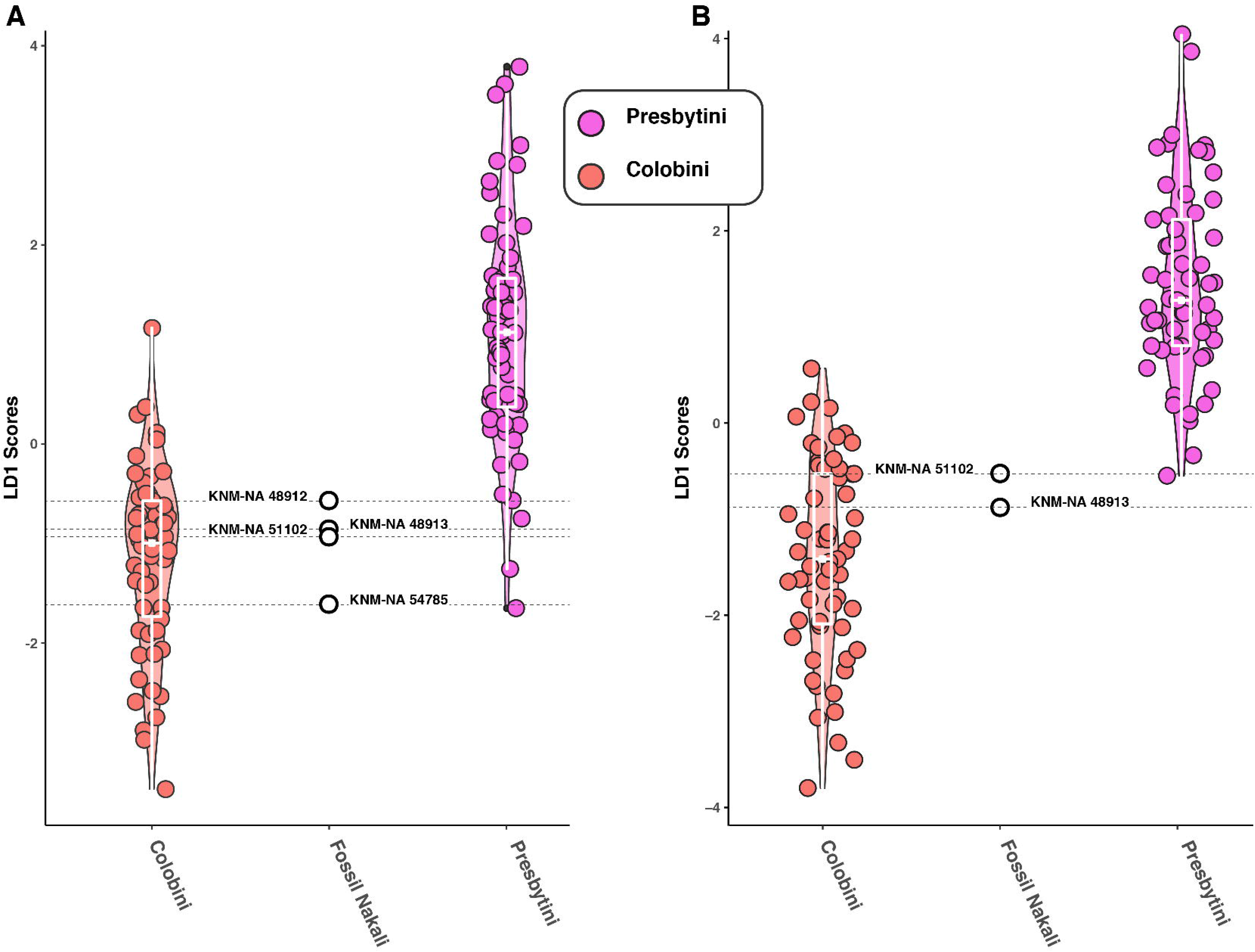
Boxplots superimposed on violin plots of LD1 scores of the linear discriminant analysis established on the PC scores of the dental principal component analysis of the sparse dataset (panel A) and complete dataset (panel B). Boxplots with first, third quartile, and median (black rectangle and horizontal line).

There are no differences in classification results between the complete and the sparse datasets. For the complete dataset, the posterior probability of belonging to a Colobini is of 93.3% for KNM-NA 48913 and 83.5% for KNM-NA 51102.

#### Qualitative dental traits

The P_4_ of KNM-NA 48913 is unworn and exhibits a distinct metaconid (lingual cusp), lower in height than the protoconid (buccal cusp) and distally offset from it (Figure 22). A similar morphology is observed in KNM-NA 48912, although the protoconid is slightly worn. An incipient metaconid, distally offset from the protoconid, is generally observed in specimens of *Colobus*. Indeed, 79% of the observed *Colobus* specimens from our sample have a distally offset metaconid (Figure 22). In contrast, the metaconid is extremely large, of nearly equal height and adjacent to the protoconid in *Nasalis* (Figure 22)*, Presbytis* and *Trachypithecus*. It is adjacent to the protoconid in 80% of the *Nasalis* specimens, 55.5% of the *Presbytis,* and 62% of the *Trachypithecus* (Figure 23). Similar to *Colobus*, *Piliocolobus* and *Procolobus* have a distally-offset metaconid (Figure 23), although its position is more variable in *Piliocolobus*, with 58% of specimens having a distally-offset metaconid, compared to 85% in *Procolobus*. The mesial shelf of the P_4_ is greatly enlarged in African colobines because of the distal positioning of the metaconid. This morphology is unlike that of *Presbytis* and *Nasalis*, which have reduced mesial shelves.

**Figure 22:**
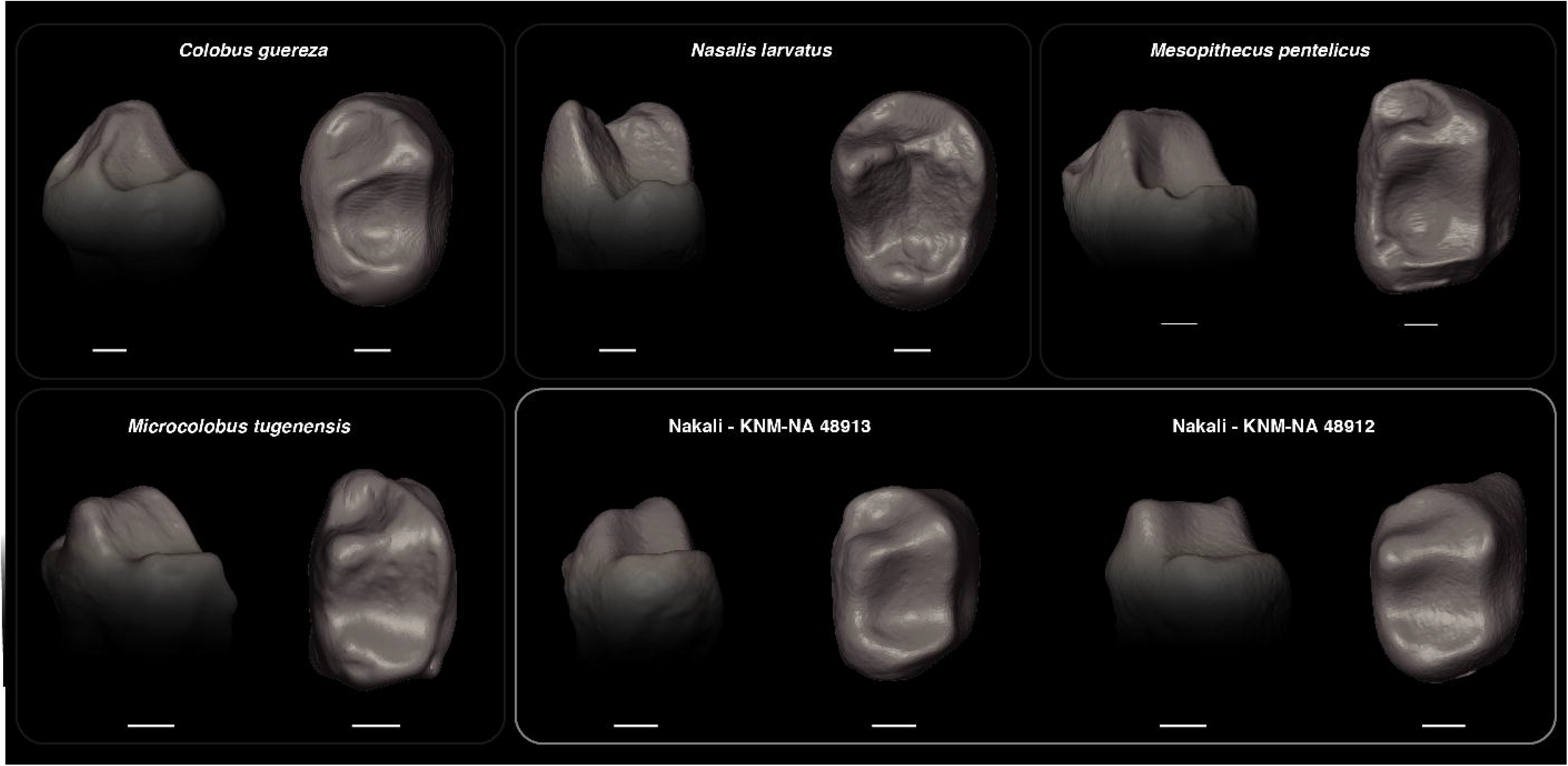
P_4_ occlusal morphology of extant and fossil colobines. Note the extremely reduced and distally-set metaconid of *Co. guereza* in comparison with *N. larvatus*.

**Figure 23:**
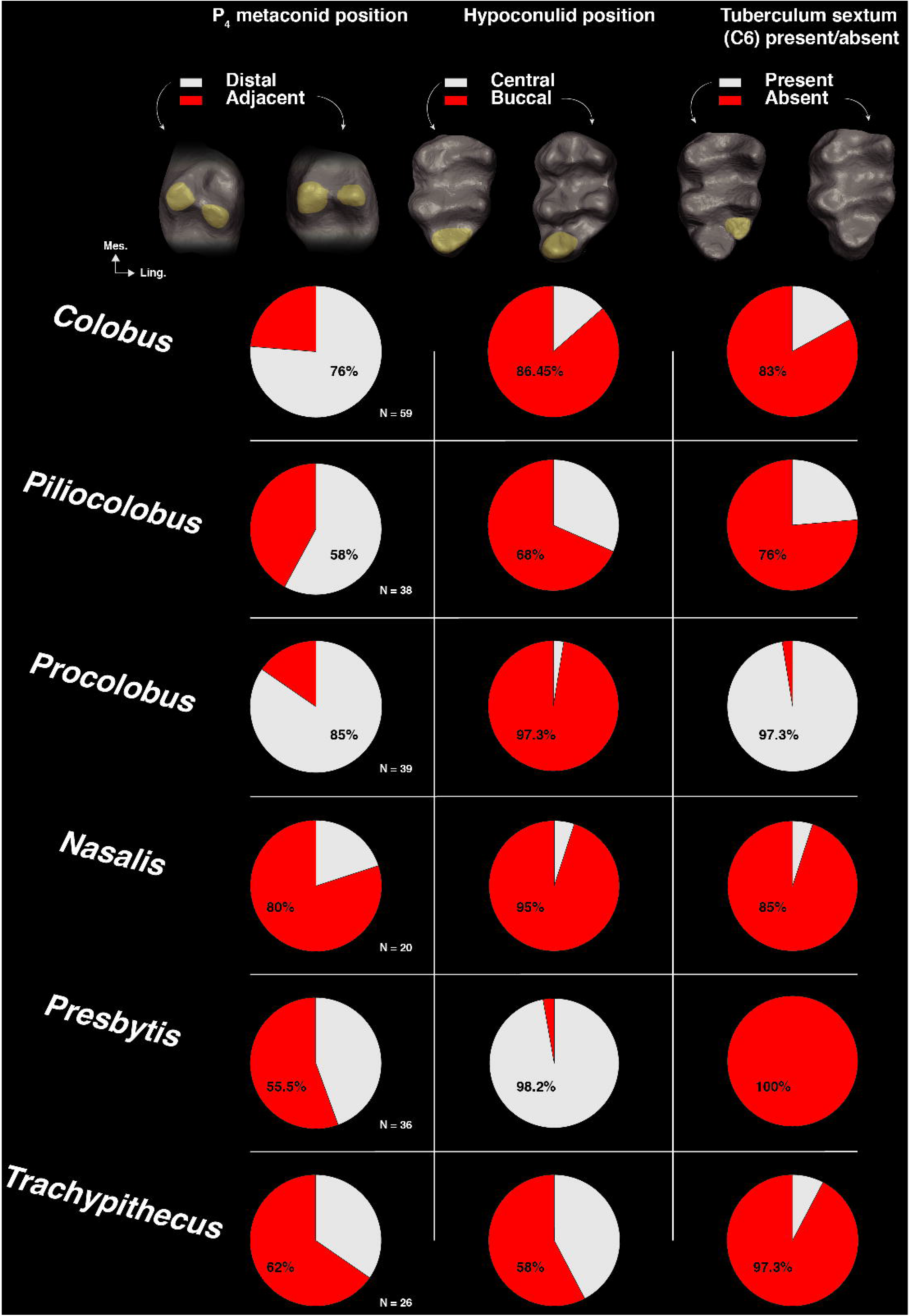
Pie plot of the frequency of observation of the position of the P4 metaconid (left panel), of the hypoconulid position (central panel), and of the presence / absence of the tuberculum sextum in extant colobines. An illustration of each character state is provided on top of the pie plots.

In comparison with fossil colobines, the mesial shelf of *Mi. tugenensis* (KNM-BN 1740) is quite large and limited to the lingual corner of the tooth, a morphology similar to that of the Nakali specimens and *Me. penteli c*(*u*Fi*s*gure 22). The large metaconid on the P_4_ of *N. larvatus* limit the dimension of the mesial shelf, contrary to the reduced or absent metaconid of *Co. guereza* and therefore its larger mesial shelf (Figure 22). *Microcolobus* is intermediate between *Co. guereza* and *N. larvatus* on this aspect. The P_4_ cusps of *Mi. tugenensis* are barely unworn and present a cusp development and a height differential similar to that of the Nakali specimens. A “reduced” metaconid “shorter than the protoconid” is also described by Benefit and Pickford (1986, p. 460) for the isolated Ngorora P_4_ KNM-BN 1251. A well-developed metaconid is frequently seen in *Me. pentelicus* and *Me. delsoni*, with only a moderate offset in height between both cusps (Delson, 1971; De Bonis et al., 1990). According to the *S. lukeinoensis* diagnosis of Gommery et al. (2022:476), the P_4_ metaconid and protoconid are ‘almost of the same height’ in this taxon. A height differential in favor of the protoconid and a distally offset metaconid is observed in *Ce. bruneti* (Pallas et al. 2019). A developed metaconid is visible in *Pa. enkorikae* and the small Lemudong’o colobine species. In both taxa, it is set distal to the protoconid. The metaconid of the Aramis sample of *K. aramisi* is described as “unreduced” by Frost and Hlusko (2008:144). In summary, the Nakali specimens conform to the general pattern presented in African colobines, that is the presence of a metaconid lower in height to the protoconid and set distally to it. None of the African fossil colobine studied here present the unreduced metaconid of *Co. guer e*, *z*n*a*or the high metaconid relative to protoconid observed in *Nasalis*.

The premolar morphology of *V. macinnesi* is quite diagnostic, with the documented presence of a metaconid on the P_3_ crown, a P_4_ metaconid higher than the protoconid, a triangular-shaped mesial fovea on the P_4_, and an oblique orientation of the P_4_ relative to the molar row long axis (Benefit, 1993). We observed no metaconid on the P_3_ of the Nakali specimens nor a developed P_4_ metaconid, characters that differentiate the Nakali colobine from *V. macinnesi*. In addition, the square-shaped P_4_ mesial fovea of the Nakali colobines also conforms to the pattern observed in extant colobines, but are unlike that of *V. macinnesi*.

The hypoconulid of the M_3_ is quite developed in KNM-NA 51103, KNM-NA 48913, and KNM-NA 48912. This is particularly evident when comparing the Nakali specimens with similar-sized specimens of *Presbytis*, which exhibits, when it is present, tiny hypoconulids. In this regard, the Nakali specimens are more similar to *Procolobus*, which, in spite of a small size, is presenting a well-developed hypoconulid. In *Procolobus*, the hypoconulid is accompanied by a tuberculum sextum (C6), which is less frequently seen in *Piliocolobus* and *Colobus*. In fact, while a C6 is seen in 97.3% of the *Pro. verus* specimens, it is absent in 76% of the *Piliocolobus* specimens, and 83% of the *Colobus* specimens (Figure 23). The presence of a C6 is also rare in *Trachypithecus* and *Nasalis*, where it is absent in 97.3% and 85% of the specimens, respectively (Figure 23). None of the Nakali specimens present a tuberculum sextum and conform to the general pattern of extant colobines. The position of the hypoconulid is also variable in extant colobines and, according to our observations, is mostly set in buccal position in Colobini. A typical case is *Pro. verus*, where a buccal hypoconulid is observed in 97.3% of the specimens (Figure 23). It is a frequency value comparable to that of *Na. larvatus* (95% in Figure 23). It is quite distinct from the small hypoconulid of *Presbytis*, which is located centrally (in 98.2% of the specimens), and the moderately developed hypoconulid of *Trachypithecus*, which is centrally placed in 42% of the specimens (Figure 23). The hypoconulid is buccally placed in all the Nakali specimens that preserve an M_3_, hence conforming to the Colobini pattern.

By exhibiting a well-developed, buccally-placed hypoconulid with no C6, the Nakali specimens are comparable in morphology to *Pa. enkorikae*, the small colobine from Lemudong’o and *Ce. bruneti*. The hypoconulid of *Me. pentelicus* is also well-developed and devoid of C6. However, it can also be observed in central position (e.g., BSM AS II 15 in Online Resource 17). The hypoconulid of *Mi. tugenensis* is slightly smaller than that of the Nakali specimens, but, qualitatively, the magnitude of difference between both samples can be encompass within the range of variation of one species (see *Co. polykomos* in Online Resource 17. It also shares with the Nakali specimens the absence of a C6 and a buccal position of the hypoconulid on the tooth.

### 3.3 MANDIBULAR MORPHOMETRY AND COMPARATIVE MORPHOLOGY

#### Univariate data

KNM-NA 54785 and KNM-NA 48913 show a breadth differential of the transverse tori superior to 1, illustrating a STT superior to the ITT (Figure 24). Both fossil specimens are in the interquartile range of variation of *Procolobus* and *Presbytis*, in the lower quartile of *Colobus*, and upper quartile of *Piliocolobus*. By presenting a moderately developed ITT, and hence a high ratio, KNM-NA 54785 and KNM-NA 48913 conform to the general pattern observed in fossil colobines, where only specimens of *Ce. bruneti* and *Me. pentelicus* present an ITT superior or equal in breadth to the STT. The relative proportion of the mandibular tori of the Nakali colobine is similar to that of *Noropithecus bulukensis* but distinct from that of *V. macinnesi*.

**Figure 24:**
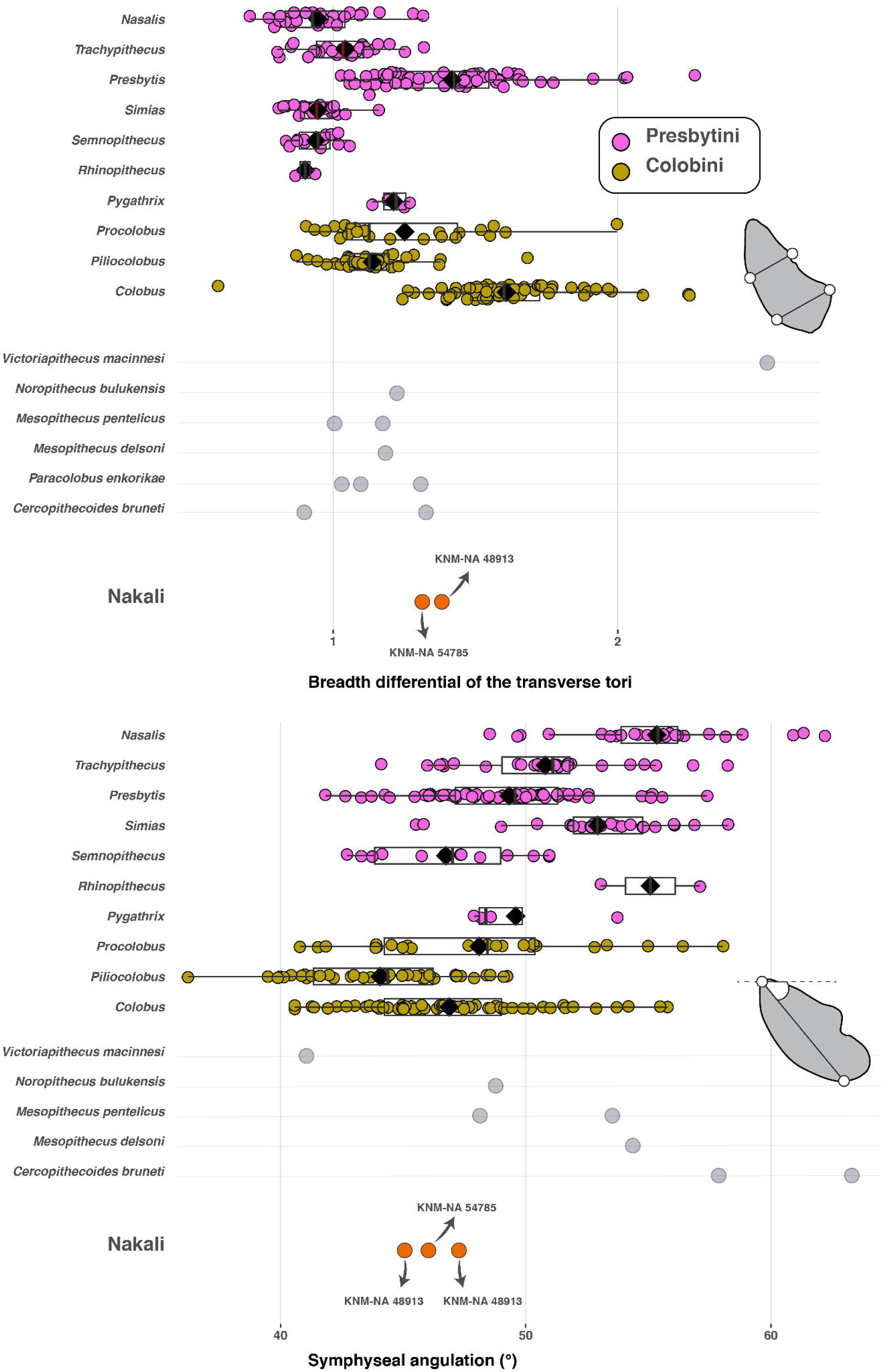
Boxplots of the breadth differential of the transverse tori (upper panel) and of the symphyseal angulation of extant and fossil colobines. Boxplots with first, third quartile, median (black line), and mean (black diamond). An illustration of the measurement protocol is provided on the right part of each panel.

The symphysis of the specimens KNM-NA 48913, KNM-NA 54785 and KNM-NA 48912 is acutely angled compared to most Asian colobines and present values in the interquartile range of variation of *Colobus* and *Procolobus* (Figure 24). The acutely angled symphysis of the Nakali colobines is distinct from the obtusely angled symphyseal profile of *Ce. bruneti* and *Me. delsoni*, and is slightly more acutely angled than that of *N. bulukensis* and *Me. pentelicus*. The symphyseal profile of *V. macinnesi* is acutely angled (ca. 40°), and does not match in absolute values that of the Nakali specimens.

#### Multivariate data

PC1 explains 41.3% of the variance, with positive scores associated with short and robust corpus and negative scores, deep and narrow corpus (Figure 25). KNM-NA 48912 exhibits a slightly negative score on the PC1 axis and falls near that of *Mi. tugenensis* KNM-BN 1740 and to the mean score of *Presbytis*. With negative scores, *N. bulukensis* and *Pa. enkorikae* present corpus slightly narrower than that of *Microcolobus* KNM-NA 48912 and KNM-BN 1740, but the magnitude of difference between those specimens does not exceed the range of variation of extant colobine species. KNM-NA 48912 and KNM-BN 1740 also fall in the range of variation of *Me. pentelicus* on PC1 but are outside that of *V. macinnesi*. Indeed, the corpus of *V. macinnesi* is extremely robust, with outlying values on PC1 for KNM-MB 18993.

**Figure 25:**
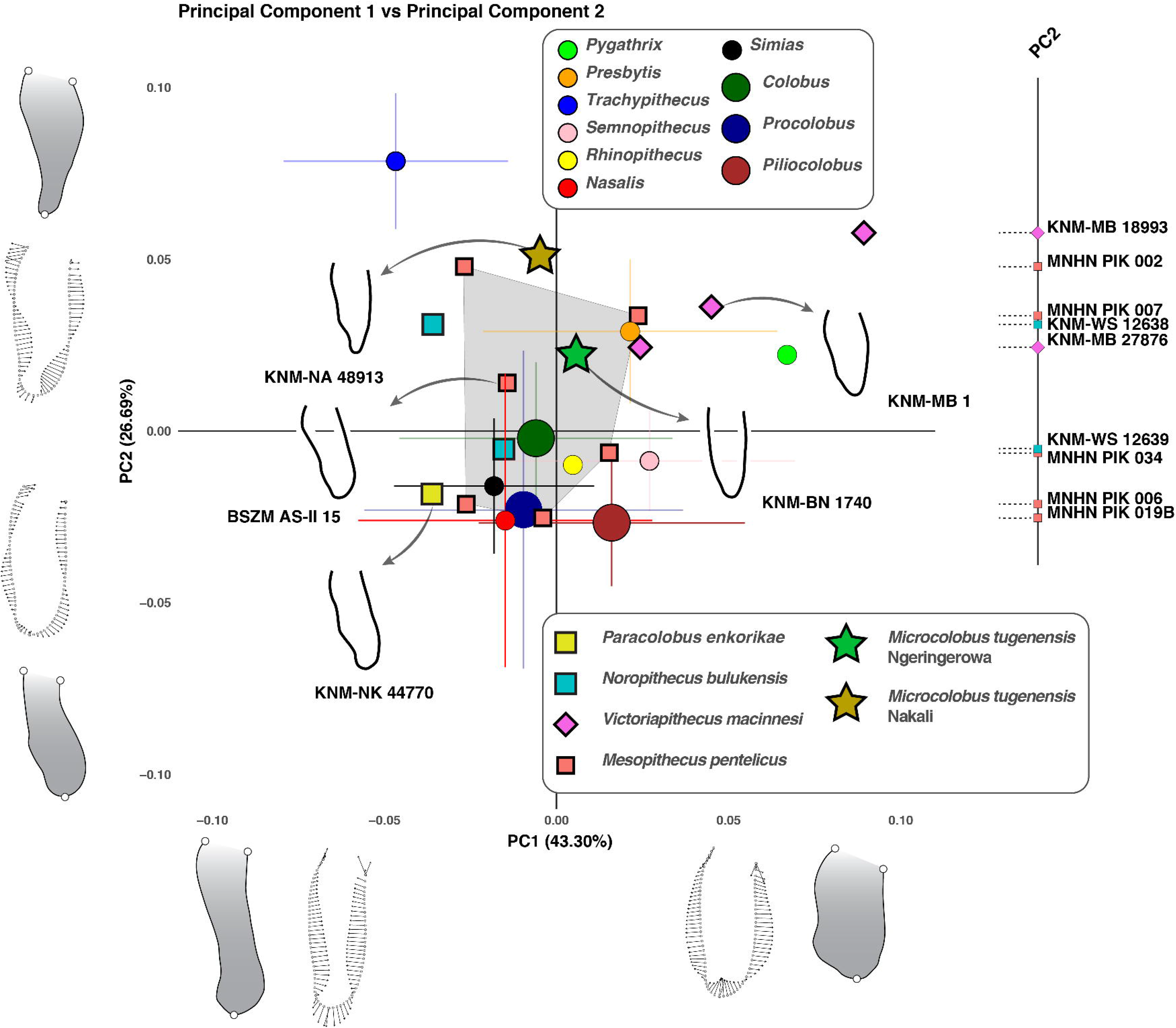
Biplot of the PC1 and PC2 scores of the Procrustes coordinates of the geometric morphometric analysis of the corpus cross-section at the M_1_-M_2_ junction of extant and fossil colobines. For each extant species, the mean (circles) is plotted along the standard deviation (horizontal and vertical bars). The outline of the cross-section of the corpus of selected fossil specimen is illustrated directly on the graph. The theoretical vector of morphological changes relative to the mean shape for minimum and maximum PC scores is visible next to each PC axis along with the minimum and maximum realized morphology. The projection of the fossil specimens on PC2 scores, along with their accession number, is provided to identified individually on the biplot.

PC2 accounts for 26.7% of the variance and positive scores on that axis are linked to distally thin corpus, with large submandibular fossae and weakly pronounced lateral prominences. Negative scores illustrate robust distal corpus and large lateral prominences (Figure 25). KNM-NA 48912 and KNM-BN 1740 share negative scores on PC2, illustrating a distally thin corpus with weakly expressed lateral prominences. As for PC1, KNM-NA 48912 and KNM-BN 1740 fall close to the mean PC2 score of *Presbytis*. Both specimens are also outside the range of variation of Colobini. The range of variation of the PC2 scores of *Me. pentelicus* encompasses KNM-NA 48912 and KNM-BN 1740, with half of the *Me. pentelicus* specimens showing negative scores (MNHN PIK 034, MNHN PIK 006, and MNHN PIK 019B). *Pa. enkorikae*KNM-NK 44770 is distinct from *Microcolobus* by exhibiting a negative PC2 score and hence developed lateral prominences. The positive PC2 scores of *Microcolobus* are similar to that *V. macinnesi* and *N. bulukensis*.

#### Qualitative mandibular traits

Qualitatively, the most inferior aspect of the ITT (attachment site of the anterior belly of the digastric) is slightly angled in the Nakali specimens and is comparable in shape to *Pa. enkorikae* (e.g., KNM-NK 42276 in Figure 26). The inferior aspect of the ITT is not as flat and as acutely-angled as in *N. bulukensis*, *Me. delsoni*, *Me. pentelicus*, and male *Ce. bruneti*. While *V. macinnesi* KNM-MB 18929 and KNM-MB 18993 have a weakly pronounced inferior aspect of the ITT, that of *V. macinnesi* KNM-MB 31283 is comparable to that of extant and fossil colobines (Figure 26). Among the Nakali colobines, the labial aspect of the symphysis is straight (KNM-NA 48913) to slightly convex (KNM-NA 54785), encompassing the range of variation seen in fossil colobines. Straight labial face of the symphysis is notably observed in *N. bulukensis*, *Me. delsoni*, *Me. pentelicus*, and *Ce. bruneti*, while a slightly more convex labial profile is seen in KNM-NK 42276.

**Figure 26:**
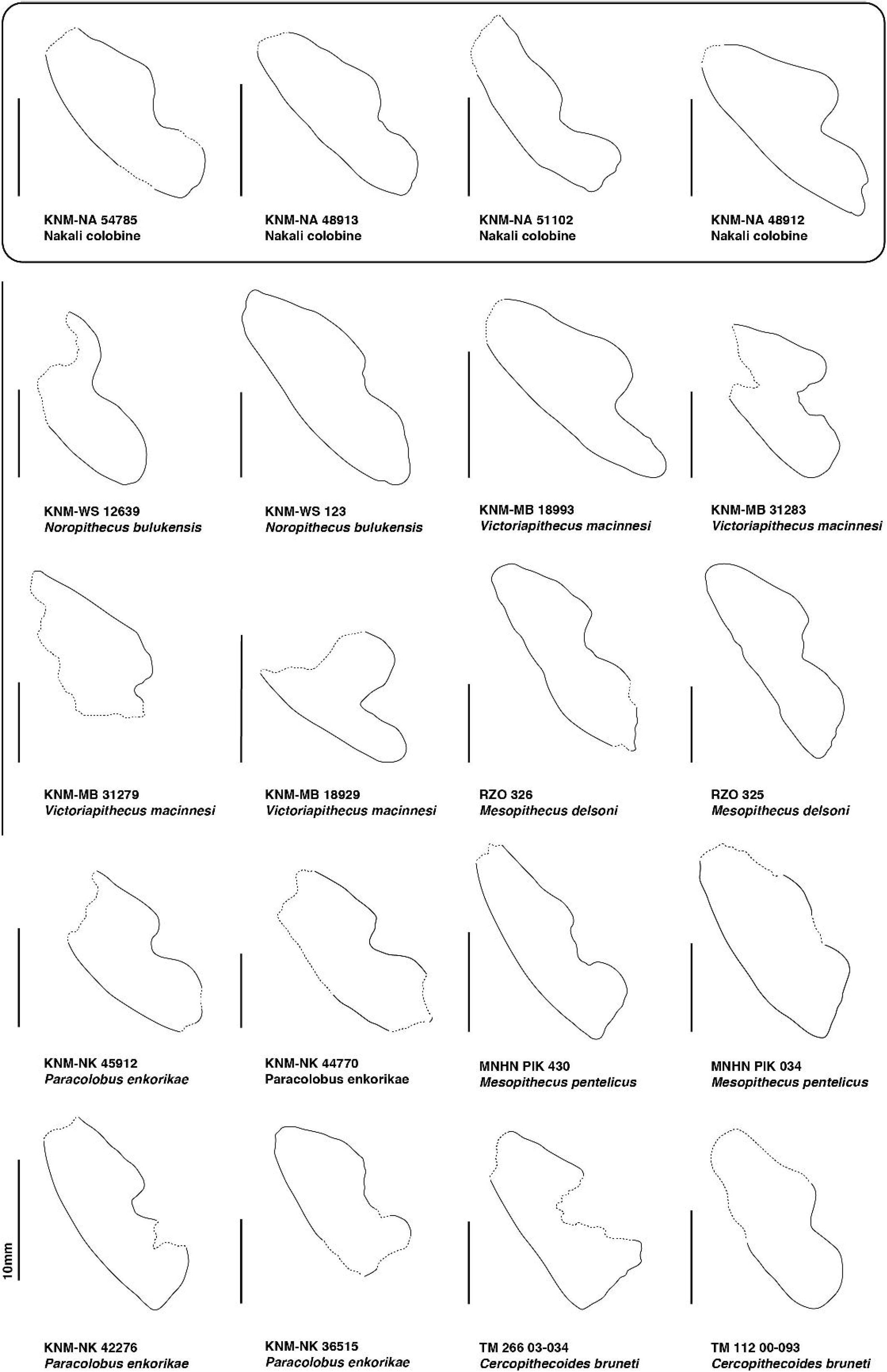
Outline of the symphyseal anatomy of fossil colobines obtained with transverse cross-section of 3D generated surface. Dotted line represents damaged bone and /or extrapolated anatomy while solid line represents reliable anatomical outline.

The planum alveolare of the Nakali *Microcolobus* specimens is flat, and is similar to most extant colobines. Variation in this trait is notably seen in *Pa. enkorikae*, with specimens showing flat (KNM-NK 42276) or slightly convex planum (KNM-NK 36515).

Irrespective of sexes, the corpus of *Microcolobus* from Nakali and Ngerngerwa have weakly-developed lateral prominences (Figure 27). *Microcolobus* is similar in this aspect to the small colobine from Lemudong’o (KNM-NK 36514), *Me. delsoni*, and female specimens of *Me. pentelicus*. The large lateral prominences of *Pa. enkorikae*, *Ce. bruneti*, and male *Me. pentelicus*differentiated them from *Microcolobus* (Figure 27). *Microcolobus* does not present the marked lateral prominences of *N. bulukensis* KNM-WS 12639 and is most similar to *V. macinnesi* (Figure 28).

**Figure 27:**
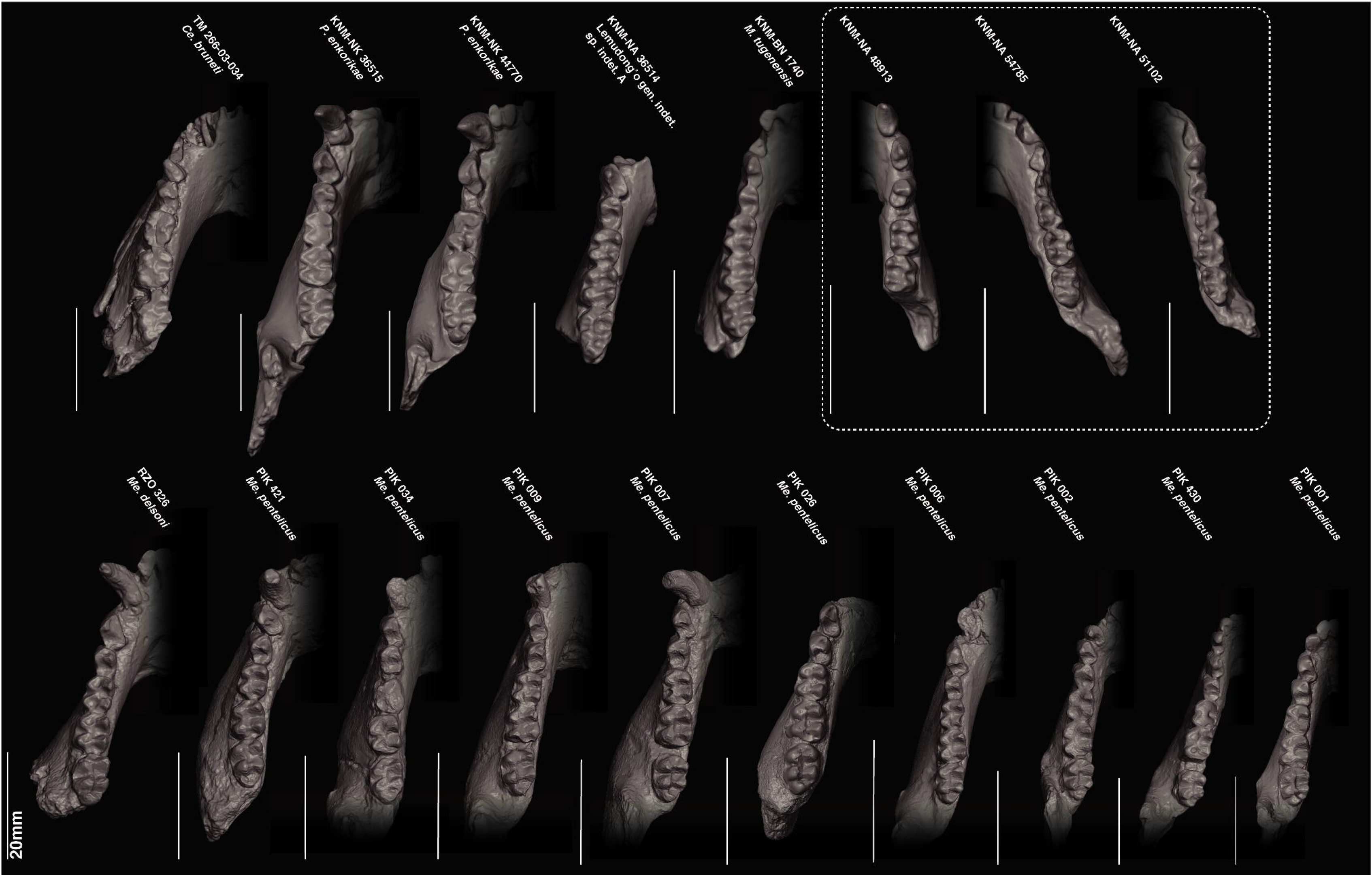
3D generated surface of the mandibular morphology of fossil colobines in occlusal view.

**Figure 28:**
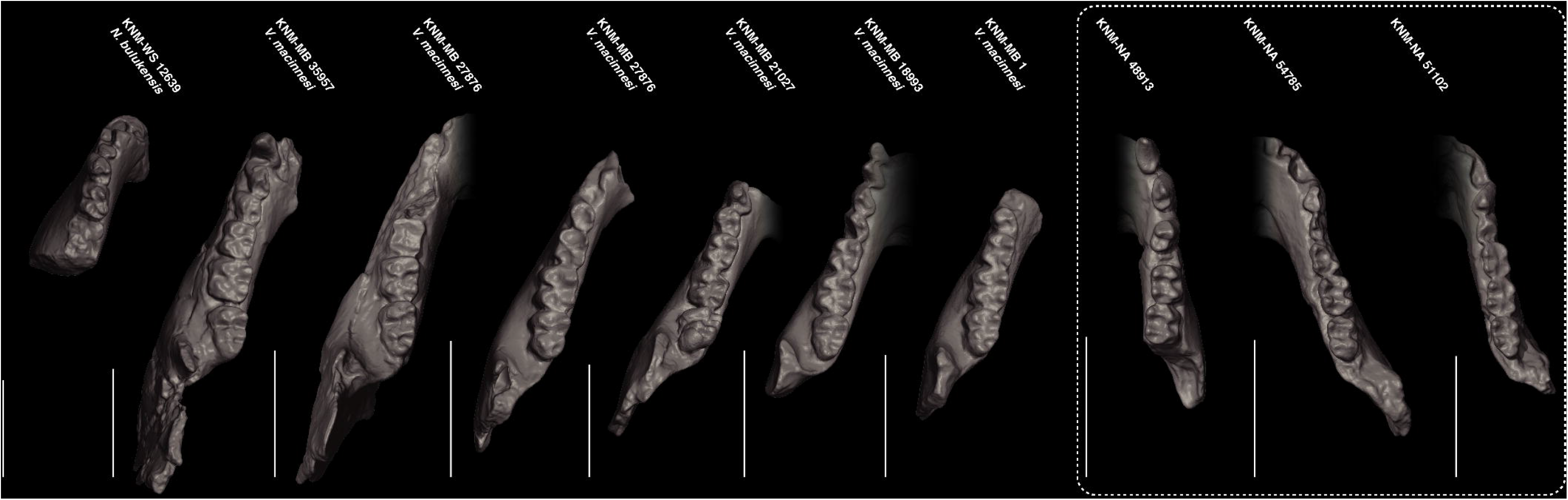
3D generated surface of the mandibular morphology of fossil colobines and victoriapithecids in occlusal view.

The lingual aspect of the corpus of KNM-NA 48913 and KNM-NA 54785 shows marked, but restricted in height, submandibular fossae, and a slight intertoral sulcus (Figure 29). This morphology is also seen in *Pa. enkorikae* KNM-NK 44770 but not in others fossil colobines. Qualitatively, the straight orientation and distal thinning of the corpus of *Microcolobus* is most comparable to *N. bulukensis* and *V. macinnesi*.

**Figure 29:**
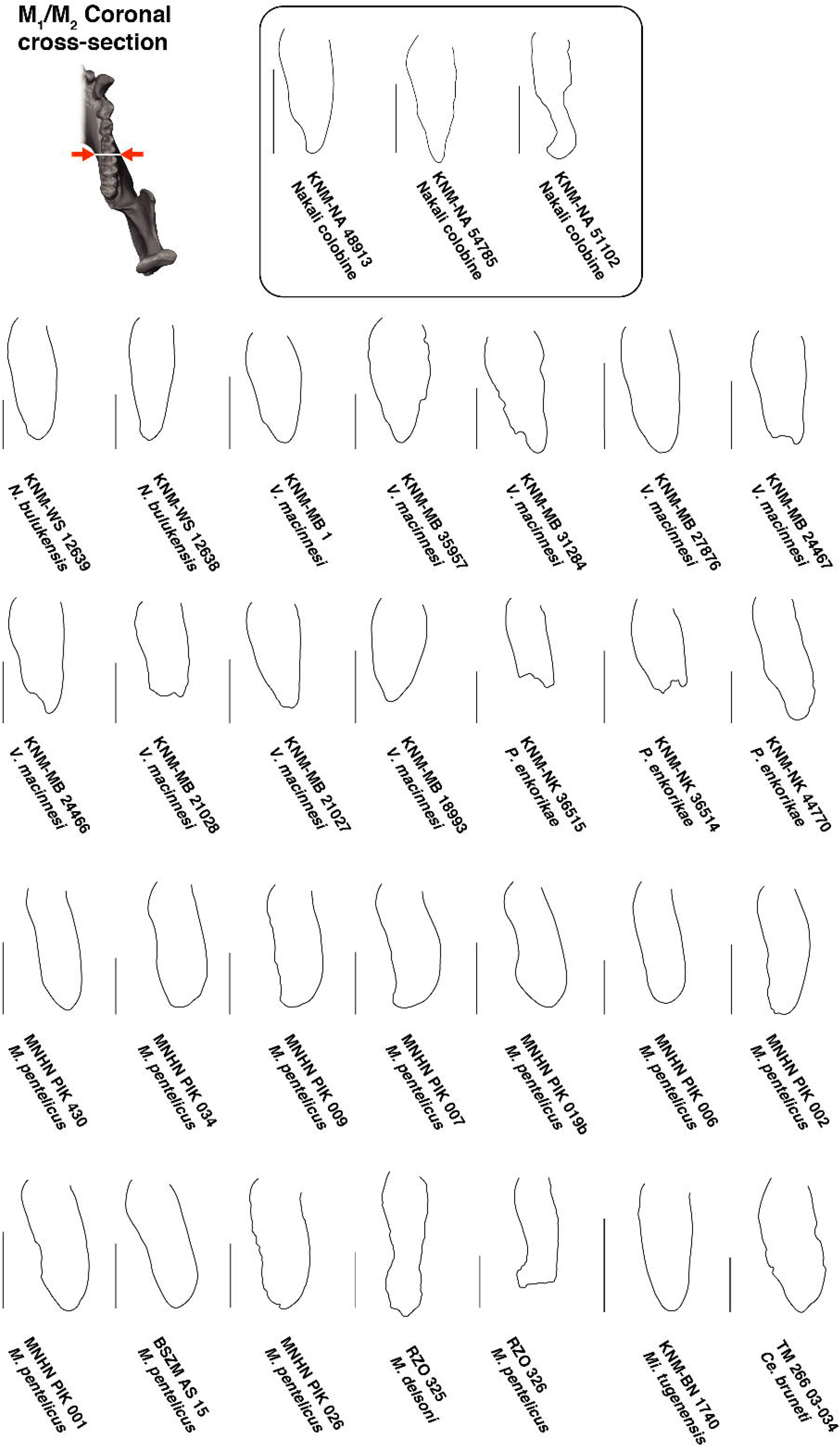
Outline of the corpus anatomy of fossil colobines obtained with transverse cross-section of 3D generated surface set at the M_1_-M_2_ junction.

### 3.4 A SUMMARY OF DENTAL AND MANDIBULAR EVIDENCE

A summary of the dental and mandibular evidence used to distinguish or relate the Nakali colobine specimens to fossil and extant colobines is provided in Table 3 (see below).

## DISCUSSION

We described six new mandibular specimens discovered from the Nakali Formation and provided new data on the dental, mandibular corpus and symphysis anatomy of Miocene and extant colobines. We have shown that the relative premolar and molar dimensions of four of these specimens are more similar to Colobini than to Presbytini. While the occlusal morphology of the P_4_ of the Nakali colobines corresponds to the anatomy of extant Colobini, the relative mesiodistal dimension of its C_1_ is different from that of extant Colobini. Similarly, while the breadth proportion of the transverse symphyseal tori of the Nakali colobine is within the range of variation of extant Colobini, its corpus shape is distinct from it and is most similar to *Presbytis* and victoriapithecids. At a taxonomic level, we failed to demonstrate unambiguous evidence of taxonomic diversity within the Nakali sample, and between the samples from Nakali, Ngerngerwa (KNM-BN 1740, type specimen of *Mi. tugenensis*), and Ngorora (KNM-BN 1251). Based on the available evidence, we attributed the Nakali sample to *Microcolobus tugenensis*.

### H0_1_ The newly discovered Nakali specimens are consistent with a single colobine species

According to the bootstrap analysis, the Nakali sample exceeds the CV of extant colobine species for only one of the six dental indices investigated. Indeed, while the M1 MD / M2 MD ratio is high, bootstrap analysis rejected the possibility that the variation in the Nakali sample exceeds that of all the colobine genera considered. The CV of the geometric size of the M_1_ of the Nakali fossil is also in accordance with that of extant colobine species, ruling out the presence of distinct species on the basis of dental size. Given the paucity of significant results regarding the comparison of CV between extant colobine species and the fossil sample from Nakali, we accept the hypothesis the Nakali colobine mandibles in this study belong to a single species.

### H0_2_ The newly discovered Nakali specimens are distinct from the penecontemporaneous colobines of Ngerngerwa and Ngorora

Benefit and Pickford (1986) suggested that the isolated M_1_ KNM-NA 305 was distinct from the type specimen of *Mi. tugenensis* from Ngerngerwa (KNM-BN 1740), mainly on the basis of the relative dimensions of the crown. Indeed, the M_1_ KNM-NA 305 is described as longer and narrower than that of KNM-BN 1740. Benefit and Pickford (1986) rationale is based on a shape index (M_1_ MD / M_1_ mesial lophid breadth), which is higher in KNM-NA 305 (1.33) than in KNM-BN 1740 (1.25). However, when the CV of the total fossil sample (i.e., Ngerngerwa and Nakali combined) is computed, it does not exceed that of extant colobine species and is even extremely small compared to them. This result is also statistically confirmed by bootstrap analyses where none of the generated distributions of extant colobine species exceed that of the combined sample from Nakali (i.e., including KNM-NA 305) and Ngerngerwa. This result does not support the hypothesis of Benefit and Pickford (1986), based on M_1_ crown shape, that the Nakali and Ngerngerwa fossil colobines were distinct taxa.

In terms of taxonomic affinity, the Nakali specimens are similar in M_1_ geometric size to KNM-BN 1740. Nakali specimens are also similar to *Mi. tugenensis* in four out of six dental indices. However, the shape index of the P_4_ of KNM-BN 1740 is slightly higher than that of the Nakali specimens and its P_4_ area relative to that of M_1_ is clearly outside the range of variation of the Nakali sample. Indeed, when KNM-BN 1740 is added to the Nakali sample, the CV of this ratio dramatically increases from 6.20 to 12.90, a value exceeding that of any of the extant colobine species considered here. However, bootstrap analyses did not fully demonstrate the significance of this result. While the combined sample of Nakali and Ngerngerwa significantly exceeds the coefficient of variation of the generated distribution of *Pi. badius*, *Tr. cristatus*, and *Pre. chrysomelas*, it does not exceed that of *Pro. verus*, *Co. guereza* and *N. larvatus*. Despite a relatively enlarged M_1_ area compared to the P_4_ area in *Mi. tugenensis* from Ngerngerwa, none of the continuous dental traits examined clearly discriminate KNM-BN 1740 from the specimens of Nakali. In addition, both samples share a similar morphology of the P_4_ cusps, with a metaconid smaller in size and set distal to the protoconid. With regard to mandibular morphology, apart from a more developed intertoral sulcus, which may reflect a greater development of the superior transverse torus, and/or, a more developed submandibular fossa in KNM-NA 48913, we did not detect major morphological differences between KNM-NA 48913 and KNM-BN 1740. This observation is firmly asserted quantitatively by a morphometric geometric analysis of the contour of the corpus, reflecting a shared morphology between both specimens characterized by a distal thinning of the corpus and weakly developed lateral prominences. The inferior transverse torus of *Mi. tugenensis* KNM-BN 1740 was described as highly reduced by Benefit and Pickford (1986), but newly acquired CT data demonstrates that this area is damaged in KNM-BN 1740, and hence, cannot be reliably assessed. We confirm here that the breadth of the inferior transverse torus of *Microcolobus* is well developed and is in the range of variation of that of extant African colobines.

KNM-BN 1251 from Ngorora is a right P_4_ identified as distinct from that of KNM-BN 1740 (*Mi. tugenensis*) by Benefit and Pickford (1986) on the basis of discrete (i.e., presence of an accessory cuspule distal to the metaconid) and continuous (i.e., crown shape index of the P_4_) traits. The P_4_ KNM-BN 1251 is narrower and longer than that of KNM-BN 1740 (Benefit and Pickford, 1986), and when the samples from Ngerngerwa, Nakali, and Ngorora are combined, the coefficient of variation of the P_4_ crown shape index exceeds that of extant colobine species, although the magnitude of difference in coefficient of variation between extant and fossil sample is not as marked as that of the P_4_ area / M_1_ area index. However, the result from the bootstrap analyses of the P_4_ shape index is ambiguous. Although the combined sample of Ngerngerwa, Nakali and Ngorora sample exceeds the variation of *Pro. verus* and *Tr. cristatus*, it is close to the mean of the generated distribution of the coefficient of variation of *N. larvatus*. In addition, the morphology of the P_4_ cusps, that is their relative height and development, does not distinguish specimens from Nakali, Ngerngerwa, and Ngorora. Conclusively, and pending inclusion of additional fossil specimens, taxic diversity based on P_4_ crown dimensions is partially supported by the comparative assessment of coefficient of variation. Similarly, discrete dental traits, including relative heights and position of the cusps, does not firmly justify the presence of several species between Ngerngerwa, Nakali and Ngorora. Although previous work (Delson 1973; Swindler and Orlosky 1974) and our analysis have demonstrated qualitative and quantitative differences in the P_4_ morphology of extant colobines, further analytical work is needed to fully grasp its usefulness in taxonomic discrimination in the fossil record. This is especially true for the presence / absence of additional cuspules, and new advances in shape analysis, notably dental topography (Thiery et al. 2017, 2021; Plastiras et al. 2022), should provide more information about this.

As with most paleontological studies, the difficulty to provide firm conclusion regarding taxonomic distinction stems from a lack of data, making taxonomic assignments uncertain for small sample size. Nevertheless, we provide here a reassessment of the hypothesis of Benefit and Pickford (1986) and demonstrate that the taxonomic distinction between the Nakali and Ngerngerwa colobines is not substantiated by M_1_ morphology. Similarly, the differences in P_4_ crown dimensions are not of such magnitude as to allow us to draw reliable conclusions, hence mitigating the conclusions of Benefit and Pickford (1986). KNM-BN 1740 shares several dental and mandibular traits with Nakali specimens (e.g., overall size, symphyseal and corpus shape, length differential of the M_1_ and M_2_, and breadth differential of the M_2_ and M_3_ lophids). The reduced area of the P_4_ relative to that of the M_1_ is the only dental traits which, given the sample available, distinguish KNM-BN 1740 from the Nakali sample. The shape index value of KNM-BN 1740 is unusual and rarely seen in extant colobines (apart from outlying *N. larvatus* and *Pro. verus* specimens) nor in fossil colobines and *V. macinnesi*. However, this could be consistent with the range of variation of a paleospecies, as a similar range of variation is observed in the generated distribution of *N. larvatus*, *Pr. verus*, and *Co. guereza*. In conclusion, we have not observed continuous and discrete traits that justify a separate generic attribution between the Nakali and Ngerngerwa colobines, therefore we attribute the Nakali fossil colobines to *Microcolobus tugenensis*.

### H0_3_ The newly discovered Nakali specimens are distinct from the Eurasian and African Miocene colobines documented to date

The sample of fossil colobines is meager as only a handful of specimens are documented in the Chorora Formation (Beticha colobine), the Toros-Menalla fossiliferous area (*Ce. bruneti*), the Lukeino Formation (*S. lukeinoensis* and possibly another species represented by KNM-TH 36742), the Mpesida Beds (one species with indeterminate generic affinity), and Lothagam (two species with indeterminate generic affinity from the Nawata Formation and one species with indeterminate generic affinity from the Apak member of the Nachukui Formation). More complete collections are nonetheless known from Lemudong’o, where three species are documented (Hlusko, 2007). An even more complete sample of the fossil colobine *Mesopithecus* is provided by late Miocene sites from Eurasia, with *Me. delsoni* and *Me. pentelicus* (Delson 1973; De Bonis et al. 1990; Koufos et al. 2003).

*Microcolobus tugenensis* from Nakali is slightly smaller in M_1_ geometric size than the Beticha colobine CHO-BT 78, *S. lukeinoensis* OCO 607’10, the small colobine from Lemudong’o KNM-NK 36514, and the colobine sp. A from Nawata KNM-LT 24107. In M_2_ geometric size, it is also comparable to the isolated M_2_ KNM-TH 36742 from Lukeino. An increased dental sample could potentially expand the range of variation of the Nakali sample and encompass those comparative fossil specimens. However, based on size, it is unlikely that we sample, at Nakali, a species as large in M_1_ and M_2_ geometric size to that of the Mpesida colobine, the Nawata colobine sp. B, *Ce. bruneti*, *Pa. enkorikae*, *K. aramisi*, *Me. pentelicus*, *Me. delsoni*, and the indeterminate colobine species from Menacer.

The isolated M_2_ KNM-TH 36742 from Lukeino has lophid breadth proportions corresponding with the lower range of variation of the Nakali specimens. KNM-TH 36742 was recognized as ‘modern-looking’ partly based on its high occlusal relief. However, *Mi. tugenensis* KNM-NA 51103 also has a high occlusal relief, quantified by the NH / NR ratio, that falls in the range of variation of the extant colobines *Co. guereza*. We are aware of the limited resolution of our 3D data to quantifiy this trait in KNM-NA 51103 (Online Resource 12), but it is likely that it is underestimated compared to value obtained with finer technics. It is also intriguing to see that NH / NR ratio values of *Me. pentelicus* are overlapping with that of KNM-TH 36742 and KNM-NA 51103 given data from Benefit and Pickford, 1986. Pending a detailed analysis of the occlusal morphology and relief of the Nakali mandibles using dental topography, we recognized the colobine from Lukeino (KNM-TH 36742) as distinct from *Microcolobus tugenensis* from Nakali. However, we draw a cautious view regarding the identification of ‘modern-looking’ colobines in the fossil record given the overlap, in occlusal relief, of KNM-NA 51103 and extant colobines.

Two species of colobines were recognized in the Nawata Formation (Lothagam), including one (Colobinae gen. indet. sp. B KNM-LT 24098) that is much larger than *Microcolobus* in M_2_ geometric size. The second species, Colobinae gen. indet. sp. A KNM-LT 24107, is consistent in dental size with *Microcolobus* and its lophid breadth differential of the M_2_ are in the upper range of variation of the Nakali sample. Too little is known from this small fossil colobine to draw a conclusion, but future discoveries of a small-sized colobine in the Nawata Formation should benefit from a comparison with *Microcolobus*.

The relative elongation of the C_1_ of *Mi*. *tugenensis* is consistent with that of *K. aramisi*, *Pa. enkorikae*, *Me. pentelicus* and an isolated colobine C_1_ from Beticha. The shape of the P_4_ of *Mi*. *tugenensis* is compatible with most fossil colobines, except *Pa. enkorikae* and *Me. delsoni*. The area of the P_4_ relative to that of M_1_ in *Mi*. *tugenensis* is also consistent with that of most fossil colobines, except *K. aramisi*. In terms of the difference in intrinsic breadth differential of the molar lophids and relative length of M_1_ compared to that of M_2_, Miocene colobines are homogenous and without extensive samples, it is difficult to distinguish them based on those dental indices. The shape of the corpus of *Mi. tugenensis* is unlike *Pa. enkorikae*, and lacks enlarged lateral prominences but its shape is consistent with that of *Me. pentelicus*. The inferior transverse torus of *Mi. tugenensis* is not as pronounced as that of *Pa. enkorikae* and *Ce. bruneti*, but the low number of specimens preserving an intact symphysis and the overlap in the range of variation between the Nakali colobines and others Miocene fossil colobines prevent us from concluding. The inclined symphysis of *Mi.* tugenensis is unlike that of *Ce. bruneti*, *Me. pentelicus*, and *Me. delsoni*.

Qualitatively, the occlusal morphology of the P_4_ of *Mi. tugenensis* is consistent with that of *Pa. enkorikae* and *Ce. bruneti* but unlike that of *Me. pentelicus*. The metaconid is indeed less developed than the protoconid and placed more distally than that of *Me. pentelicus*. The M_3_ of *Mi*. *tugenensis* is broadly similar to that of others fossil colobines in exhibiting a moderately-developed hypoconulid devoid of C6. Qualitatively, the morphology of the corpus of *Mi*. *tugenensis* corresponds to the small Lemudong’o colobine KNM-NA 36514 by presenting weakly developed lateral prominences. All the *Me. delsoni* corpus included here (RZO 325 and RZO 326) are slightly distorted, but in cross-section at M_1_-M_2_ junction, they are much narrower than those of *Microcolobus*.

In conclusion, it is difficult to discuss the possible congenericity or conspecificity of the Nakali colobines with isolated teeth from Chorora, Lothagam, and Lukeino. The C_1_ from Beticha is described as lacking the elongated crown of extant Colobini, but this pattern is seen in all the African Miocene colobines studied here and is therefore not granted as taxonomically diagnostic. The intrinsic molar breadth differentials of the Beticha molars are consistent with that of *Microcolobus* (leaving aside the lack of resolution regarding the tooth loci of the Beticha molars) and are also consistent in size with it, hence supporting the presence of *Microcolobus* in Chorora. Likewise, there is no definitive evidence to rule out a close affinity between the small Lemudong’o colobine and *Microcolobus*, as they are similar in dental indices, in P_4_ and M_3_ occlusal morphology, and corpus shape. It is also complicated to distinguish *Microcolobus* from *Sawecolobus* based on our dental indices and the corpus and symphysis of *Sawecolobus* is too damaged to provide a solid comparative basis. The smaller P_4_ metaconid relative to protoconid of *Sawecolobus* is nevertheless consistent with *Microcolobus*. The increased sample of upper teeth and maxillae from *Sawecolobus* should provide a better basis of comparison, waiting for the description of the upper dentition and maxillae of *Microcolobus* from Nakali.

### H0_4_ The newly discovered Nakali specimens are distinct from extant African colobines (Colobini)

Given its relatively long thumb, the partial skeleton KNM-NA 48915, assigned to *Microcolobus*, was hypothesized to be a stem Colobinae. Its affinity with either the African (Colobini) or Asian (Presbytini) tribes remains to be confirmed based on dental features.

Our analysis provided ambiguous results regarding the phylogenetic affinity of *Microcolobus* as it shows a mosaic of dental features shared either with Colobini or Presbytini. Indeed, the lower canine of male *Microcolobus* is not as elongated as that of extant Colobini. Its P_4_ occlusal morphology is nevertheless consistent with the anatomy of African colobines, and our linear discriminant analysis, based on the dimensions of the premolars and molars (excepting M_3_ length) classifies *Microcolobus* among the African colobines with a high posterior probability (>80%). The symphysis of *Microcolobus*, in its relative proportion of the transverse tori and its symphyseal inclination, is not incompatible with the symphyseal anatomy of *Procolobus* and *Piliocolobus*. However, it is distinguished from African colobines by having weakly-developed lateral prominences.

Phenetically, therefore, it is likely that *Microcolobus* is a stem Colobinae. However, affinity with the stem branch of the Colobini is a hypothesis that should be considered in future studies, with the later acquisition in crown Colobini of an elongated C_1_ and enlarged lateral prominences compared to *Microcolobus*. Future phylogenetic analyses revealing dental traits polarity in colobines will be of great value. In the following section, we emphasized the comparison of *Microcolobus* with *Victoriapithecus* and *Noropithecus* to discuss potential trait polarity and morphological novelty of the mandibular anatomy of early colobines compared to the earliest cercopithecoids.

### Implication on the evolution of Miocene colobines

*Victoriapithecus* and *Noropithecus* are Miocene victoriapithecids lacking dental features seen in crown cercopithecids, such as the loss of hypoconulid on M_1_ and M_2_ and completion of bilophodonty on upper molars (Benefit 1993, 1999; Miller et al. 2009; Gilbert et al. 2010; Locke et al. 2020). In four dental indices, *V. macinnesi* exhibits a range of variation inconsistent with that of extant colobines. It shows a more equal ratio of the P_4_ area relative to the M_1_ area, a marked M_1_ -M_2_ length differential in favor of the M_2_, and a distinct breadth differential of the M_2_ and M_3_ lophids, with the mesial lophid broader than the distal lophid. In all these dental indices, *Microcolobus* deviates from the victoriapithecid condition.

It is less obvious to differentiate victoriapithecids from *Microcolobus* based on corpus and symphyseal shape. While the symphysis of *V. macinnesi* is acutely angled and presents a diminutive inferior transverse torus relative to the superior transverse torus, the symphysis of *N. bulukensis* is extremely similar to that of *Microcolobus* and *Me. delsoni* and is distinct from *V. macinnesi* by its well-developed inferior transverse torus and its less acutely angled symphysis. Phenetically, this demonstrates the presence of a colobine-like and a cercopithecine-like symphysis in victoriapithecids. While the symphysis of *V. macinnesi* is different from that of *Microcolobus*, the shape of its corpus cross-section, as seen for example in KNM-MB 27876, is fully consistent with that of *Microcolobus* in exhibiting a distal thinning of the corpus and weakly expressed lateral prominences. The same is true for *N. bulukensis*, which has a slightly less inflated corpus than *V. macinnesi* but is still similar in shape to *Microcolobus*.

The dental anatomy of *Noropithecus* and *Victoriapithecus* is distinct from that of crown cercopithecids and, as our study and others have demonstrated, can be metrically distinguished from extant cercopithecid subfamilies (Benefit 1993, 2008; Miller et al. 2009; Gilbert et al. 2010; Locke et al. 2020). It is more complicated to distinguish victoriapithecids and stem representatives of the crown cercopithecids using the shape of the corpus and symphysis. As already suggested (Benefit, 2008; Miller et al. 2009; Locke et al. 2020), and demonstrated independently here, the symphysis of *Noropithecus* shows features typical of extant colobines (e.g., broad inferior transverse torus, except in *Colobus*) and that of *Victoriapithecus* typical of extant cercopithecines (e.g., narrow inferior transverse torus in *Cercopithecus*). Here we demonstrate that *Microcolobus*, the oldest fossil colobine to preserve the mandibular anatomy, is consistent in corpus and symphyseal shape to *N. bulukensis*. Benefit and Pickford (1986) and Locke et al. (2020) raised the possibility that fossil taxa of extant cercopithecid subfamilies could be represented in the victoriapithecid fossil record based on dental and symphyseal traits, respectively, but then disregarded this hypothesis citing the lack of parsimony that would implicate a parallel dental evolution between cercopithecid subfamilies (e.g., complete bilophodonty and loss of M_1_ and M_2_ hypoconulids). Although we endorse this view, we cannot dismiss the working hypothesis of the presence of a stem colobine and/or a stem cercopithecine in the probably paraphyletic assemblage of victoriapithecids (Miller et al. 2009; Rasmussen et al. 2019). Identification of traits polarity of the considered symphyseal and corpus traits would be decisive in this issue.

### New insights in dental differences between extant and fossil colobines

Based on the comparison of the dental indices of the three extant Colobini genera (*Procolobus*, *Piliocolobus*, and *Colobus*) with three Presbytini genera (*Nasalis*, *Presbytis*, and *Trachypithecus*), we demonstrated the presence of significant differences between subtribes in intrinsic breadth of the M_1_, M_2_, and M_3_ lophids, as well as in P_4_ shape and relative length of the C_1_. These results have implications for previous assessments of the fossil record (Gilbert et al. 2010; Gommery et al. 2022). Most notable is the use of the breadth differential of the M_3_ lophids in distinguishing Colobini from Presbytini in the fossil record (Gilbert et al. 2010; Gommery et al. 2022). There is indeed a significant difference between the two subtribes, but subtler interpretation of the data is needed to fully grasp the usefulness of this trait. In fact, this difference is mainly due to the reduced breadth of the M_3_ distal lophid of *Pre. chrysomelas* rather than by an enlargement of the distal lophid in Colobini, which are indistinguishable from *N. larvatus*, and apart from *Co. guereza*, indistinguishable from *Tr. cristatus*. The same is true for the M_1_ lophid differential, which is mainly driven by the subequal lophids of *Tr. cristatus*.

Stronger statistical evidence regarding the Colobini - Presbytini distinction is borne out from the P_4_ shape and relative length of the C_1_ indices, where each Colobini genus can be differentiated from that of Presbytini. Likewise, standard linear dimensions of the premolars and molars (except M_3_ length), when included into a linear discriminant analysis via principal component scores, can effectively discriminate each subtribe. Overall, our morphometric analyses will benefit from the inclusion of additional Presbytini genera (*Pygathrix*, *Rhinopithecus*, and *Semnopithecus*) to fully conclude on the usefulness of dental indices to distinguish extant subtribes and to justify the inclusion of fossil specimens, on phenetic bases, in these subtribes.

## CONCLUSION

The description and morphometric analyses of six new mandibles from the Miocene site of Nakali (Kenya) provide new data on the dental, symphyseal, and corpus anatomy of *Microcolobus*. The sample of fossil mandibles from Nakali is morphologically homogenous and corresponds to a single species, *Microcolobus tugenensis*. Isolated dental elements, the P_4_ KNM-BN 1251 and the M_1_ KNM-NA 305, previously recognized as distinct from *Mi. tugenensis*, are here recognized as part of a single paleospecies, *Mi. tugenensis*. A linear discriminant analysis established on a principal component analysis of dental dimensions classified four of the most complete *Microcolobus* specimens as belonging to the African colobine tribe (Colobini). The symphyseal anatomy and the occlusal morphology of the P_4_ of *Microcolobus* is also similar to extant African colobines. However, the shape of the corpus of *Microcolobus* and its lower canine dimensions does not provide evidence for a close affinity with Colobini, but rather indicate, on a phenetic basis, a stem Colobinae status for *Microcolobus*.

## Supporting information

Supplementary Informations and Table 3

## ACKNOWLEDGEMENTS

The authors thank the curators and staff of the National Museums of Kenya (Osteology Division, Nairobi), the Royal Museum for Central Africa (Dr. Emmanuel Gilissen and Mathys Rotonda from the Division of Mammals, Tervuren), the Center for the Evolutionary Origins of Human Behavior (Assoc. Pro. Takeshi Nishimura the Division of Mammals), the US National Museum of Natural History (D. Schlitter, Division of Mammals), and the Bavarian State Collection for Zoology (Dr. Anneke van Heteren and Michael Hiermeier) for allowing them to study of skeletal collections under their care. We also thank the NSF BCS 1552848 attributed to Dr. D. Boyer, and Assoc. Prof. Claire Terhune (University of Arkansas) for data sharing via the MorphoSource platform. We are grateful to the Domestic Research Program of Ryokoku University. This research was financially supported by the Japan Society for the Promotion of Science JSPS Kakenhi 25257.

