## Supplementary Informations and Table 3 for "Description of new mandibular remains of *Microcolobus* from Nakali (ca. 10 Ma, Kenya): implications on the evolution of Miocene colobines": ESM_9_12_13_14_15_17.pdf

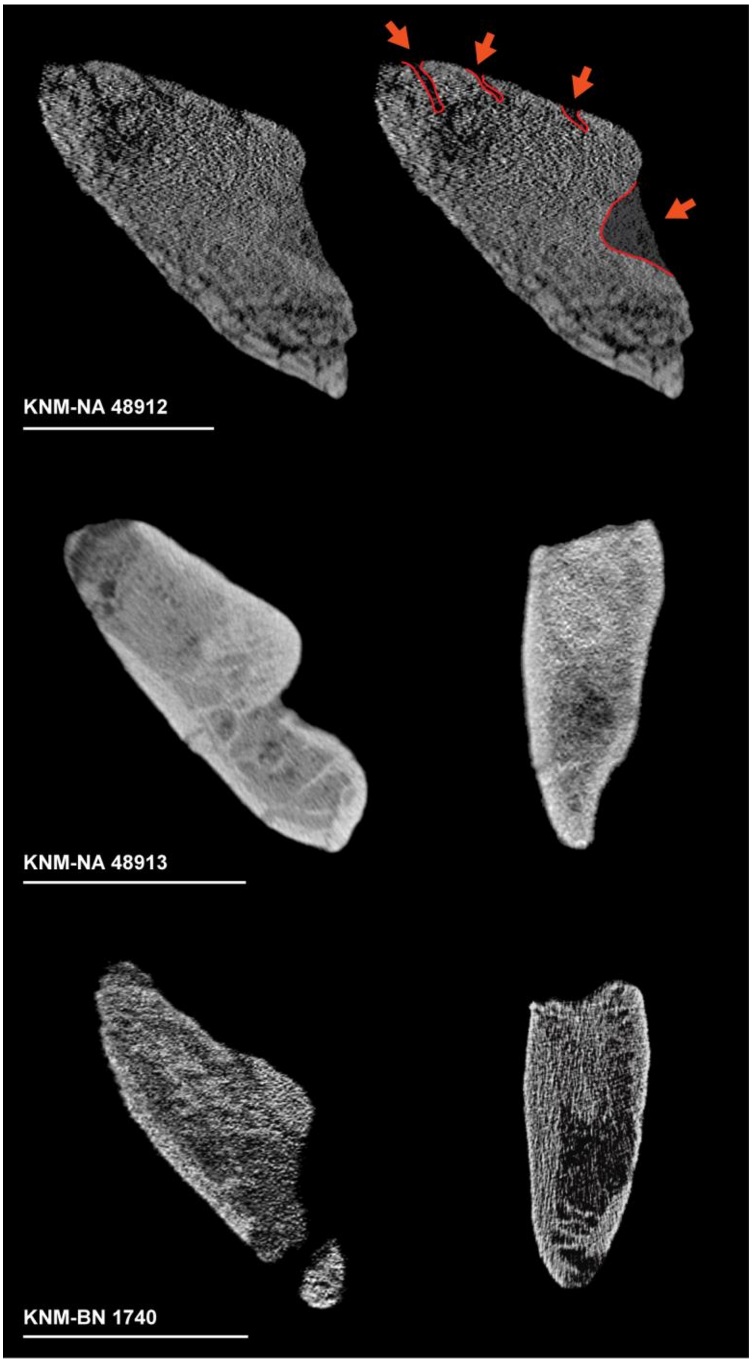


Online Resource 9: Transverse cross-sections of the symphysis of KNM-NA 48912, KNM-NA 48913, and KNM-BN 1740 along with corpus cross-section at $M_1$-$M_2$ junction obtained using a PQ-CT. Red arrows denote the deformation of the superior surface of the planum alveolare of KNM-NA 48912 and matrix infilling of its genioglossal fossa. Note the damaged inferior transverse torus of *Microcolobus tugenensis* KNM-BN 1740.

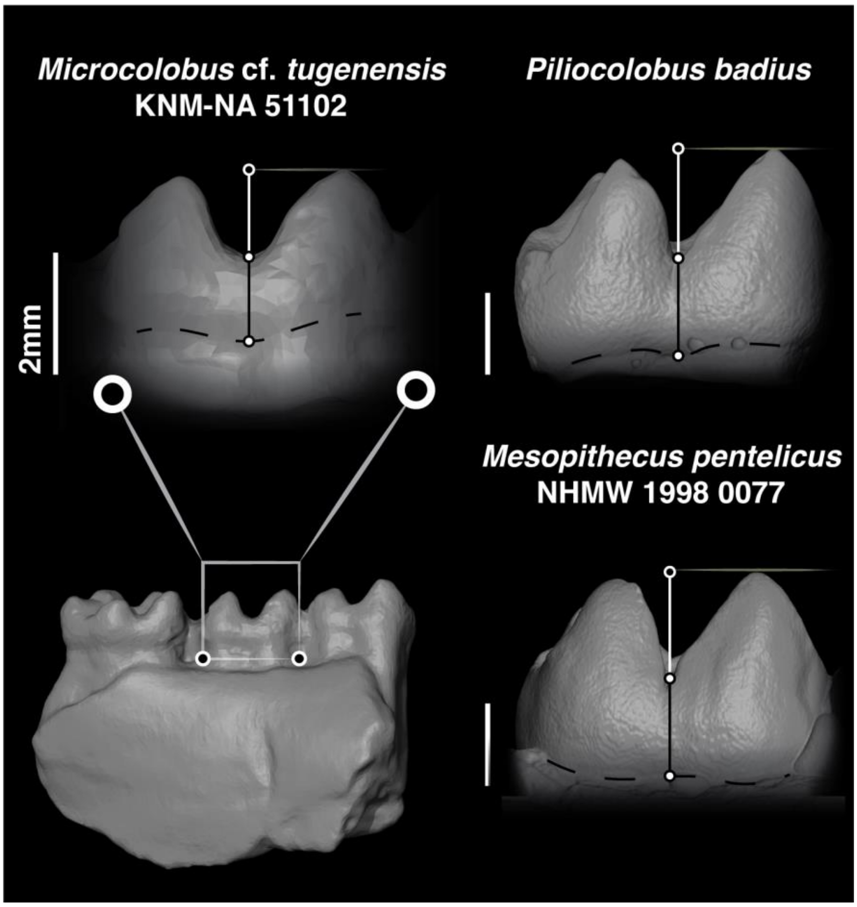


Online Resource 12: Dental occlusal relief of the $M_2$ KNM-NA 51102 in comparison to that of *Piliocolobus badius*. The black line represents the height of the crown below the notch (NR *sensu* Benefit and Pickford, 1986) and the white line represent the height of the cusp above the notch (NH *sensu* Benefit and Pickford, 1986).

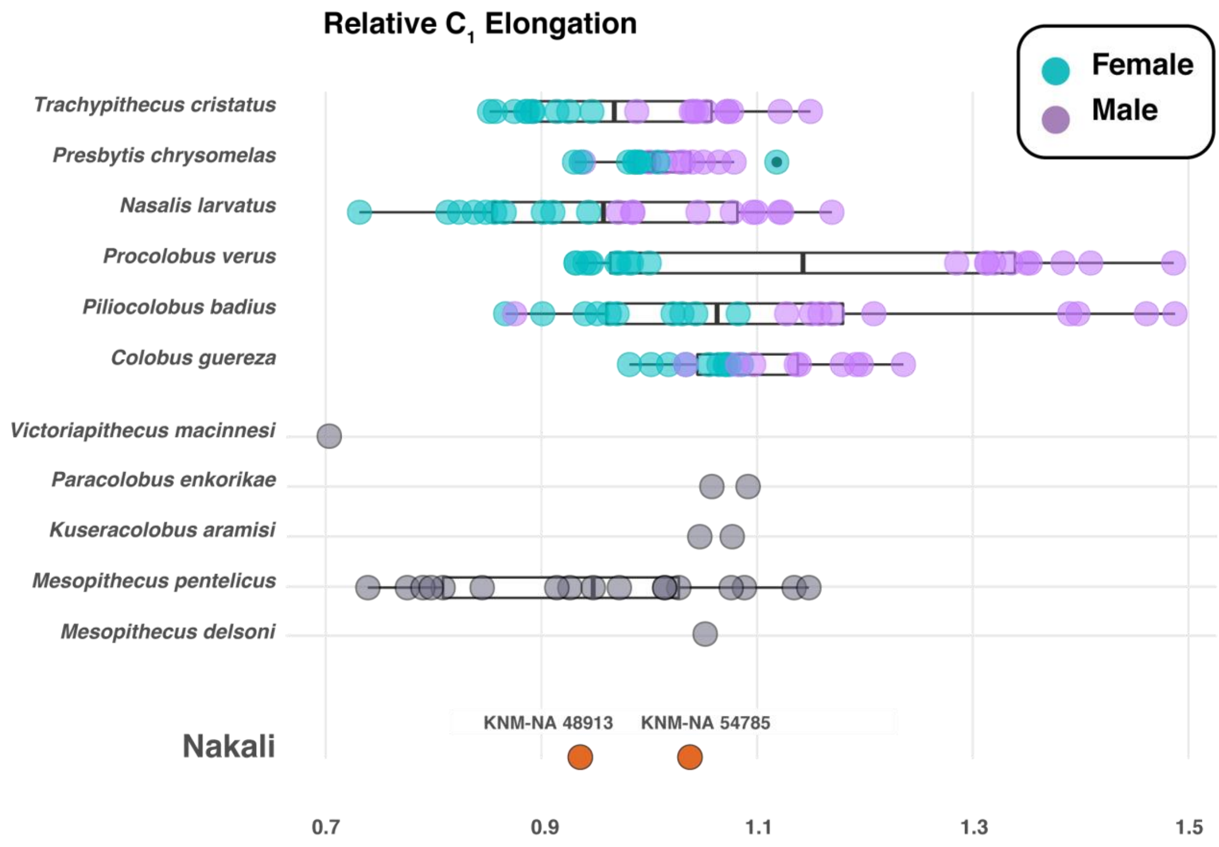


Online Resource 13: Boxplots of the index of the relative elongation of the $C_1$ of extant and fossil colobines. Boxplots with first, third quartile, and median (black line). Note the highly elongated C1 of male *Pi. badius* and *Pro. verus*.

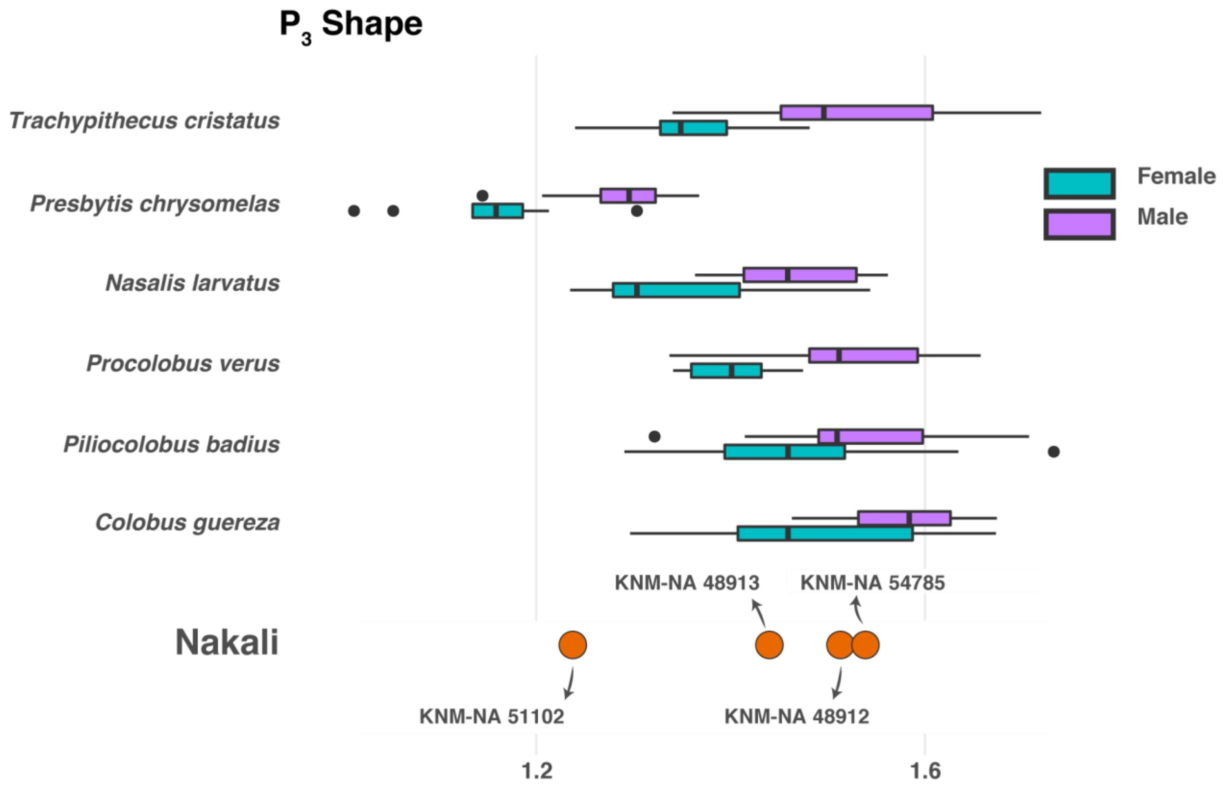


Online Resource 14: Boxplots of the $P_3$ shape index of extant and fossil colobines. Boxplots with first, third quartile, and median (black line). Note the marked dimophism and absence of overlap of interquartile range in *Tr. cristatus*, *Pre. chrysomelas*, *N. larvatus*, and *Pro. verus*.

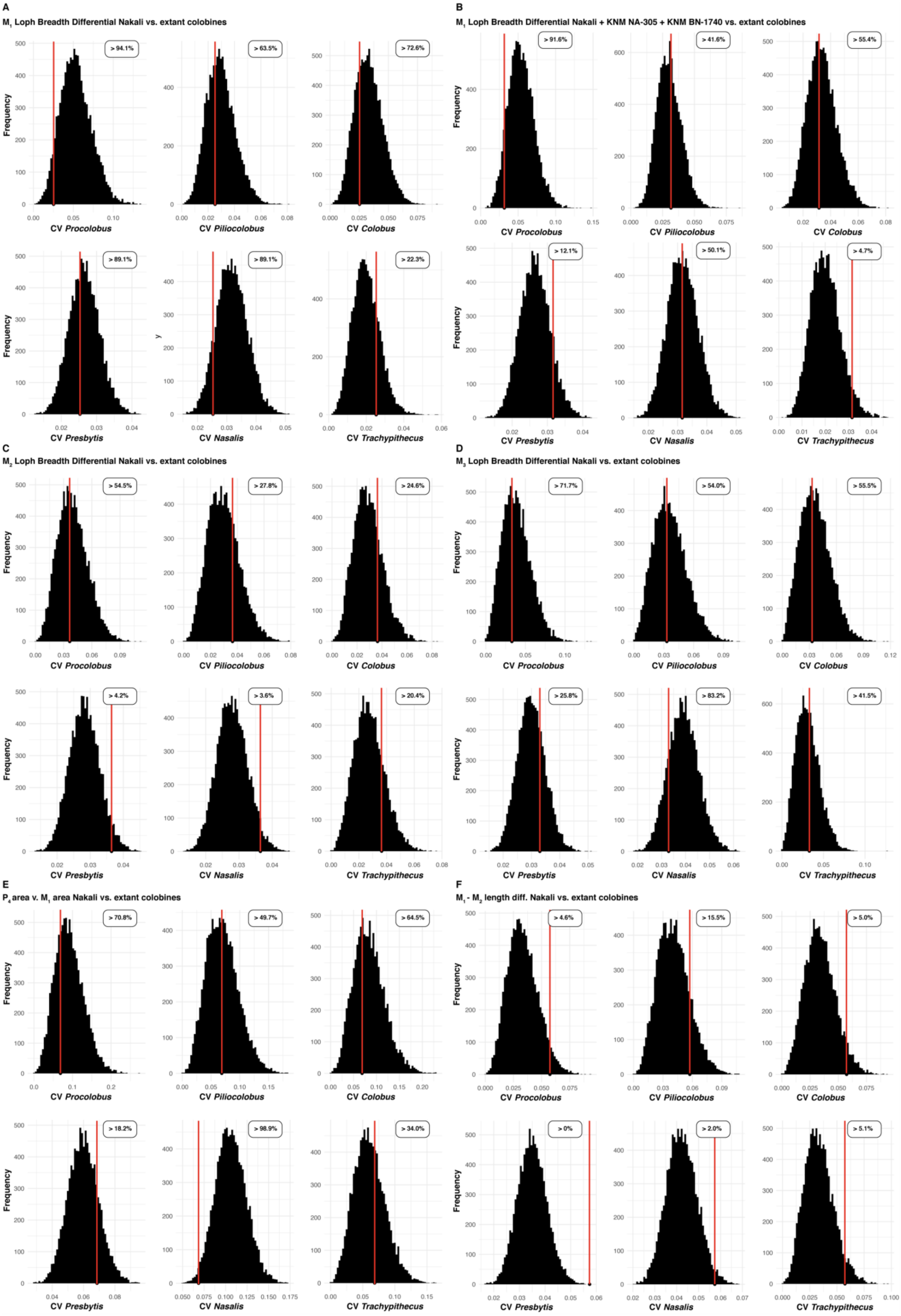
A
$M_1$ Loph Breadth Differential Nakali vs. extant colobines
> 94.1%
> 63.5%
> 72.6%
> 89.1%
> 89.1%
> 22.3%
CV Procolobus
CV Piliocolobus
CV Colobus
CV Presbytis
CV Nasalis
CV Trachypithecus
Frequency
B
$M_1$ Loph Breadth Differential Nakali + KNM NA-305 + KNM BN-1740 vs. extant colobines
> 91.6%
> 41.6%
> 55.4%
> 12.1%
> 50.1%
> 4.7%
C
$M_2$ Loph Breadth Differential Nakali vs. extant colobines
> 54.5%
> 27.8%
> 24.6%
> 4.2%
> 3.6%
> 20.4%
D
$M_3$ Loph Breadth Differential Nakali vs. extant colobines
> 71.7%
> 54.0%
> 55.5%
> 25.8%
> 83.2%
> 41.5%
E
$P_4$ area v. $M_1$ area Nakali vs. extant colobines
> 70.8%
> 49.7%
> 64.5%
> 18.2%
> 98.9%
> 34.0%
F
$M_1$ - $M_2$ length diff. Nakali vs. extant colobines
> 4.6%
> 15.5%
> 5.0%
> 0%
> 2.0%
> 5.1%

Online Resource 15 (previous page): Frequency distribution of the coefficient of variation of the generated distributions of extant colobine species for the $M_1$ lophids breadth differential (A and B), $M_2$ lophids breadth differential (C), $M_3$ lophids breadth differential (D), $P_4$ area relative to $M_1$ area (E), and $M_1$ - $M_2$ length differential. Probability of observing a coefficient of variation higher than that of the considered fossil sample is shown on the upper right of each frequency distribution. The observed coefficient of variation for the considered fossil sample is indicated by a red vertical line.

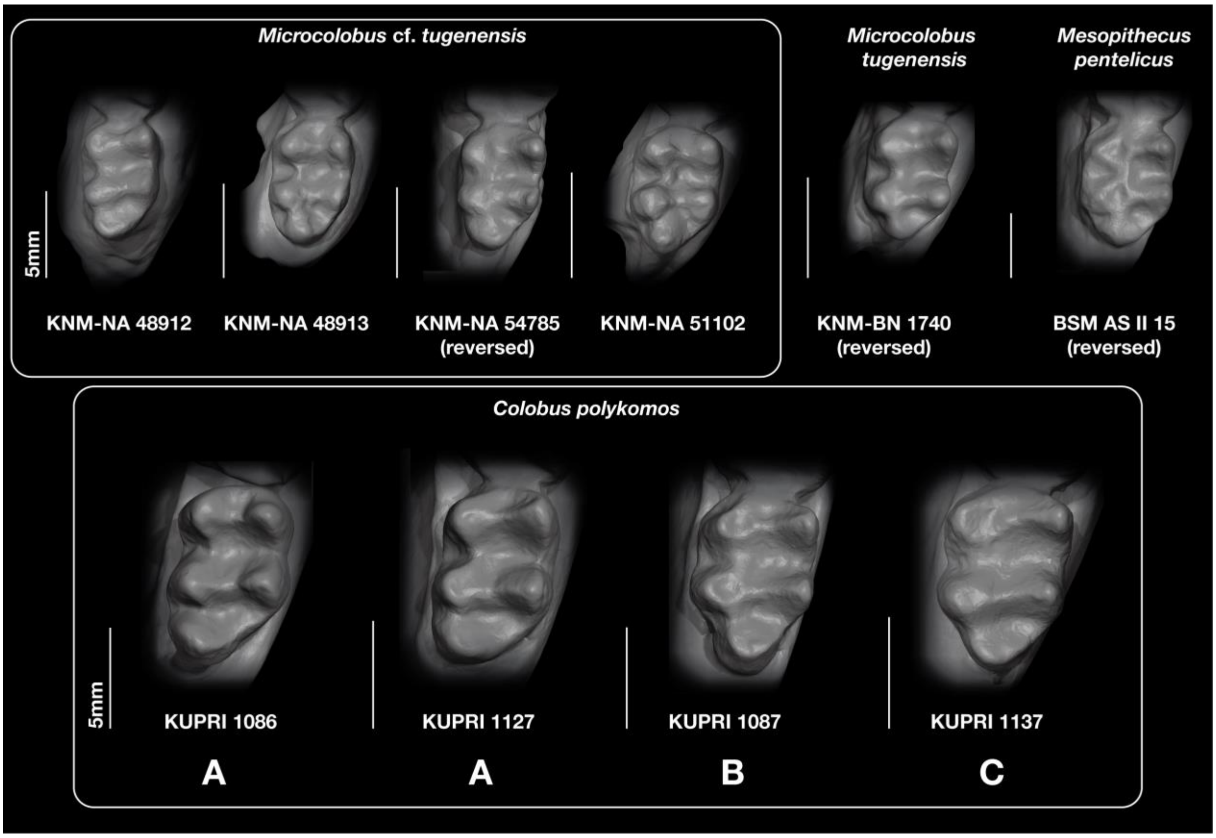


Online Resource 17: Comparison of the $M_3$ morphology of fossil colobines and a selected set of *Colobus polykomos* specimens to highlight differences in hypoconulid development. *Colobus* specimens of the A category illustrates the most common morphology of hypoconulid (i.e., buccally-placed and moderately-developed), the specimen of the category B illustrates moderately-developed and centrally-placed hypoconulid, and that of category C illustrates small hypoconulid centrally-placed. Note the variation in hypoconulid placement and development in the Nakali sample and the similariy in hypoconulid shape between *Me. pentelicus* BSM AS II 15, *Mi. tugenensis* KNM-BN 1740, *Mi.* cf. *tugenensis* KNM-NA 51102, and *Co. polykomos* KUPRI 1137.
