## Supplementary Informations and Table 3 for "Description of new mandibular remains of *Microcolobus* from Nakali (ca. 10 Ma, Kenya): implications on the evolution of Miocene colobines": ESM_11_and_16.docx

Online Resource 11: Results of the normality and homoscedasticity tests for the dental indices

| Formula | Normality (Shapiro-Wilk) | Homoscedasticity (Bartlett) |
| --- | --- | --- |
| M1 loph. diff. ~ Tribes | W = 1  *p*-value = 0.1 | K^2^ = 0.3  *p*-value = 0.6 |
| M1 loph. diff. ~ Genera | W = 1  *p*-value = 0.002 | K^2^ = 21  *p*-value < 0.001 |
| M2 loph. diff. ~ Tribes | W = 1  *p*-value = 0.4 | K^2^ = 2  *p*-value = 0.2 |
| M2 loph. diff. ~ Genera | W = 1  *p*-value = 0.4 | K^2^ = 4  *p*-value = 0.5 |
| M3 loph. diff. ~ Tribes | W = 1  *p*-value = 0.03 | K^2^ = 0.01  *p*-value = 0.9 |
| M3 loph. diff. ~ Genera | W = 1  *p*-value = 0.05 | K^2^ = 6  *p*-value = 0.3 |
| P4 ar. M1 ar. ~ Tribes | W = 1  *p*-value = 0.4 | K^2^ = 2  *p*-value = 0.2 |
| P4 ar. M1 ar. ~ Genera | W = 1  *p*-value = 0.7 | K^2^ = 7  *p*-value = 0.2 |
| M1 lg. M2 lg. ~ Tribes | W = 1  *p*-value = 0.4 | K^2^ = 5  *p*-value = 0.02 |
| M1 lg. M2 lg. ~ Genera | W = 1  *p*-value = 0.7 | K^2^ = 2  *p*-value = 0.9 |
| P4 shape ~ Tribes | W = 1  *p*-value = 0.01 | K^2^ = 30  *p*-value < 0.001 |
| P4 shape ~ Genera | W = 1  *p*-value = 0.5 | K^2^ = 24  *p*-value < 0.001 |
| C1 Rel. ~ Tribes | W = 1  *p*-value = 0.002 | K^2^ = 13  *p*-value < 0.001 |
| C1 Rel. ~ Genera | W = 1  *p*-value = 0.03 | K^2^ = 43  *p*-value < 0.001 |

Online Resource 16: Significance of the parametric and non-parametric tests regarding differences in dental indices between extant colobines

| **Formula** | **Test** | ***p*-value and test parameters** | |
| --- | --- | --- | --- |
| M1 loph. diff. ~ Tribes | ANOVA | F = 5.21  *p*-value = 0.024 | |
| M1 loph. diff. ~ Genera | Dunn (post-hoc) | *Pre. chrysomelas* - *N. larvatus* | Z = -3.391, *p*-value = 0.008 |
|  |  | *Tr. cristatus* - *Co. guereza* | Z = -3.391, *p*-value = 0.02 |
|  |  | *Tr. cristatus* - *N. larvatus* | Z = -5.534, *p*-value < 0.001 |
|  |  | *Tr. cristatus* - *Pi. badius* | Z = -4.993, *p*-value < 0.001 |
|  |  | *Tr. cristatus* - *Pro. verus* | Z = -4.067, *p*-value < 0.001 |
| M2 loph. diff. ~ Tribes | ANOVA | F = 29.6  *p*-value < 0.001 | |
| M2 loph. diff. ~ Genera | Tukey HSD | *Pre. chrysomelas* - *N. larvatus* | *p*-value = 0.02 |
|  |  | *Pre. chrysomelas* - *Pi. badius* | *p*-value = 0.001 |
|  |  | *Pre. chrysomelas* - *Pro. verus* | *p*-value = 0.002 |
|  |  | *Tr. cristatus* - *Pro. verus* | *p*-value = 0.03 |
| M3 loph. diff. ~ Tribes | ANOVA | F = 19.5  *p*-value < 0.001 | |
| M3 loph. diff. ~ Genera | Tukey HSD | *Pre. chrysomelas* - *Co. guereza* | *p*-value < 0.001 |
|  |  | *Pre. chrysomelas* - *N. larvatus* | *p*-value < 0.001 |
|  |  | *Pre. chrysomelas* - *Pi. badius* | *p*-value < 0.001 |
|  |  | *Pre. chrysomelas* - *Pro. verus* | *p*-value < 0.001 |
|  |  | *Tr. cristatus* - *Co. guereza* | *p*-value = 0.018 |
|  |  | *Tr. cristatus* - *Pre. chrysomelas* | *p*-value = 0.001 |
| P4 Ar. M1. Ar. ~ Tribes | ANOVA | F = 0.27  *p*-value = 0.61 | |
| P4 Ar. M1. Ar. ~ Genera | Tukey HSD | *Co. guereza* - *N. larvatus* | *p*-value < 0.001 |
|  |  | *Co. guereza* - *Pi. badius* | *p*-value < 0.001 |
|  |  | *Co. guereza* - *Pro. verus* | *p*-value < 0.001 |
|  |  | *N. larvatus* - *Pre. chrysomelas* | *p*-value = 0.005 |
|  |  | *N. larvatus* - *Tr. cristatus* | *p*-value < 0.001 |
|  |  | *Tr. cristatus - Pi. badius* | *p*-value = 0.002 |
|  |  | *Tr. cristatus - Pro. verus* | *p*-value < 0.001 |
|  |  | *Pre. chrysomelas - Pro. verus* | *p*-value = 0.010 |
| M1. lg. M2. lg. ~ Tribes | Kruskal-Wallis | X^2^ =3, *p*-value = 0.09 | |
| M1. lg. M2. lg. ~ Genera | Tukey HSD | *Pre. chrysomelas* - *Co. guereza* | *p*-value = 0.010 |
|  |  | *Pre. chrysomelas* - *Pi. badius* | *p*-value = 0.025 |
|  |  | *Pre. chrysomelas* - *Pro. verus* | *p*-value = 0.015 |
|  |  | *N. larvatus* - *Pi. badius* | *p*-value = 0.002 |
|  |  | *N. larvatus* - *Pre. chrysomelas* | *p*-value < 0.001 |
|  |  | *N. larvatus* - *Pro. verus* | *p*-value = 0.003 |
|  |  | *Tr. cristatus* - *Pi. badius* | *p*-value = 0.036 |
|  |  | *Tr. cristatus* - *Pro. verus* | *p*-value < 0.001 |
| P4 Shp ~ Tribes | Kruskal-Wallis | X^2^ =80, *p*-value < 0.001 | |
| P4 Shp ~ Genera | Dunn (post-hoc) | *N. larvatus* - *Co. guereza* | Z = -3.656, *p*-value = 0.002 |
|  |  | *N. larvatus* - *Pi. badius* | Z = 3.125, *p*-value = 0.01 |
|  |  | *N. larvatus* - *Pro. verus* | Z = 3.989, *p*-value < 0.001 |
|  |  | *N. larvatus* - *Pre. chrysomelas* | Z = -3.107, *p*-value = 0.01 |
|  |  | *Pre. chrysomelas* - *Co. guereza* | Z = -6.665, *p*-value < 0.001 |
|  |  | *Pre. chrysomelas* - *Pi. badius* | Z = -6.148, *p*-value < 0.001 |
|  |  | *Pre. chrysomelas* - *Pro. verus* | Z = 6.989, *p*-value < 0.001 |
|  |  | *Tr. cristatus* - *Co. guereza* | Z = -4.326, *p*-value < 0.001 |
|  |  | *Tr. cristatus* - *Pi. badius* | Z = -3.795, *p*-value = 0.001 |
|  |  | *Tr. cristatus* - *Pro. verus* | Z = -4.659, *p*-value < 0.001 |
| C1 Rel ~ Tribes | Kruskal-Wallis | X^2^ =13, *p*-value < 0.001 | |
| C1 Rel ~ Genera | Dunn (post-hoc) | *Co. guereza* - *N*. *larvatus* | Z = 3.221, *p*-value = 0.020 |
|  |  | *Co. guereza* - *Pre. chrysomelas* | Z = 4.049, *p*-value < 0.001 |
|  |  | *Co. guereza* - *Tr. cristatus* | Z = 2.915, *p*-value = 0.040 |
|  |  | *Pi. badius* - *Pre. chrysomelas* | Z = 3.111, *p*-value = 0.022 |
|  |  | *Pro. verus* - *N. larvatus* | Z = -2.934, *p*-value = 0.040 |
|  |  | *Pro. verus* - *Pre. chrysomelas* | Z = -3.782, *p*-value = 0.002 |
